# Towards an unbiased characterization of genetic polymorphism: a comparison of 27 *A. thaliana* genomes

**DOI:** 10.1101/2024.05.30.596703

**Authors:** Anna A. Igolkina, Sebastian Vorbrugg, Fernando A. Rabanal, Hai-Jun Liu, Haim Ashkenazy, Aleksandra E. Kornienko, Joffrey Fitz, Max Collenberg, Christian Kubica, Almudena Mollá Morales, Benjamin Jaegle, Travis Wrightsman, Vitaly Voloshin, Alexander D. Bezlepsky, Victor Llaca, Viktoria Nizhynska, Ilka Reichardt, Christa Lanz, Felix Bemm, Pádraic J. Flood, Sileshi Nemomissa, Angela Hancock, Ya-Long Guo, Paul Kersey, Detlef Weigel, Magnus Nordborg

## Abstract

Our view of genetic polymorphism is shaped by methods that provide a limited and reference-biased picture. Long-read sequencing technologies, which are starting to provide nearly complete genome sequences for population samples, should solve the problem—except that characterizing and making sense of non-SNP variation is difficult even with perfect sequence data. Here we analyze 27 genomes of *Arabidopsis thaliana* in an attempt to address these issues, and illustrate what can be learned by analyzing whole-genome polymorphism data in an unbiased manner. Estimated genome sizes range from 135 to 155 Mb, with differences almost entirely due to centromeric and rDNA repeats that are difficult to assemble. The completely assembled chromosome arms comprise roughly 120 Mb in all accessions, but are full of structural variants, largely due to transposable elements. Even with only 27 accessions, a pan-genome coordinate system that includes the resulting variation ends up being ∼ 70% larger than the size of any one genome. Our analysis reveals an incompletely annotated mobile-ome: we not only detect several novel TE families, but also find that existing TE annotation is a poor predictor of elements that have recently been active. In contrast to this, the genic portion, or “gene-ome”, is highly conserved. By annotating each genome using accession-specific transcriptome data, we find that 13% of all (non-TE) genes are segregating in our 27 accessions, but most of these are transcriptionally silenced. Finally, we show that with short-read data we previously massively underestimated genetic variation of all kinds, including SNPs—mostly in regions where short reads could not be mapped reliably, but also where reads were mapped incorrectly. We demonstrate that SNP-calling errors can be biased by the choice of reference genome, and that RNA-seq and BS-seq results can be strongly affected by mapping reads only to a reference genome rather than to the genome of the assayed individual. In conclusion, while whole-genome polymorphism data pose tremendous analytical challenges, they also have the potential to revolutionize our understanding of genome evolution.

## Introduction

The last 25 years have witnessed an explosion of genetic polymorphism data, fueled by Human Genome Project-inspired collaborations and the development of massively parallel technologies for sequencing and genotyping. Such data allow us to study population history, selection, and genetic architecture of traits—as well as the evolution of the genome itself. However, our current view of genetic polymorphism has been substantially shaped by technologies that attempt to align short sequence fragments to a reference genome in order to detect sites that differ. As a result, our knowledge of genome variation has remained incomplete and biased towards simple variants in regions that are easy to align— a small fraction of the genome in many species. A further source of bias arises from the use of a single reference genome.

All this is beginning to change now that long reads are making it possible to assemble high-quality, full-length chromosomal sequences from population samples. During the last couple of years, nearly complete genomes have been produced for large numbers of eukaryotic species, including yeast, animals (including fruit flies, humans, and cichlids), and many plants, including rice, tomato, soybean, grapevine, wheat, barley, maize, millet, *Brassica oleracea, Eucalyptus, Populus* and *Marchantia* sp.—as well as *A. thaliana* ^1–27^. Impressive as these studies are, they have also highlighted how difficult it is to make sense of whole-genome polymorphism data, primarily because sequence alignment is not unambiguous. Pan-genome graphs ^28,29^ may provide elegant and computationally efficient ways of representing such data, but they do not solve this fundamental problem. To compare genomes and interpret their differences properly, we require a modeling framework that reflects the mutational mechanisms and recombination history that gave rise to these differences, but such a framework is still largely missing.

Here we illustrate this problem by analyzing the genomes of 27 natural inbred accessions of *A. thaliana*, chosen to cover the global genetic diversity of the species. Our focus is on obtaining an unbiased picture of polymorphism in the more easily alignable chromosome arms and comparing it to existing data built over almost two decades ^30–34^. To provide an unbiased picture of the “gene-ome”, *i*.*e*., the collection of genes across multiple genomes, we complement our genome assemblies with transcriptomes from multiple tissues for the entire sample. In addition, we seek to lay the foundation for a community resource that will eventually comprise complete genomes for the thousands of natural inbred accessions that are publicly available for this model plant, thus connecting whole-genome polymorphism data to experimentally accessible germplasm and knowledge of a wide range of morphological, life-history, physiological and molecular phenotypes, as well as precise collection information (see 1001genomes.org).

## Results

### The organization of genome variation

We selected 27 accessions to cover global genetic variation of the species based on the original 1001 Genomes project ^34^, and additional samples from eastern Asia ^35^, Africa ^36^, and Madeira ^37^ (Extended Data Fig. 1). We sequenced their genomes with PacBio continuous long reads (CLRs) to high depth, assembled them into contigs with Canu ^38^, and polished these contigs with Illumina PCR-free short reads ^39^ (for assembly statistics, see Supplementary Table 1). To reconstruct chromosomes, we generated hybrid scaffolds with individual Bionano optical maps for eight accessions, which allowed us to determine the most appropriate parameters for scaffolding ^40^ based on the TAIR10 reference genome ^41^. Although this potentially introduces some reference bias, our focus is on the gene-dense chromosome arms, which are well-covered by large contigs, and appear to harbor few large-scale rearrangements.

Like many plants, *A. thaliana* has experienced recent episodes of TE activity, leading to nearly identical sequences inserted across each genome ^42^. These make short-read alignments difficult, but the PacBio CLR technology used here produced reads long enough to bridge such insertions. However, extensive tracts of identical or near-identical tandem repeats, such as centromere satellites and 5S rDNAs, consistently break our assemblies (Extended Data Figs. 2-3 We note that centromeres can now be assembled with PacBio HiFi reads ^16^. CLRs are less accurate, but they are on average longer, hence marginally better for chromosome arms ^43^. 45S rDNA clusters remain challenging regardless of technology ^44^.

Our 27 assemblies are all ∼ 120 Mb in size, whereas the full genomes, consistent with previous results ^45,46^, are estimated to range from 135 to 155 Mb (Fig. 1A). A BLAST-based approach indicates that centromeres and sDNA clusters alone account for up to 92% of the estimated variation (Fig. 1B-C), with the importance of 45S rDNA variation having been appreciated before ^46^. While individual TE families can vary greatly in size across accessions (Extended Data Fig. 4), we confirm that the cumulative effect of all TEs on genome size variation appears to be small in *A. thaliana* ^46^—contrary to the major role they play in inter-specific variation ^47,48^.

**Figure 1.**
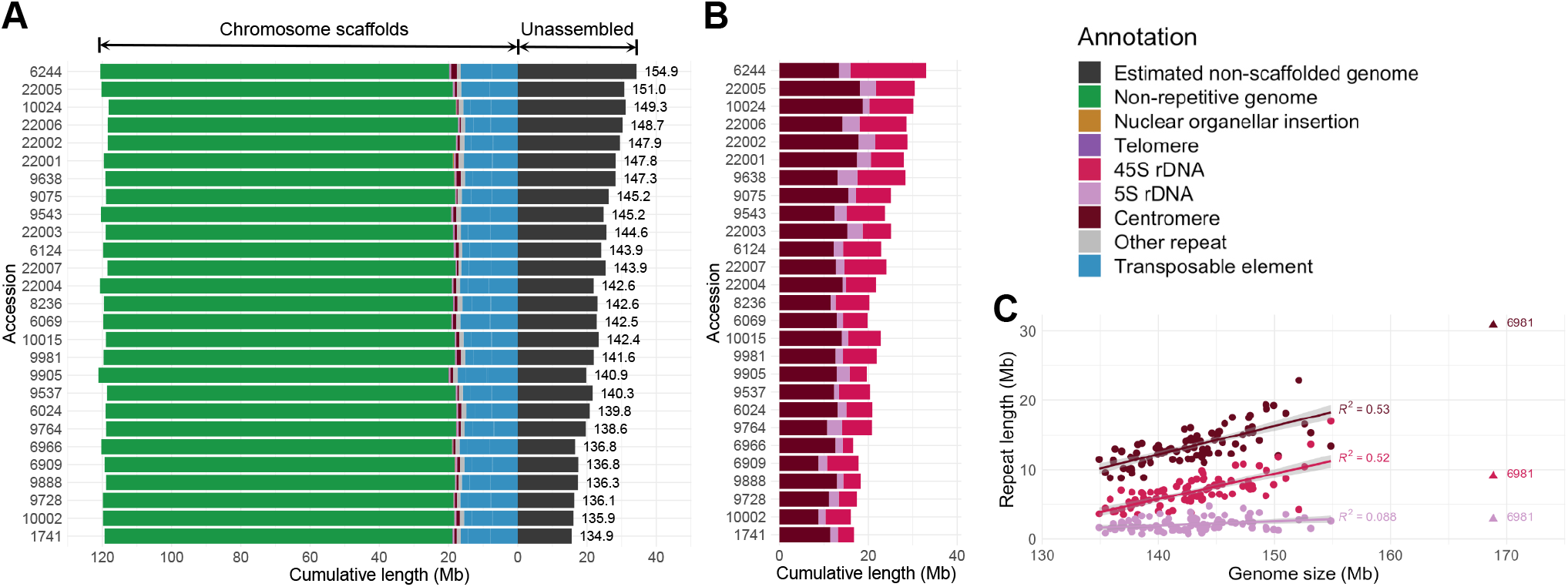
Genome assemblies and size variation. **A.**Histogram of genome sizes estimated from k-mers in PCR-free short reads with total assembly sizes superimposed (values to the left of “zero”). Most of the variation in genome size can be attributed to unassembled regions (values to the right of “zero”). **B**. The amount of centromeric, 5S rDNA and 45S rDNA repeats in the genome, as estimated from PCR-free short reads with a BLAST-based approach. **C**. Correlation between genome size and each of the three major satellite repeats. Their estimated amounts jointly explain up to 92% (*P* < 2.2 × 10^−16^) of total genome size variation (see Supplementary Note 1.3 for details). The regression line and estimates in the figure exclude accession 6981 (Ws-2; indicated with a triangle), because of its high centromeric repeat content (∼8 Mb).

### Detecting structural variation

In agreement with others ^23^, we find the chromosome arms to be not only similar in length across accessions, but also largely syntenic—as expected for a sexually reproducing organism with normal recombination (Fig. 2A). This said, thirteen Mb-scale rearrangements were readily apparent in whole-genome alignments. These include the previously described 1.2-Mb paracentric inversion associated with a heterochromatic knob on chromosome 4^49,50^, which is accompanied in a subset of the knob-less accessions by an 877-kb inversion that is 100 kb closer to the centromere (Extended Data Fig. 5; Supplementary Table 2), and a very large reciprocal translocation in accession 22001 from the Yangtze River region, which has swapped the distal portions of chromosomes 3 and 5 (Supplementary Fig. 1 in Supplementary Note 5). This latter rearrangement, which would presumably lead to decreased fertility in heterozygotes, appears to be rare, as the translocation was not detected in any of the other 117 accessions from the region that had previously been sequenced with Illumina short reads ^35^. In the analyses, we treated the rearranged segments as if they had remained in their original positions (and will refer to this modified assembly as 22001m; see Supplementary Note 1.1).

**Figure 2.**
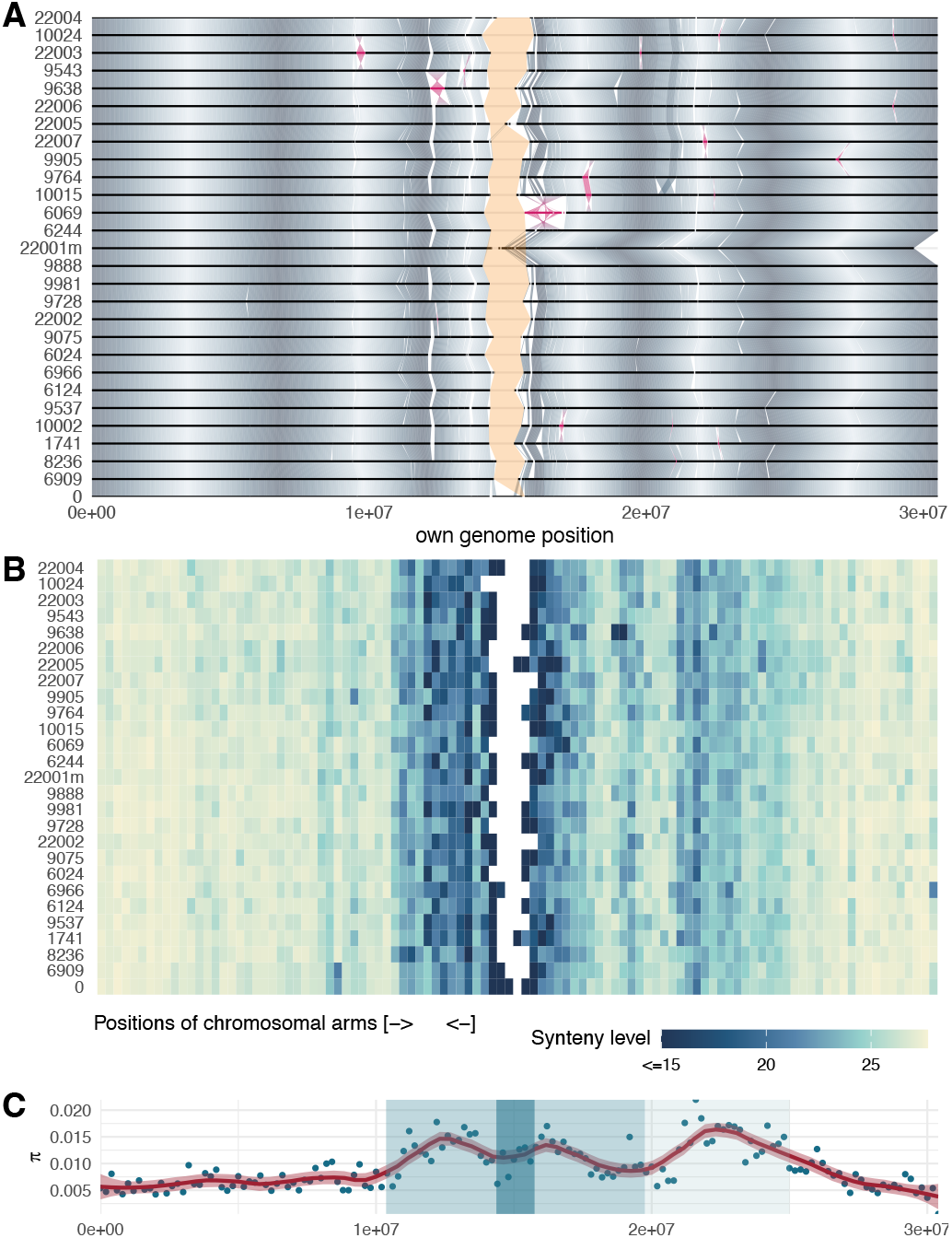
Distribution of variation on chromosome 1. **A.**Whole-genome alignment of the assembled parts illustrates that the genomes are generally structurally conserved away from centromeric regions. Chromosomes are aligned from both ends to emphasize the contiguity of the arms, with the unassembled centromeric regions indicated in yellow, and inversions in pink. Os-cillating gray shades highlight homologous regions (the periodicity of the gradients reflects repeated use of the color scheme, and has no biological meaning). For chromosomes 2-5, see Extended Data Fig. 5. **B**. The density of pan-genome graph bubbles, reflecting higher diversity in pericentromeric regions and at ∼ 20 Mb, where an ancestral centromere was lost through chromosome fusion since the divergence of *A. thaliana* from the rest of the genus ^46^. Synteny level refers to average sharing of links (1-27) between nodes across the 27 genomes in 300 kb blocks. For chromosomes 2-5, see Extended Data Fig. 6. **C**. Distribution of nucleotide diversity based on SNPs called from a multiple alignment. Dark blue corresponds to the centromeric region, with lighter blue highlights the pericen-tromeric area and lightest blue the ancestral lost centromere ^46^. For chromosomes 2-5, see Extended Data Fig. 7.

Obvious large-scale rearrangements aside, a comprehensive characterization of structural variants (SVs, by which we mean any alteration that causes variation in length, orientation, or local context of sequence) remains difficult. While SVs, along with SNPs, can be identified in genome alignments, the characterization of SVs is a fundamentally different problem from SNP-calling. The latter can be viewed as a technical issue—how to distinguish single-nucleotide polymorphisms from sequencing errors—but SV-calling is challenging even with flawless chromosomal sequences. The reason is that the SVs identified between genomes depend on the alignment method and parameters used, and there is no obvious ground truth. Given these uncertainties, we pursued two complementary approaches.

First, we developed a whole-genome multiple-alignment pipeline, Pannagram, that produces an intuitive representation for genome-browser visualization ^51^ (Fig. 2A). This approach can be considered an extension of pairwise-alignment methods ^52^ to handle multiple genomes in a reference-free manner. We derived a pan-genome coordinate system based on the resulting alignment and anchored it to the TAIR10 reference genome. Second, we used Pan-Genome Graph Builder (PGGB) ^29^, which yields a graph representation of multiple genomes by collapsing identical sequences into single nodes, connected by multiple paths forming bubbles where the genomes differ (Fig. 2B). The resulting object is computationally efficient for genotyping, but is also highly complex, with neither nodes nor bubbles having an obvious biological interpretation.

Comparing the SVs identified by these two conceptually approaches was not straightforward. SVs identified by Pannagram were typically covered by PGGB variants, which included nearly twice as much sequence, especially in highly polymorphic pericentromeric regions (Extended Data Fig. 8). A trivial reason for this difference is that Pannagram masked centromeric regions full of tandem repeat arrays (Fig. 2A-C), but we also identified several less obvious causes (see Extended Data Fig. 9–12 for details and examples).

First, the presence of physically distant but closely related sequences (*e*.*g*., reflecting recent TE activity) can lead to large loops in the PGGB graph that do not reflect actual SVs. Masking repetitive sequences will reduce this problem, but requires good repeat annotation—and would also make it impossible to study genome-variation comprehensively (one of our goals in this paper). Second, even in the case of tandem duplications, the graph combines duplicated sequences into a single node and hence counts all these sequences as part of SVs, even if not all of them are variable. Third, PGGB and Pannagram rely on different alignment parameters. To reduce the number of uninterpretable nodes, PGGB requires strict similarity criteria, whereas Pannagram can use more relaxed cutoffs to maximize homology detection. The *A. thaliana* genome contains many highly diverged regions ^53^, and these tend to be treated by PGGB as long SVs, whereas Pannagram often finds short alignments, resulting in local clusters of shorter variants. Whether Pannagram or PGGB results are more biologically relevant ultimately depends on the question and the cause of high divergence.

A final difference between PGGB and Pannagram lies in how nested length variants are represented. PGGB shows these as easily interpreted loops-within-loops, whereas Pannagram treats them as complex SV regions. This does not affect the size of the region covered by SVs, but does cause differences in the SV counts.

We emphasize that the differences between two algorithms designed to do different things should not be interpreted as bias, and that, as noted above, there is no ground truth. However, since Pannagram produces easily interpretable SVs along with convenient pan-genome coordinates, and all Pannagram SVs are covered by PGGB SVs, we based our further SV analyses on Pannagram.

### Characterization of structural variants

SVs come in many types and sizes, reflecting diverse mutational mechanisms. In what follows, we will focus on length variants, which are by far the most numerous (we identified fewer than a hundred inversions, for example). We further divide them into bi-allelic presence/absence polymorphisms, consistent with *simple* insertion/deletion (indel) mutations (sSV), and more *complex* multi-allelic polymorphisms (cSV) (Extended Data Fig. 10). We primarily consider the former, as they are easier to interpret, and also constitute the majority (over 80% overall), especially in the chromosome arms (Extended Data Fig. 13).

We identified 532,178 sSVs, affecting over 37.5 Mb in total—with the length-distribution being heavily skewed towards short variants (Fig. 3). To gain further insight into the nature of these polymorphisms, we consider sSVs between 15 bp and 20,000 bp length, because shorter variants are more sensitive to choice of alignment parameters, and larger ones are too few for statistical treatment. Determining the ancestral state of these variants using an outgroup species was in general impossible, as the homologous regions are typically intergenic and difficult to align. Nevertheless, the frequency distribution for the presence-allele of sSVs was consistent with sSVs mostly being due to rare insertions or rare deletions. Specifically, long presence alleles tend to be rare, and short ones common—suggesting that sSVs are mainly caused by long insertions and short deletions. As noted, both types are more frequent in intergenic than in genic regions, and they are also more often observed in introns than in exons, consistent with purifying selection removing many of them (Extended Data Fig. 14).

**Figure 3.**
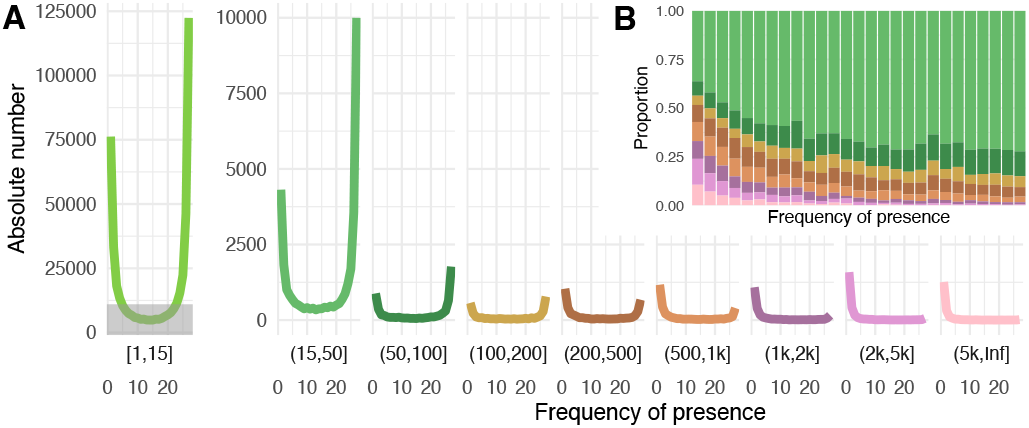
Structural variation. **A.**The frequency distribution of the presence alleles of all sSVs with length ≥ 15 bp, by length of variant. The height of the gray block in the left panel is equivalent to the height of the complete panel on the right. **B**. The proportion of length classes of sSVs in each frequency bin.

Finally, rare nuclear insertions of organellar DNA are easy to spot. We found 108 such insertions, almost all of them singletons or doubletons, ranging from a few hundred bp to entire organellar genomes (Extended Data Fig. 15, Supplementary Table 3). None of our genomes, other than our strain of the reference accession Col-0, 6909, shared the large nuclear insertion of mitochondrial DNA in chromosome 2 of the TAIR10 reference ^43,54,55^.

### Structural variants and annotated TEs

Active TEs produce SVs of diverse nature, both insertions and deletions, and serve as templates for various mutational processes *e*.*g*., double-stranded break repair ^56^. To investigate the role of TEs in generating SVs, we classified the presence alleles of 17,447 sSVs of length ≥ 100 bp based on their BLAST-identified coverage by ∼ 35,000 annotated TEs sequences, spanning ∼ 15% of the *A. thaliana* reference genome (*cf*. Fig. 1A). We defined different categories reflecting the extent of overlap (see Fig. 4A; for an analysis broken down by TE superfamily, see Supplementary Note 3). In total over 60% of sSVs showed some overlap with TE annotation, confirming a strong connection between our sSVs and TEs (Fig. 4B).

**Figure 4.**
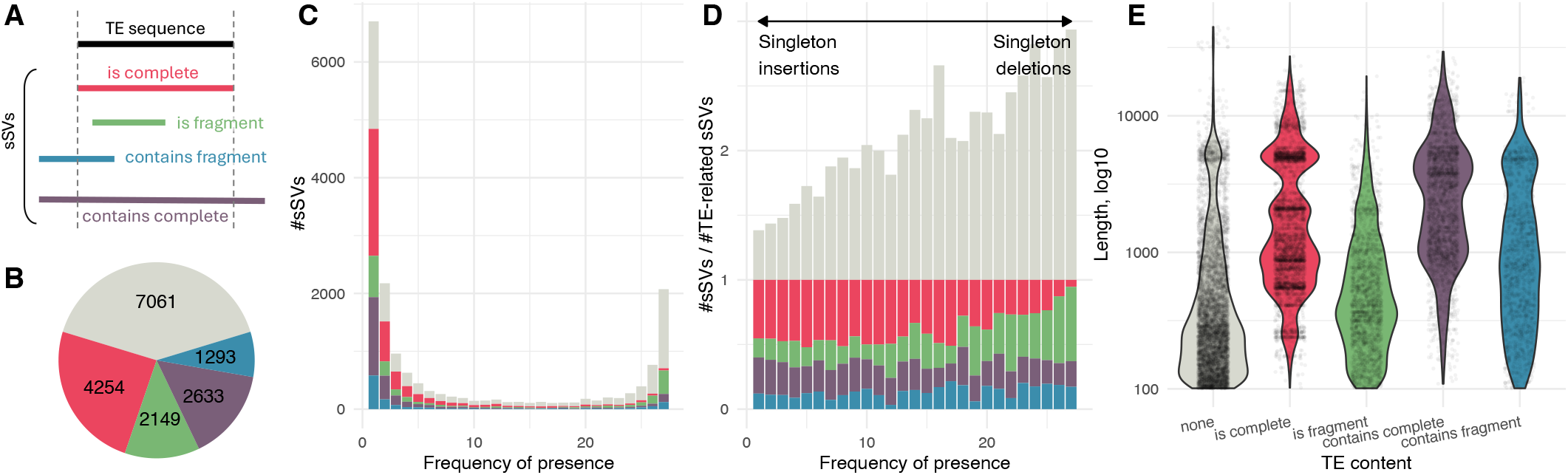
The role of TEs in bi-allelic indels. **A.**Categories of overlap between presence allele of sSVs and TE sequences from annotation. **B**. Total annotated TE content in sSVs. Grey, no TE content. **C-D**. Histograms of presence frequency showing annotated TE content as a function of frequency, either (C) raw counts or (D) relative to TE-related SVs, which are colored by categories, as shown in and (B). **E**. Distribution of the length of presence-alleles in different categories.

Likely insertions (presence allele in 1-3 accessions) tend to be longer than likely deletions (absence allele in 1-3 accessions). The former also more likely correspond to complete TEs, consistent with recent TE activity (Fig. 4C-D). The likely deletions more often correspond to incomplete TEs, suggesting that they are decaying elements (Fig. 4C-D). We tested this assertion by examining the regions flanking sSVs with similarity to TE fragments: 71% of likely deletions were within a TE, while the same was true for only 24% of likely insertions.

As expected, sSVs corresponding to complete TEs are enriched for particular lengths (specifically around 5 kb), reflecting activity of complete TEs (Fig. 4E). Similar patterns of enrichment were also found for sSVs in all other categories except for those corresponding to TE fragments. This supports recent reports that active TEs are far from perfectly annotated ^57,58^.

### The mobile-ome

Mobile elements that have been active in the history of our sample of genomes will have generated segregating insertions and deletions with similar sequence in different locations the genomes. We can use this property to look directly for mobile elements without relying on TE annotation. We will refer to the set of such elements as the mobile-ome, noting that our usage differs somewhat from that of others ^42^. To identify the mobile-ome, we used Pannagram to cluster all presence-alleles from sSVs based on sequence similarity, and represented the output as a graph of nestedness, where nodes represent sequences. The graph consists of many connected components, which we expected to correspond to distinct TE superfamilies or families (see Supplementary Note 3). We note that several similar approaches have recently been proposed ^59–61^.

Almost all sSVs that corresponded to complete TEs are part of the graph, consistent with their being part of the mobile-ome (Fig. 5A). Among other TE-related sSVs, approximately 70% are linked to the graph, and are thus also related to the mobile-ome, presumably reflecting a mixture of incompletely annotated TE families, complex insertion behaviors, and deletions within active TE families. The remaining 30% could result from low-activity TEs or decay of inactive TEs.

**Figure 5.**
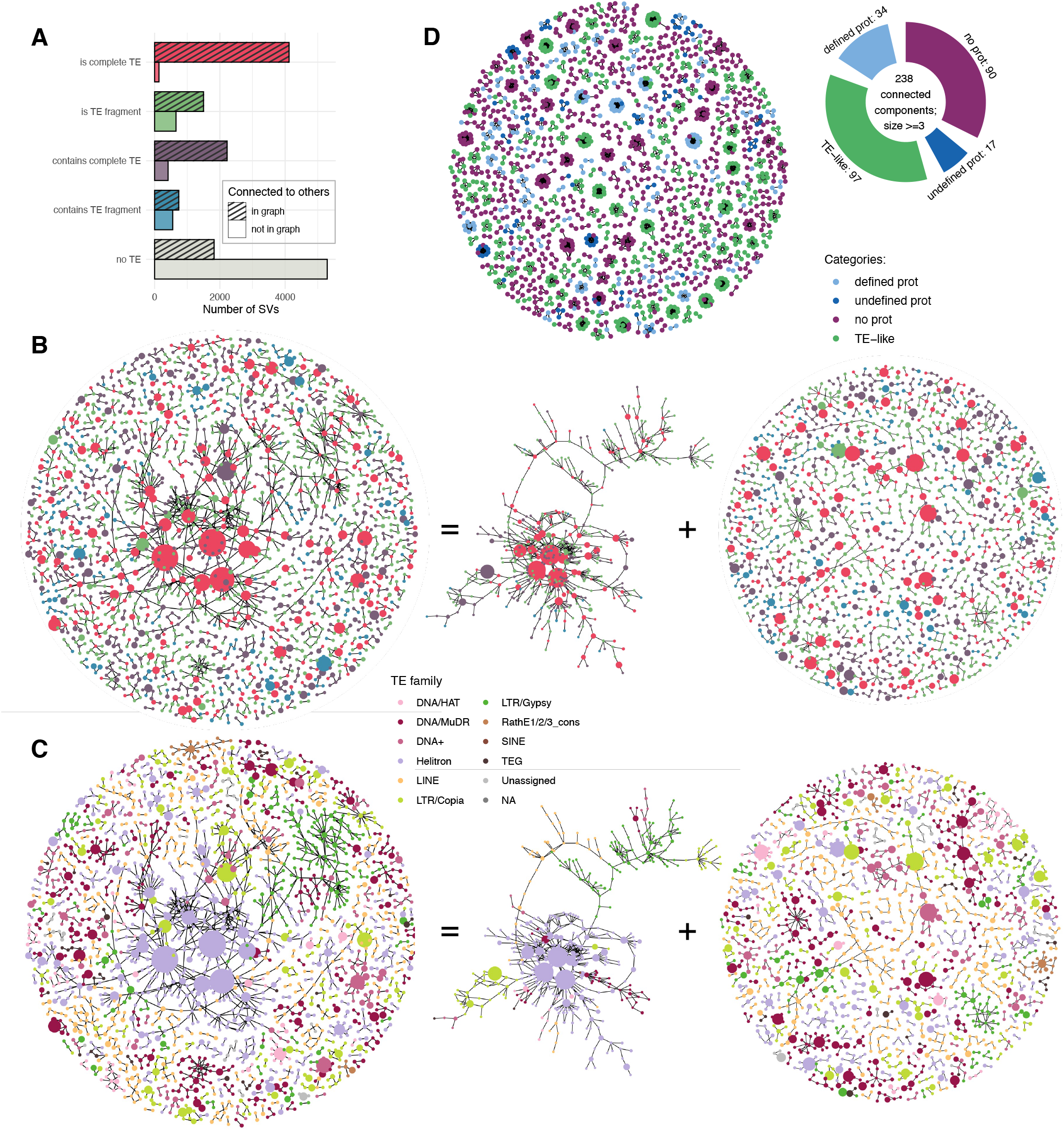
The graph structure of sSVs. **A.**The number of sSVs included (or not) in the graph of nestedness as a function of TE content. **B**. Graph illustrating the nestedness of sSVs, including only presence alleles with non-zero TE content. Each node represents a set of mutually nested sequences, using a similarity threshold 0.85, with node size reflecting the number of included sequences. Nodes are colored by TE content as shown in (A). **C**. The same graph as in (B), but colored by TE superfamilies, as shown in the center on top. Note that the algorithm used no prior knowledge of TE superfamilies. A large, dominant component connecting all TE superfamilies is also seen in a graph built using the entire *A. thaliana* TE annotation, suggesting that our sSVs and the reference annotation have comparable properties (Extended Data Fig. 17). **D**. A graph of nestedness for sSVs without TE content, colored by ORF content based on protein BLAST, as shown on the right.

To understand the relationship between annotated TEs, we filtered out sequences without TE content, and merged sequences with high similarity into larger nodes. The resulting subgraph consists of one dominant component along with numerous smaller ones (Fig. 5B). As expected, there are multiple large nodes corresponding to complete TEs, but we also see large nodes corresponding to sequences ostensibly containing complete TEs or TE fragments, demonstrating that many of these sequences are in fact part of the mobile-ome (see also Extended Data Fig. 16). Coloring the nodes by TE superfamily reveals that, while the smaller components mostly can be attributed to individual TE families, the dominant component contains members of all known TE superfamilies (Fig. 5C, Extended Data Fig. 17). Possible reasons include TEs inserting into existing TEs, and closely related sequences being mis-annotated as belonging to different superfamilies (*e*.*g*., Extended Data Fig. 18). Further evidence for the incompleteness of the TE annotation comes from small graph components that contain large nodes of longer elements that have only partial TE content. Closer examination of such nodes reveal many clear examples of un- or mis-annotated TE families (see Extended Data Figs. 19– 20).

In the set of sSVs without TE content, the majority (74%) is not connected in the graph, presumably reflecting unique events. Some are gene duplications; some may also be un-annotated TEs that are too rare in our limited set of genomes to be detectable by our method. Among the remaining 26%, we observed 238 connected components with ≥ 3 nodes, similar in structure to small components corresponding to single TE families in the analysis above (Fig. 5D). We hypothesized that these components contain previously un-annotated TEs. Protein BLAST analysis of ORFs from the sequences of these components revealed that 97 are indeed TE-like, with similarity to TE proteins from *A. thaliana* or other species. Among these, we observed a few TE families with evidence of horizontal gene transfer, as they have greater protein similarity to species outside a panel of five Brassicaceae species (*e*.*g*., Extended Data Fig. 21). We also found potentially new mobile element families lacking clear protein-coding potential. They form relatively large components in the graph (Fig. 5D, purple islands), composed of multiple sequences of similar length. They are not low-complexity and some are exclusive to *A. thaliana* (see Extended Data Fig. 22 for an example).

Annotated TEs in *A. thaliana* are generally epigenetically silenced. However, most published results have relied on a single reference genome, which makes it difficult to distinguish active from inactive TEs. Our mobile-ome data identifies segregating insertions corresponding to recently active TE families, and we can also consider the age of insertions, which should be proportional to their population frequency.

We investigated footprints of silencing in sSVs with existing methylation data ^62^. As expected, sSVs with annotated TE content (Fig. 4) are generally methylated, and at least three pattens support our conclusions above about the nature of these sSVs (Extended Data Fig. 23). First, sSVs corresponding to complete TEs are most highly methylated, followed by those corresponding to TE fragments. sSVs containing TEs or TE fragments are more variable, consistent with a subset of these sSVs corresponding to un- or mis-annotated TEs. Second, for sSVs corresponding to complete TEs, methylation increases with frequency, indicating that older insertion are more highly methylated. For sSVs containing TEs or TE fragments, this pattern changes at high frequencies, which could reflect high-frequency presence alleles not being insertions of un- or mis-annotated elements, but rather deletions that happen to contain TEs. Third, methylation is higher on sSVs that are part of the graph of nestedness (Fig. 5), consistent with methylation targeting the mobile-ome.

Finally, sSVs without annotated TE content behave similarly to sSVs containing TEs or TE fragments, but are on average less methylated, consistent with a smaller fraction of these sSVs corresponding to un-annotated TEs. Note in particular that sSVs that are part of the graph of nestedness are almost as highly methylated as previously annotated TEs, whereas those that are not tend to be completely unmethylated.

Expression patterns from existing RNA-seq data ^63^ are consistent with the methylation data: sequences in sSVs corresponding to complete TEs are barely expressed (except in pollen, where some TE expression is known to occur ^64^), while the behavior of sequences in other sSVs is more variable (Extended Data Fig. 24). Especially sSVs without annotated TE content are highly variable, with a small subset, presumably corresponding to protein-coding genes, having evidence of high expression levels.

### The gene-ome

To investigate how much the portion of the genome containing protein-coding genes—the “gene-ome”—varies among accessions, while minimizing reference-bias, we annotated each genome independently with sequence-based gene modeling and RNA-seq data from four tissues ^63^. To connect the annotations, we used the pan-genome coordinate system (rather than matching orthologs ^23^, which is the only possibility for distantly related genomes). Despite 83% of genetic loci having a one-to-one correspondence in the aligned genomes, gene model predictions varied considerably between accessions. While some of these differences likely represent genuine genetic variation, we lacked data to distinguish this from artifacts, and thus adopted a majority voting approach to harmonize annotations (Methods and Supplementary Note 4.1). After including 1,789 TAIR10 genes that were not re-identified in our study (unsurprising given that we used only four tissues for RNA-seq ^41,65^) and filtering out low-confidence genes (Methods), the final annotation contained 34,153 genes, ∼ 28,138 of which are *bona fide* protein-coding genes (Fig. 6A; Extended Data Fig. 25A; Supplementary Note 4.2). Of these, 2,661 were not previously annotated (for a detailed analysis of new genes, see Supplementary Note 4.3).

**Figure 6.**
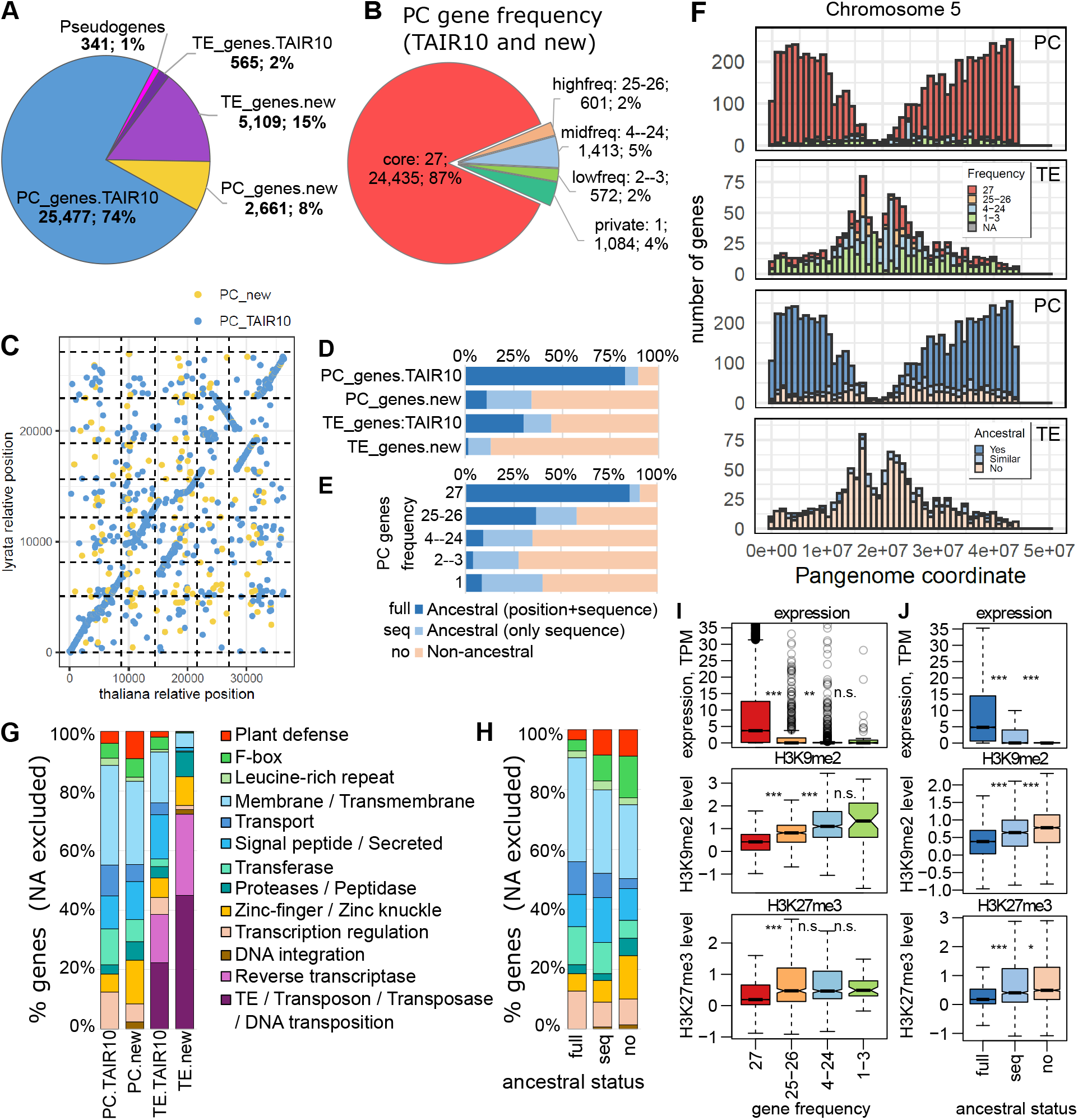
Analysis of the gene-ome. **A.**The 34,153 annotated genes categorized based on whether they correspond to the TAIR10 annotation and on whether they bear resemblance to TEs (based one sequence similarity and TE-like protein domains; see Extended Data Fig. 25A). PC genes, protein-coding genes. **B**. Genes categorized based on locus presence frequency across the 27 genomes. For TE gene presence frequency distribution, and a comparison with published results ^23^, see Extended Data Fig. 25B–C. **C**. Synteny comparison with *A. lyrata*. Genes are plotted by their consecutive gene IDs along the chromosomes. For TE genes, see Extended Data Fig. 25D. **D**. Ancestral status of genes of TAIR10 and newly annotated genes (“position+sequence” means that both gene sequence and syntenic position is conserved). **E**. Ancestral status of different-frequency protein-coding genes. **F**. Distribution of protein-coding and TE genes along chromosome 5, grouped by frequency (top) and ancestral status (bottom). For chromosomes 1–4, see Extended Data Fig. 26. **G**. Functional domains of annotated genes. Genes not matching a functional category have been excluded: for plots including these, and breakdowns by presence frequency see Extended Data Fig. 27. **H**. Functional domains of protein-coding genes grouped by ancestral status. **I–J**. Expression (in 9-leaf rosettes) and H3K9me2/H3K27me3 levels (in mature leaves) by gene frequency and ancestral status. Data shown ^63^ are for accession 6024; for other accessions, as well DNA methylation and 24 nt sRNA-coverage, see Extended Data Figs. 28–29. Significance estimates are from Mann-Whitney tests (^***^: *P* < 10^−10, **^: *P* < 10^−5, *^: *P* < 10^−2^, n.s.: *P* ≥ 10^−2^).

Focusing on presence-absence variation of genomic loci rather than annotated gene models, we found that 13% of loci were segregating in the population of 27 accessions (Fig. 6B; Extended Data Fig. 25B–C). This variation could reflect deletions (in some accessions) of ancestral genes, or segregating insertions of new genes specific to *A. thaliana*. To resolve this, we compared our genes to those in *A. lyrata*, to determine whether they were present and, if so, whether they were syntenic (Fig. 6C; Extended Data Fig. 25D). This analysis revealed a striking difference between TAIR10 genes and the previously un-annotated protein-coding genes: while most TAIR10 genes are ancestral, most newly identified genes are not (Fig. 6D). There was also a clear difference between segregating and fixed genes: as expected, most of the latter are ancestral, but the vast majority of the segregating genes is not (Fig. 6E). Although it is likely that we have underestimated the fraction of ancestral genes due to our reliance on the *A. lyrata* gene annotation, our results strongly suggest that segregating ancestral deletions are rare.

Segregating genes appear to be more common near centromeres, while fixed genes are clearly more common in the arms (Fig. 6F; Extended Data Fig. 26; *P* < 5 × 10^−4^ for all chromosomes using a Chi-Square test). Syntenic ancestral genes are also enriched in the arms, while other categories are evenly distributed across the genome. As expected, TE genes of higher frequency are more likely to be found near centromeres, consistent with selection against TE insertions in the arms ^42^.

To investigate functional enrichment in different gene categories, we searched for homologous proteins in UniProt KB (see Methods). We found that new protein-coding genes were enriched in defense and Zinc-finger genes, while TAIR10 genes were enriched in housekeeping functions such as transcription regulation and membrane proteins (Fig. 6G). A similar difference was found between ancestral and non-ancestral genes (Fig. 6H), as well as between fixed and segregating genes (Extended Data Fig. 27). Interestingly, new TE genes were strikingly more enriched for TE-function proteins than already annotated TE genes, suggesting that the former are more likely to be active TEs—which might be expected given that they are segregating.

Finally, there is a striking difference in expression levels between fixed and segregating genes: even genes that are only absent in one or two accessions tend to be almost silent in the remaining accessions (Fig. 6I). Consistent with this, epigenetic silencing is a function of population frequency: both gene-like Polycomb silencing (H3K27me3) and TE-like silencing (H3K9me2 and DNA methylation) are highest for genes of intermediate frequency (Fig. 6I). Likewise, non-ancestral genes have reduced expression and increased silencing (Fig. 6I–J; Extended Data Figs. 28–29), with increased H3K9me2 levels for all gene frequencies (Extended Data Fig. 29).

### The pan-genome

The term “pan-genome” is currently applied with a variety of meanings, from the original use in prokaryotes to describe the observation that genomes from different strains of the same species vary enormously in gene content ^66^, to the human genetics ambition of representing all polymorphism in a single “pan-genome graph” ^15,28^. We will discuss the utility of these concepts further below—here we simply consider how the pan-genome grows with sample size, and why. As shown in Figure 7A, all components of variation considered in this paper grow with sample size, but at different rates: the mobileome grows faster than the full genome and the gene-ome more slowly, consistent with the former being under stronger purifying selection. All components increase faster than the logarithmic growth expected under neutrality in a constant population, and more slowly than the linear growth expected in an exponentially growing population ^67^.

**Figure 7.**
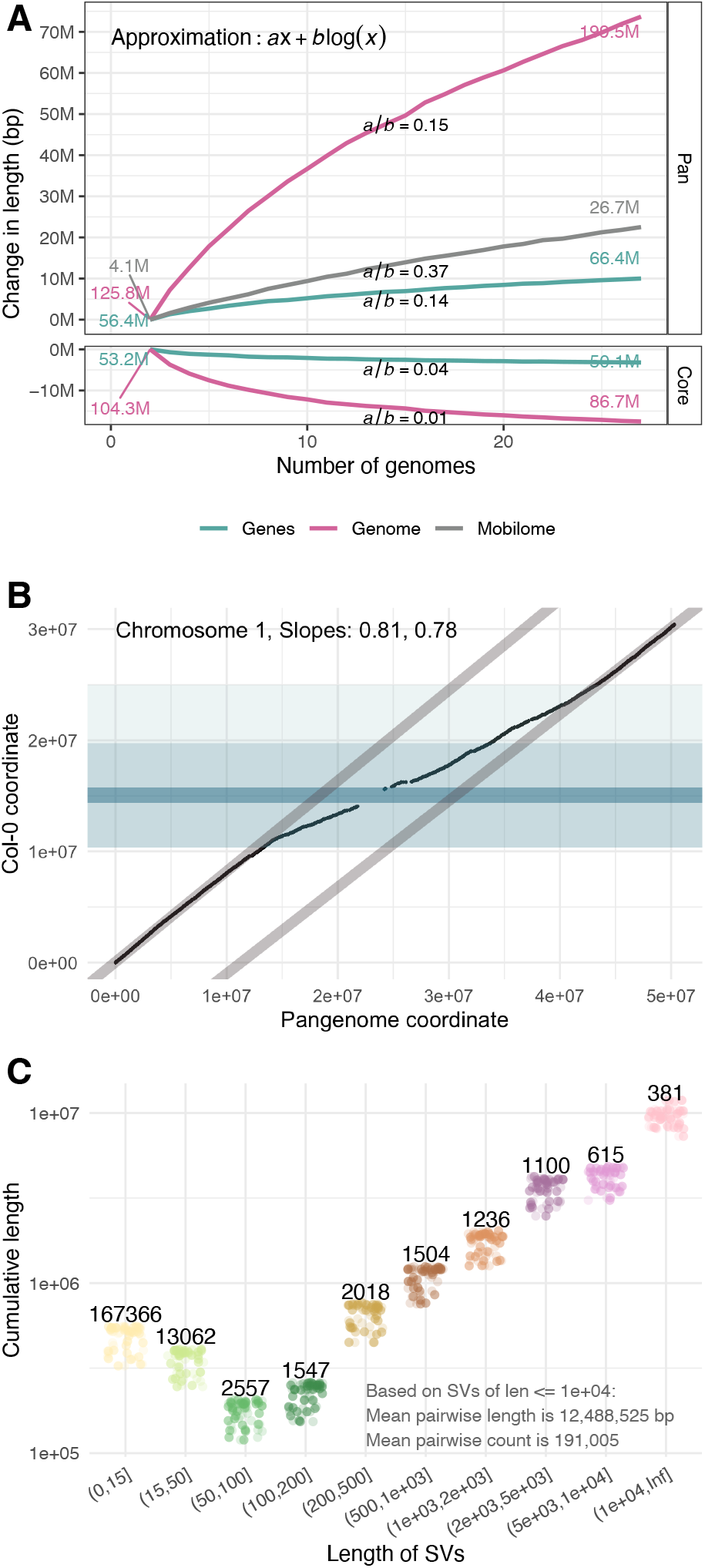
The growth of the pan-genome. **A.**The dependence on sample size for the union (“pan”) and intersection (“core”) sequence length, separately for the full genome, the mobile-ome, and the gene-ome (for saturation by sSV length and normalized views, see Extended Data Figs. 30–31). **B**. Pan-genome vs. reference genome coordinates. The pericentromeric region (light blue) shows a higher “dilution” of the spatial coordinates due to the increased number of SVs in this region, but note that the slope is never 1, reflecting their ubiquity. The centromeric region is dark blue (for other chromosomes, see Extended Data Fig. 32). **C**. The contribution of sSVs of different length to differences between pairs of genomes. Numbers are counts of sSVs. Shorter sSVs are more numerous, but longer and rarer sSVs contribute more to total length variation.

The growth of the pan-genome is reflected in the coordinate system and is not uniform along each chromosome, because most of the variation is found in centromeric regions (Fig. 7B). Already with 27 accessions, the pan-genome chromosomes are 63–76% longer than the TAIR10 chromosomes.

### Missing polymorphism

As part of the 1001 Genomes Project, we previously “resequenced” 1,135 accessions using short reads ^34^. We were well aware that the data were both incomplete and error-prone: we only called SNPs and short SVs, and only an average of 84% of the reference genome was covered by short reads from any particular accession.

With our whole-genome polymorphism data, we are now in a position to assess how much variation was previously missed. In the 1001 Genomes data, a pair of accessions differed, on average, at ∼ 440,000 SNPs. In our Pannagram alignment, the corresponding number is 600,000–800,000 SNPs, depending on how SNPs are defined (see Methods). In other words, we previously missed 25–45% of the SNP variation. In addition, whole-genome alignments of two genomes reveal on average ∼ 190k SVs (of length < 10 kb) covering a total over ∼ 12.5 Mb of sequence—approximately 10% of each assembled genome (Fig. 7C).

We investigated the causes of the missing SNPs by calling SNPs for our 27 genomes using the PCR-free, high-coverage short reads that were generated for this study, and comparing the results to those from pair-wise whole-genome alignment of complete assemblies (Fig. 8A and Supplementary Note 5). The main reason for missing SNPs from short reads is that the former lie in regions where no calls are made because the regions are not reliably covered by reads due to mapping problems. The extent of such regions depends on the mapping parameters used, but less conservative read-mapping will generally come at a cost of higher error rates. In our test SNP-calling with PCR-free data, we were able to reduce the fraction of missing SNPs to below 20%, but our FDR was then close to 7%. Consistent with a trade-off between conservative and aggressive read-mapping, *bona fide* SNP-calling errors, be they false positives or false negatives, were overwhelmingly due to read mismapping. Rampant pseudoheterozygosity caused by segregating duplications that are absent from the reference genome is particularly worrisome, in agreement with previous observations ^57^ (actual residual heterozygosity in inbred lines can be readily detected, but only one of our accessions had very limited amounts of residual heterozygosity, see Methods). Differences in local sequence alignment between algorithms also contributed to discrepancies, whereas traditional base-calling errors played an insignificant role (Supplementary Fig. 12 in Supplementary Note 5).

**Figure 8.**
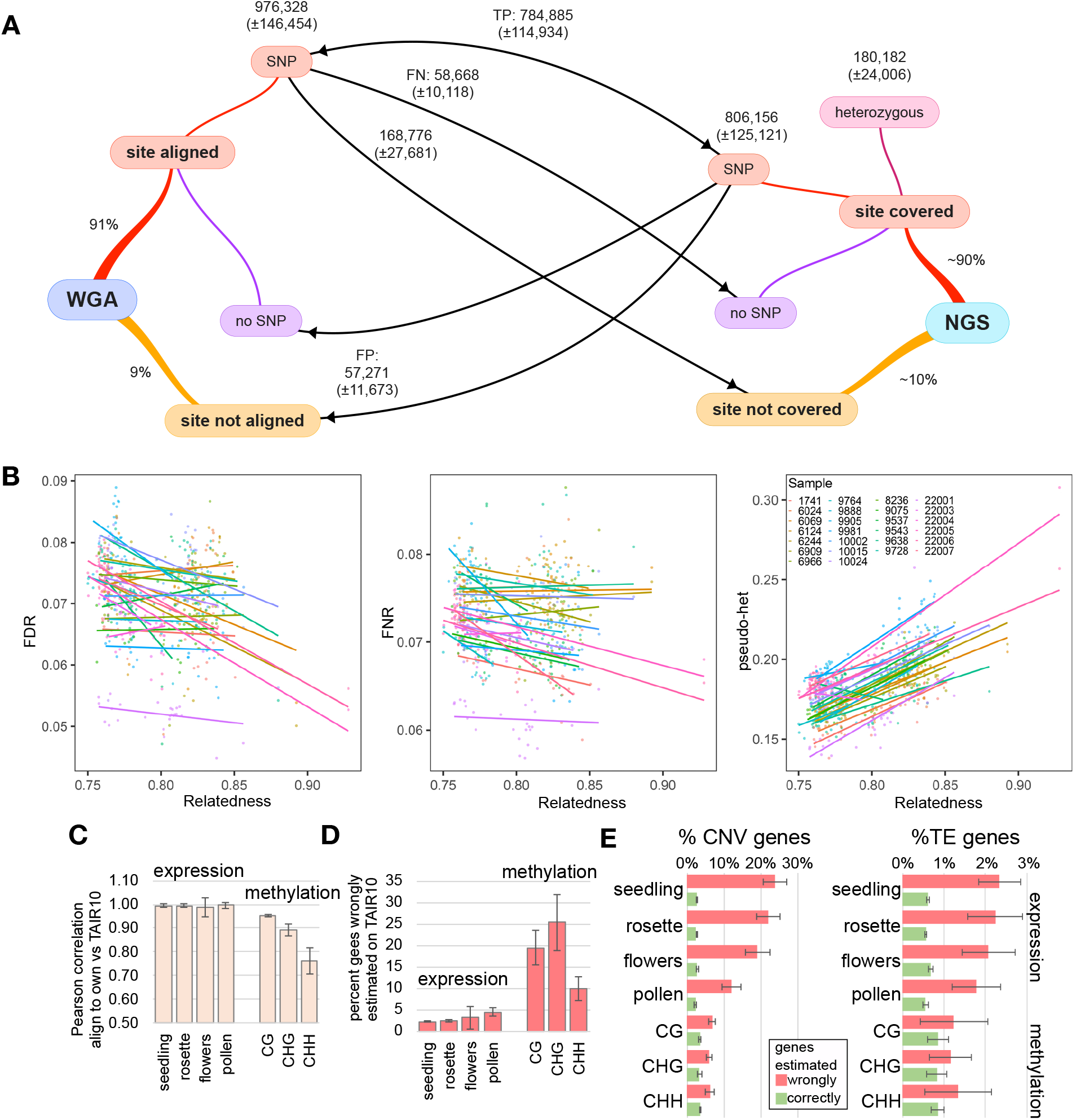
Read-mapping and reference bias. **A.**To investigate SNP-calling errors, SNPs were identified from pairwise whole-genome alignment (WGA) as well as by designating one genome as a reference and calling SNPs using short reads from the other (NGS). See Supplementary Note 5 for details. Numbers are averages across all accession pairs (± s.d.). From the point of view of WGA, each site can either be aligned or not. Conversely, from the point of view of NGS, each site in the reference genome can be covered by sample reads, or not. Arrows pointing right refer to WGA SNPs, which we assume to be correct; arrows pointing left refer to NGS SNPs. WGA SNPs are: true positives (TPs) if also called by NGS; false negatives (FNs) if missed by NGS; and uncalled if in regions not covered by NGS reads. Conversely, NGS SNPs are TPs if also found in the WGA and false positives (FPs) if not. The latter come in two flavors: those that correspond to *bona fide* non-SNPs in the WGA, and those that correspond to regions that were not aligned. Heterozygous calls in completely inbred lines are obviously FP and are treated separately. **B**. SNP-calling error rates depend on the relationship between the reference and the sampled genome. Each line is the regression of SNP-calling errors for a different choice of reference genome (identified by color). **C**. Correlation between expression- and methylation-level estimates derived from mapping reads to the TAIR10 reference genome and the genome from the sampled individual. **D**. Percentage of genes for which expression or methylation levels differ by more than 30% or 50%, respectively, when mapping reads to the TAIR10 reference instead of to their “own” genome. See Extended Data Fig. 33 for scatter plots. **E**. Enrichment of copy-number variable genes and TE genes among those with discordant expression- or methylation-level estimates.

### Reference bias

It is obvious that mapping short reads to a reference genome can cause reference bias, *i*.*e*., the results will depend on which genome is used as reference. Because we now have multiple genomes to choose the reference from, our data allow for systematic investigation of this problem. Starting with SNP-calling, we found that all SNP error rates depend both on the reference genome and on the relatedness between the reference and the sampled genome, often in unpredictable ways (Fig. 8B). Since many population genetic analyses rely on SNPs to estimate the relatedness between samples, this is troubling, and the consequences of non-randomly varying levels of bias in samples for downstream analyses such as GWAS or demographic inference merit further investigation.

Reference bias may also affect standard -omics techniques that quantify molecular phenotypes by mapping short reads to a reference genome rather than to the genome of the individual being studied. We illustrated this problem for transcriptome and methylome profiling by comparing the results of mapping RNA-seq and BS-seq reads to the TAIR10 reference as well as to the actual genome of the accession in question. Expression estimates between the two approaches were strongly correlated on average (Fig. 8C), but a subset of genes diverged markedly (Fig. 8D). These were strongly enriched for copy-number variable genes and TE genes (Fig. 8E). Methylome profiling was even more sensitive to the choice of genome for mapping, not surprising given that methylation in *A. thaliana* predominantly targets TEs, which in turn vary greatly in copy number between accessions and are therefore prime targets for cryptic mismapping ^57^. A scan for Differentially Methylated Regions (DMRs) in 100-bp windows across the genome revealed that profiling methylation by mapping reads to the TAIR10 reference rather than to each sample’s own genome produced a large number of spurious DMRs (Supplementary Fig. 17 in Supplementary Note 5). How serious this problem is will depend on the application, but it seems clear that it must be considered.

## Discussion

Over the last several years, population samples of more-or-less complete eukaryotic genomes have been appearing at an increasing rate, and there has been much excitement over the (variously defined) “pan-genome”, especially in plants ^68^. At the same time, it has become abundantly clear that a detailed and in-depth characterization of all the differences between individual genomes is very difficult. The problem is manifestly not a technical one, because even with perfect chromosomal sequences, we have to decide how to align them and how to interpret the differences. For complex structural variation, especially in highly diverged regions ^53^ where short stretches of similar or identical sequence are interspersed with variable stretches of diverged sequence, this is not trivial, and there is no obvious “gold standard” by which to evaluate algorithms. Furthermore, the ultimate reason for comparing genomes matters. If we are interested in using the pattern of polymorphism to answer questions about evolutionary history and mutational mechanisms, we must employ models that reflect actual historical events. In contrast, if the goal is simply to develop easily usable genetic markers, it may be irrelevant whether there is any correspondence between designated variants and the molecular processes that generated them.

With these caveats in mind, we tried two different approaches. First, we aligned the genomes using Pannagram ^51^, a whole-genome multiple-alignment pipeline, and created a pan-genome coordinate system anchored to the TAIR10 reference genome, facilitating the visualization of alignments in standard genome browsers (see Data availability). Second, we created a pan-genome graph with PGGB ^15,29^, which was developed primarily for human genomes. This matters because *A. thaliana* genomes have higher levels of polymorphism, stronger population structure, and many recently active TEs—all of which complicate graph building and interpretation. For these reasons, as well as general convenience, we based most of our analysis on the Pannagram output.

It has long been clear that SVs contribute substantially to polymorphism in higher organisms ^69,70^, and *A. thaliana* is no exception. In addition to massive variation in tandem repeat regions (Fig. 1), the readily alignable chromosome arms (Fig. 2) are highly polymorphic, with two accessions differing at ∼ 191,000 SVs covering ∼ 12.5 Mb on average. Although we also uncover massive amounts of new SNP variation, this still means that at least an order of magnitude more nucleotide sites are affected by SVs than by SNPs. The allele-frequency distribution of SVs suggests purifying selection, and also a mutational process that involves insertion of longer segments coupled with deletion of shorter segments. This is consistent with TE activity, and the overwhelming majority of presenceabsence variants longer than 100 bp involves annotated TE sequences. As expected in an organisms with active TEs ^42,71^, we found thousands of examples of what appear to be recent insertions of presumably complete TEs. Importantly, although many of these correspond perfectly to annotated TEs, many do not, demonstrating that our understanding of the mobile-ome remains highly incomplete—as is also becoming evident from long-read transcriptome analysis ^58^. To explore this further, we developed a graph-based analysis that identified several putative new TE families, providing a glimpse into how much is left to discover.

Turning to the gene-ome, we note that the term “pangenome” was originally invented to describe the rather fluid genomes of prokaryotes ^66^, which appear to be characterized by a relatively small number of essential (or “core”) genes, and a large cloud of presumably dispensable (or “accessory”) genes, which are shared even between distantly related taxa thanks to rampant horizontal gene transfer ^72^. It has been argued that this paradigm should apply to eukaryotes as well, for example in the context of rapidly evolving plant-pathogen interactions ^73^, but the overall picture in *A. thaliana* is clearly very different: the gene-ome is highly conserved, with 87% of genes detected in all 27 genomes, and the number of segregating genes growing considerably more slowly with sample size than other types of variation (Fig. 7). Furthermore, we distinguish between two types of segregating genes: a minority with homologs in the closely related *A. lyrata*, and a majority without (Fig. 6). The former, which correspond either to gene duplications or segregating deletions of ancestral genes, tend to be expressed at significantly higher levels than the latter, which often were characterized by TE-like epigenetic silencing. The extent to which these are genes with identifiable biological function will require further investigation.

Finally, we demonstrate that algorithms for SNP-calling, transcriptome profiling, and methylation profiling that rely on mapping short-read data to a reference genome can be highly biased, at least in a species with active TEs and many gene duplications. This is not surprising, but the problem may not have been appreciated fully, and it is likely to be worse in organisms with larger and more polymorphic genomes than *A. thaliana*. Also, outcrossing makes problems like pseudo-heterozygosity due to cryptic duplications far more difficult to detect ^57,74^. While it is still impractical to completely abandon the use of SNP-calling based on short reads in favor of whole-genome polymorphism data, we note that a recent paper that did this in order to estimate demographic parameters in *A. thaliana* reported that parameter-estimates changed two orders of magnitude ^75^! In general, utilizing at least a sample of diverse high-quality genomes for read-mapping should greatly help quantify the biases.

In conclusion, we recall Aravinda Chakravarti’s prediction at the dawn of the SNP era that models would be needed to “make sense out of sequence”—and that this would lead to “a rejuvenation of population genetics” ^76^. We think that the advent of unbiased whole-genome polymorphism data will have a similar effect. Most obviously, our understanding of TE dynamics will be revolutionized by our ability to see segregating TE insertions reflecting recent activity. While interspecific genome comparisons can reveal that bursts of activity have occurred, they lack the resolution to understand their dynamics, and cannot readily distinguish active TEs from the accumulated layers of dead and decaying TE sequences that litter most genomes.

More subtly, in order to make sense of complex polymorphisms, we need to understand the history of mutations that gave rise to them. The problem is analogous to phylogenetic analysis, where the estimated relationship between species is used to deduce the history of complex traits—what evolved first, and what evolved multiple times? In the present context, we need to estimate local coalescent trees (the so-called Ancestral Recombination Graph ^77,78^) and use them to infer the sequence of mutational and recombination events that gave rise to the sampled sequences. This analysis must be informed by a better understanding of the molecular mechanisms that cause structural variation. Although a considerable literature on phylogeny-guided statistical alignment exists ^79–83^, methods for doing this on a whole-genome scale, using an appropriate population genetic framework ^67^ and incorporating knowledge about molecular mechanisms, are missing.

In this context, it is important to remember that alignment-based methods developed for humans often do not work well in other species for a variety of reasons, including much higher levels of polymorphism and TE activity. *Arabidopsis thaliana* genomes are not unusual in these respects. In species like maize, where intergenic space is essentially un-alignable even between relatively closely related agricultural varieties ^69^, the idea of representing a whole-genome multiple alignment as a graph that captures all variation may be neither practicable nor useful ^84^.

## Methods

### DNA extraction and sequencing

For long-read sequencing, we began with 3-week-old plants grown in soil that had been transferred to darkness for 24-48 h before harvesting to reduce the starch content. 20-30 g of flash-frozen rosette tissue from pooled individuals was ground in liquid nitrogen with pestle and mortar. Nuclei were isolated as described for accession Ey15-2^43^, and high-molecular-weight (HMW) DNA purified with Genomic-tips 100G (Qiagen; #10243) following manufacturer’s instructions. 10 *μ*g of HMW DNA were sheared with either Megaruptor 3 (Diagenode; #B06010003) or a needle (FINE-JECT^®^ 26Gx1” 0.45×25mm; #14-13651) to ca. 75 kb, and used as input for long-read library preparation with the SM-RTbell Express Template Preparation Kit 2.0 (Pacific Biosciences; #101-693-800). These libraries were size-selected with the BluePippin system (SageScience) with a 30 kb cut-off in a 0.75% DF Marker U1 high-pass 30–40 kb vs3 gel cassette (Biozym; #BLF7510). Libraries for accessions 9981 (Angit-1; CS76366) and 10002 (TueWal-2; CS76405) were sequenced on a Sequel II system (Pacific Biosciences), and the others on a Sequel I system.

To prepare PCR-free libraries for short-read sequencing, the genomic DNA was fragmented to 250-350 bp using a Covaris S2 Focused Ultrasonicator (Covaris). The libraries were prepared according to manufacturer’s instructions with either the TruSeq DNA PCR-Free (Illumina, #20015962) or the NxSeq^®^ AmpFREE Low DNA Library Kit (Lucigen; # 14000-2). In total, libraries for 89 accessions (including the main 27 for which we assembled their genomes) were sequenced in paired-end mode on an HiSeq 3000 system (Illumina).

The ultra-HMW DNA extraction and sample preparation for optical maps was done as described ^43,85^ at Corteva Agri-science (Johnston, Indiana, USA) using the Direct Labeling and Stain (DLS) technology (Bionano Genomics).

### Assembly

The CLR subreads were assembled with Canu v1.71^38^. Since accessions 9981 and 10002 had been sequenced at higher coverage on a Sequel II instrument, only about 200X genome coverage worth of reads were used for assembly. We performed two rounds of polishing on the resulting contigs of all assemblies: first with the CLR subreads and Arrow v2.3.2 (https://github.com/PacificBiosciences/GenomicConsensus), and then with PCR-free short reads and Pilon v1.22^86^.

For scaffolding, we generated hybrid scaffolds with optical maps for eight accessions (Supplementary Table 1) using Bionano Access v1.5 and Bionano Solve v3.6. The assembly was performed in pre-assembly mode using parameters `non-haplotype` and `no-CMPR-cut`, without `extend-split`. Based on what we learned from these hybrid assemblies, we set the parameters for *in silico* scaffolding of the other genomes. We scaffolded contigs > 150 kb with RagTag v1.1.1^40^ (`scaffold -q 60 -f 10000 -I 0.6 -remove-small`) using the TAIR10 reference with hard-masked centromeres, rDNAs, telomeres, and nuclear insertions of organelles to prevent misplacement of contigs due to reference bias ^43^. All scaffolded assemblies were manually curated to specifically discard low-confidence centromere satellite-rich contigs or to invert contigs with satellite repeats at their edges indicative of their correct orientation. These edits were implemented in the agp files, which were converted to fasta format with the RagTag `agp2fa` function ^40^. To detect traces of residual heterozygosity, we aligned the original long reads to their corresponding chromosome scaffolds using pbmm2 v1.3.0 with the parameters `align -sort -log-level DEBUG -preset SUBREAD -min-length 5000`. Unmapped reads, as well as secondary and supplementary alignments, were filtered out using samtools v1.9 (`view -b -F 2308 <input.bam> Chr1 Chr2 Chr3 Chr4 Chr5`). The resulting BAM file was then analyzed with NucFreq v0.1 (`-minobed 2`) to assess genome-wide coverage of primary and secondary alleles ^87^. Agp files before and after manual curation, as well as NucFreq plots are available in the GitHub repository of this project.

### Repeat annotation

Repetitive elements were annotated as described ^43^. We ran RepeatMasker v4.0.9 (`-cutoff 200 -nolow -gff -xsmall`) using a custom library that included various consensus sequences for the CEN178^88^, 5S rDNA ^89^, 45S rDNA ^90^, and telomere repeats. We identified insertions of organellar DNA in the nuclear genomes by aligning the organellar genomes from the TAIR10 reference ^41^ with minimap2 v2.16^91^ (`-cx asm5`). We annotated tRNAs with tRNAscan-SE v2.0.6^92^, and TEs with Extensive de-novo TE Annotator (EDTA) v1.9.7^93^ (`--step all –sensitive 1 --anno 1`), a pipeline that combines several TE annotation tools (LTRharvest, LTR FINDER, LTR retriever, TIR-Learner, HelitronScanner, TEsorter) ^94–101^. Finally, to understand the causes of contig breaks, we determined what type of repetitive element was closest to each contig edge, considering the first 2 kb from each edge in contigs >10 kb.

### Pannagram

Pannagram is a toolkit designed for reference-free **pan**-genome alignment, **ann**otation, and **ana**lysis, as well as for generating dia**grams** ^51^.

We represent the whole-genome alignment as a matrix of corresponding positions, where rows represent accessions, and columns represent homologous positions. The construction of the alignment is done in a reference-free manner (see below). However, to visualize the alignment in genome browsers, columns must be sorted in some manner, *e*.*g*., to correspond to the TAIR10 sequence order. Then, columns of the pan-genome are used as positions in the pan-genome coordinate system.

To perform reference-free whole-genome alignment, we developed a 3-step pipeline. First, we use several accessions as reference and build draft pairwise alignments between each and all other accessions. This process results in several reference-based matrices of corresponding positions. Next, we intersect these matrices, selecting only those columns that are present in all reference-biased matrices, which produces reliable and reference-independent correspondences. In the final step, we resolve unaligned sequences between blocks of corresponding positions using multiple sequence alignment tools. Once the reference-free alignment is complete, it can be sorted according to the desired order of accessions. In our case, we employ an alphabetical order, with the TAIR10 genome first.

For the pairwise alignments between a reference genome (not necessarily TAIR10) and another accession, the focal accession genome is divided into blocks of 5,000 bp, and each block is then mapped to the corresponding chromosomes of the reference genome using BLAST, with exactly one best hit retained for each block through this process. Next, the BLAST hits that are not in close proximity to each other in both genomes are removed. An additional BLAST search is performed to align corresponding unaligned sequences between remaining hits.

To resolve any unaligned blocks after the reference-randomization procedure, MAFFT ^102^ is used. Blocks longer than 30 kb cannot be aligned within a reasonable time using MAFFT, so they are considered to be highly diverged. We found the final unaligned regions to be primarily associated with centromeric regions, rDNA clusters, telomeres and complex regions of multiple and long insertions and deletions, which are regions that are not of primary interest in this paper.

Given the whole-genome alignment, SNPs can simply be output as sequence differences. However, sequence differences can arise from ambiguities in local alignment and do not necessarily correspond to SNPs (see also Supplementary Note 5). If we take all sequence differences as SNPs, a pair of accessions differs at over 800,000 positions, on average, but if we restrict ourselves to isolated sequence differences, the number shrinks to 600,000.

### Pan-genome graph

#### Graph construction

We constructed genome graphs for each of the five chromosomes using the Pan-genome Graph Builder (PGGB) pipeline ^29^. First, we prepared the assemblies by splitting them into chromosomes and removing all unplaced contigs. To enforce linearity for simpler analysis and comparison, we used a modified version of accession 22001 with the genome rearranged to a consensus pan-genomic order (suffix: “f”). We added the TAIR10 reference genome to the graph to enable anchoring and presentation of results in a reference frame work.

We executed the PGGB pipeline (downloaded 25 January 2024) with the parameters `-s 10000 -p 90 -n 27`. PGGB consists of three methods: An all-against-all alignment with wfmash (v0.12.4-5-g0b191bb), graph induction using seqwish (v0.7.9-2-gf44b402), and two rounds of pangenome ordering (odgi v0.8.3-26-gbc7742ed) followed by normalization with smoothxg (v0.7.2-11-g9970e0d). The graph was used for the analysis of the pan-genome and synteny and for variation detection using vg deconstruct ^103^.

#### Similarity

We exploited graph properties to classify different levels of similarity between genomes. Nodes traversed in all accessions are labeled as core, nodes traversed in only one accession are private, and all other nodes (> `1 and` < `28 traversals`) are shell (soft). Note that nodes can be traversed multiple times by the same genome, which affects the total number of core nodes. Since each node contains a specific sequence, node count can easily be converted to the actual amount of sequence and respective genomic location.

#### Synteny windows

Every node in the graph can be translated to its exact position for each path. This direct connection allows us to create sliding window approaches for each sample/path using graph-based statistics. Here, we used non-overlapping windows of 300 kb and calculated the average similarity (see above) of these regions. This was performed for each graph and path independently and the results represented in a heat map.

#### Saturation analysis

A saturation analysis was performed using a bootstrapping approach. In each iteration we removed a specific number of paths from our graph and performed the same pan-genome categorization as above (s. 6.4.2 Similarity). In addition, we added the total pan-genome, which describes the total amount of sequence (core+shell+private sequence). We performed 20 different (unique) combinations for each size (number of genomes).

#### Deconstructing the graph

To achieve full insights into graph variation and cover all bubbles in the graph, `VG deconstruct` was run multiple times with each accession reference path once (`vg deconstruct -a -e`. After, the reported VCF (v1.54.0 “Parafada”) files were converted to a BED file with all important information provided. In addition, each chromosome was merged and genotype information was concluded and added. Bubbles were identified by start and end position, also reporting all traversals within these bubbles. Scripts can be found the in the repository.

### The mobile-ome

The mobile-ome refers to the collection of insertions and deletions that are likely to have occurred recently and are therefore not fixed in our sample. We hypothesize that each mobile event results in an SV, specifically a presence-absence polymorphism at the location of the insertion or deletion. Consequently, our initial approach involves extracting all presence-absence SVs and systematically decomposing them step by step. To distinguish between simple biallelic presence-absence polymorphisms (indels) and complex multi-allelic polymorphisms, we analyzed the lengths of alleles within the SVs. We distinguish two types based on the similarity threshold *s*, with *s* = 0.9 in our case. We consider a simple indel as one that contains alleles of two length types: those that are shorter than (1 − s) of the SV length (absence allele) and those that are longer than *s* of the SV length (presence allele). The distinction between simple and complex presence-absence polymorphisms is partially a computational construct to filter SVs and simplify further analysis. Simple indels and complex presence-absence polymorphisms form a continuum, and by relaxing the similarity threshold (*s* < 0.9, in our case), some complex SVs become classified as simple indels. Additionally, there is an natural bias towards complex presence-absence polymorphisms. Consider a scenario with a simple presence-absence polymorphism where an indel occurs within the presence allele. If the presence allele was initially observed in only one accession, then the new event does not reclassify the initial region as not belonging to the simple presence-absence polymorphisms category. However, if the presence allele was observed in multiple accessions, the new event is likely to be reclassified as complex presence-absence polymorphisms. To simplify and clarify the analysis, we considered only the simple polymorphisms. In order to determine the known portion of the mobile-ome within indels, we conducted BLAST searches using pan-genome consensus sequences of indels against known *A. thaliana* TEs, as well as against themselves. The indels that exhibited some similarity to known TEs were divided into the following groups:

**is complete** Significant similarity to known TEs and can be classified as TEs themselves.

**contains complete** Contained regions with similarity to known TEs but also additional sequences.

**is fragment** Contained only partially sequences with similarity to known TEs.

**contains fragment** Partial coverage by BLAST hits of TE segments but also additional sequences unrelated to known TEs.

We consider all these indels as parts of the mobile-ome. Indels without similarity to known TEs but showing nested similarities within the indel data set (where one sequence is a subsequence of another) were considered as potential candidates for new mobile-ome elements. In order to investigate their potential function, we obtained all six open reading frames (ORFs) within each of these indels. From each translated sequence, we selected either all continuous stretches without stop codons that were longer than 100 codons, or the longest stretch that exceeded 30 codons without a stop. Subsequently, we performed a BLAST search using the obtained amino acid sequences against the NCBI protein database and classified the potential proteins into four categories. If the BLAST results for an sSV contained keywords related to transposable elements (TE), we assigned the sequence to the TE-like category. These key words were: “transcriptase”, “reverse”, “transpos”, “gag-”, “pol-”, “integrase”, “gag/pol”, “gagpol”, “retrovirus’, “rna-directed dna polymerase”, “rnadependent dna polymerase”. sSVs that only had BLAST hits with descriptions such as “hypothetical protein”, “unnamed protein product”, “uncharacterized protein”, “predicted protein”, “PREDICTED:”, “putative protein”, and “unknown” were categorized as “undefined proteins”. Indels without any BLAST hits were classified as “no protein”. In all other cases, sSV was categorized as a “defined protein.”

### Gene annotation

#### Preliminary annotation

Gene annotation was mainly based on Augustus (v3.3.3) ^104^. Augustus-predicted gene models were trained using parameters obtained from “hints” from three different sources. First, we ran BUSCO (v4.0.1) ^105^ with `-m` genome option. Second, the *A. thaliana* reference gene annotation was projected onto each genome using Liftoff ^106^ with the `-exclude partial` and `-copies` options. Third, the RNA-seq data for each accession were used: wiggle hints were generated using `bam2wig` and `wig2hints` and EST hints were generated using `bam2hints` (all three tools provided by Augustus). Augustus was run with the following non-default parameters:

~~~
--softmasking 1
--species=BUSCO_retraining
--gff3=on
--extrinsicCfgFile=Custom_Config
--hintsfile=Liftoff_hints
~~~

For every accession, the GFF3 output of Augustus was run through the Augustus-provided tool `getAnno.pl` to translate gene annotations into protein sequences. Finally, for each annotation the Augustus output was combined and evaluated using `augustus GFF3 to EVM GFF3.pl` (provided by `EVidenceModeler` ^107^).

In addition to the Augustus-generated annotations, we used two types of independent evidence for gene models: from the SNAP *de novo* annotation tool ^108^ and from Cufflinks transcriptome assemblies ^109^. Annotations produced by Augustus, SNAP, and Cufflinks were combined and then subdivided into 1 Mb windows with 1 kb overlap using `partition EVM input.pl` (provided by `EVidenceModeler`). We ran `EVidenceModeler` with annotation gff files, the assembly fasta file, the partitions and a weight matrix. We chose weights for each input based on their ability to recreate the Araport11 gene annotation. Running `EVidenceModeler` produced the final annotation compilation for each accession. We retained only the longest isoform for each gene using `gffread` ^110^.

#### Reconciling annotations

To enable comparison between the independent annotations, we utilized the pan-genome coordinate system, reconciling discrepancies using majority voting. Additionally, we compared the sequences of each gene across different accessions. If a gene showed significant variation because it was located in regions heavily influenced by SVs, we excluded it from the analysis. Our approach generated a total of 34,153 putative genes: 3,438 of these were the result of splitting preliminary annotations; 1,020 were the result of merges.

For details, see Supplementary Note 4.1.

#### Ancestry analysis

All protein-coding sequences from all accessions were compared using DIAMOND’s `blastp` module ^111^ (version 2.0.11) against the *A. lyrata* MN47 proteome (version 2, Gen-Bank: GCA 944990045.1) and the best hit was considered as the *A. lyrata* homolog. To avoid bias due to mis-annotated genes in the *A. lyrata* proteome we further applied Liftoff v1.63^106^ to annotate all *A. thaliana* genes from all accessions on *A. lyrata* MN47 (v2, https://doi.org/10.6084/m9.figshare.22285444.v1) and *A. lyrata* NT1 (v2, https://doi.org/10.6084/m9.figshare.22293196.v1) assemblies. Next, each annotation group from *A. thaliana* was assigned to the *A. lyrata* homolog (by LiftOff or proteome similarity) that was common to at least 50% of its members, sharing at least 80% percent identity and covering at least 80% of the *A. thaliana* coding sequence. *Arabidopsis thaliana* annotation groups were defined to be ancestral relative to the *A. lyrata* gene, if they were part of a co-linear segment (of at least two) genes. To that end, all *A. thaliana* genes was ordered according to its relative position in the pan-genome coordinate system. Each pair of consecutive genes in *A. thaliana* was assigned to the same co-linear segment as its homologs in *A. lyrata*, if the homologs were separated by fewer than six genes. The ancestral state was defined as ‘similar’ for cases where the genes from *A. lyrata* and *A. thaliana* were not part of the same co-linear segment but shared at least 80% sequence identity over at least 80% of the length *A. thaliana* gene.

Further details in Supplementary Note 4.2 for TE analysis and Supplementary Note 4.3 for origin of new genes.

### Expression analysis

#### RNA-seq read mapping and gene expression calculation

Raw RNA-seq reads from 7-day-old seedlings, 9-leaf rosettes, flowers, and pollen ^63^ were aligned either to the TAIR10 reference genome or the corresponding accession accession genome using STAR v2.7.1^112^ with custom options:

~~~
--alignIntronMax 6000
--alignMatesGapMax 6000
--outFilterIntronMotifs RemoveNoncanonical
--outFilterMismatchNoverReadLmax 0.1
--outFilterMismatchNmax 999
--outFilterMismatchNoverLmax 0.3
--outFilterMultimapNmax 1
--alignSJoverhangMin 8
--outSAMattributes NH HI AS nM NM MD jM jI XS
~~~

(for read statistics, see Supplementary Table 4). Expression levels were assessed using `featurecounts` from Subread v2.0.1^113^ on each RNA-seq sample with either the TAIR10 gene annotation or the accession-specific annotations from this study. The entire locus including exons and introns was used for expression estimaties. Expression levels were normalized by calculating TPMs, the number of read counts divided by the gene length in kb and the total number of counts per kb for all genes divided by one million.

#### Expression of duplicated genes

The expression of gene copies was calculated only on the whole-locus level (disregarding exon-intron structure, as many copies were not part of our annotation and we did not have exon information) using the same method as abpve, but normalizing expression to the total number of counts covering the full annotation for a given sample and not the total number of reads covering the gene copies.

#### Mapping to TAIR10 vs. own genome

To determine whether the gene expression calculation was consistent between RNA-seq mapping in TAIR10 versus accession-specific genomes, we focused on the annotation groups with a one-to-one correspondence with an Araport11 gene. For each RNA-seq sample, we obtained the Pearson’s correlation coefficient between the exonic counts obtained from TAIR10 mapping and accession-specific mapping. We also determined the number of genes that were correctly or wrongly estimated using TAIR10 mapping. We called a gene “wrong” if the counts in TAIR10 and the counts in its own genome differed by more than 30% (`Ncounts min/Ncounts max` ≤ `0.7`). Only genes with at least 6 counts in either calculation were analyzed.

### ChIP-seq analysis

We used ChIP-seq data from 6 accessions and sRNA-seq data from 14 accessions ^63^. We used STAR ^112^ to map ChIP-seq reads with these non-default options:

~~~
--alignIntronMax 5
--outFilterMismatchNmax 10
--outFilterMultimapNmax 1
--alignEndsType EndToEnd
--twopassMode Basic
~~~

The ChIP-seq data were log2-normalized to input using bamCompare (deeptools package ^114^) using

~~~
--operation log2
--effectiveGenomeSize 119481543
--ignoreDuplicates
--outFileFormat bedgraph
~~~

The ChIP-seq coverage was estimated using bedtools map

-mean ^115^. The ChIP-seq coverage was further normalized to obtain value range similarity across accessions. For this, we applied quantile-normalization using an R function:

~~~
function(x) {
 (x-quantile(x,.20)) / (quantile(x,.80) - quantile(x,.20))
}
~~~

which equalized the 20% and 80% quantile values of each ChIP-seq sample. After quantile-normalization, the replicated samples were averaged.

### sRNA-seq analysis

We used sRNA-seq data for 14 accessions ^63^. To process the sRNA-seq data, we trimmed the reads using cutadapt ^116^: `cutadapt -a AACTGTAGGCACCATCAAT --minimum-length 18`. We then used STAR ^112^ with the following non-default options to map sRNA-seq reads to the corresponding genome:

~~~
--runRNGseed 12345
--alignEndsType Extend5pOfRead1
--alignIntronMax 5000 --alignSJDBoverhangMin 1
--outReadsUnmapped Fastx --outSAMmultNmax 100
--outSAMprimaryFlag AllBestScore
--outSAMattributes NH HI AS nM NM MD jM jI XS
--outFilterMultimapNmax 10
--outFilterMatchNmin 16
--outFilterMatchNminOverLread 0.66
--outFilterMismatchNmax 2
--outFilterMismatchNoverReadLmax 0.05
--outFilterIntronMotifs RemoveNoncanonicalUnannotated
--twopassMode None
~~~

We extracted 24-nt reads, calculated read coverage for each position of the genome using `genomeCoverageBed` (bed-tools v.2.27.1), normalized it by the total number of uniquely-mapped reads in each sample, and calculated 24-nt sRNA coverage for each locus of interest using `bedtools map -mean` function.

### DNA methylation analysis

To estimate DNA methylation levels, we used published BS-seq data for 12 accessions ^62^. After trimming with Trim-Galore (https://github.com/FelixKrueger/TrimGalore) with `--clip r1 10 --clip r2 15 --three prime clip r1 10 --three prime clip r2 10`, reads for each accession were mapped to its corresponding genome with Bismark ^117^ with `--score min L,0,-0.5` for a relaxed mismatch threshold and the `–un --ambiguous` parameters to obtain additional unmapped and multiply-mapping reads. Methylation was called as described ^118^. CG, CHG, and CHH methylation levels for genes and SVs in each accession were then calculated for each gene by focusing on all Cs in the specific context within the gene and calculating the ratio between the total number of methylated and unmethylated reads across all sites.

#### Mapping to TAIR10 vs. own genome

To estimate reference bias we mapped BS-seq data for all accessions to the TAIR10 genome and performed CG, CHG and CHH methylation level estimation in the same way as for own genomes. We then focused on annotation groups with a one-to-one correspondence with an Araport11 gene (the current annotation of the TAIR10 genome). We calculated Pearson’s correlation coefficient between the methylation level estimates obtained from TAIR10 mapping and accession-genome mapping. We also estimated the number of genes that were correctly or wrongly estimated using TAIR10 mapping. For each methylation context, we called a gene “wrongly estimated” if the methylation level in TAIR10 and the own genome differed by more than 50% (`methlevel min/methlevel max` ≤ `0.5`). For a more refined analysis of reference bias, see Supplementary Note 6.

## Data availability

The raw data (PacBio CLR and Illumina PCR-free short reads) and genome assemblies are deposited in the European Nucleotide Archive (ENA) under project accession number PRJEB73474 (ERP158243). Illumina PCR-free data for 61 additional accessions used to investigate the contribution of satellite repeats can be accessed under project accession number PRJEB73476 (ERP158245). In addition, assemblies, annotations, TEs, RNA-Seq, BS-seq, and ChIP-Seq data can be downloaded from 1001genomes.org, where a collection of accession-based JBrowse2 genome viewers and a pangenome JBrowse2 browser are also available.

## Code availability

All scripts can be found in an *ad hoc* 1001 Genomes+ repository. The Pannagram toolkit ^51^ can be found in a GitHub repository as can the Pangenome-graph methods. The genome assembly pipeline can be found in another GitHub repository.

## Acknowledgments

We thank Zhigui Bao, Alejandra Duque and Gal Ofir for discussion and comments on the manuscript. Elizaveta Grigoreva supplied the polymorphism data for Fig. 2C. This work was funded by the Deutsche Forschungsgemeinschaft (DFG), the Austrian Science Fund (FWF) and the Biotechnology and Biological Sciences Research Council (BBSRC) through ERA-CAPS Project 1001GenomesPlus (BBSRC BB/S004661/1; P.K., D.W., M.N.), the Austrian Academy of Sciences (M.N.), the Max Planck Society (D.W.), and the European Union’s Framework Programme for Research and Innovation Horizon 2020 (2014-2020) under the Marie Curie Skłodowska Grant Agreement Nr. 847548 (H-J.L.) as well ERC Advanced Grant EPICLINES to M.N.

## Author contributions

P.K., D.W. and M.N. conceived and supervised the project. P.F., S.N, A.H. and Y.G. contributed materials. F.R., A.K., M.C., A.M., V.L., V.N., I.R., C.L, and F.B generated data. F.R., H.A., B.J. and T.W. assembled the genomes. A.I., H.A., M.C. and C.K. annotated the genomes. H.L. and A.K analyzed expression and epigenetics. A.I. and S.V. performed the SV analysis. A.I. analyzed the mobile-ome. A.I. and A.B. analyzed the pangenome alignment. A.I., H.A., A.K., and V.V. analyzed the gene-ome. H.L. and A.K. analyzed short-read mapping bias. J.F. developed the genome browser. A.I, S.V., F.R., H.L, H.A., A.K., D.W. and M.N. wrote the paper.

## Extended Data Figures

**Extended Data Figure 1.**
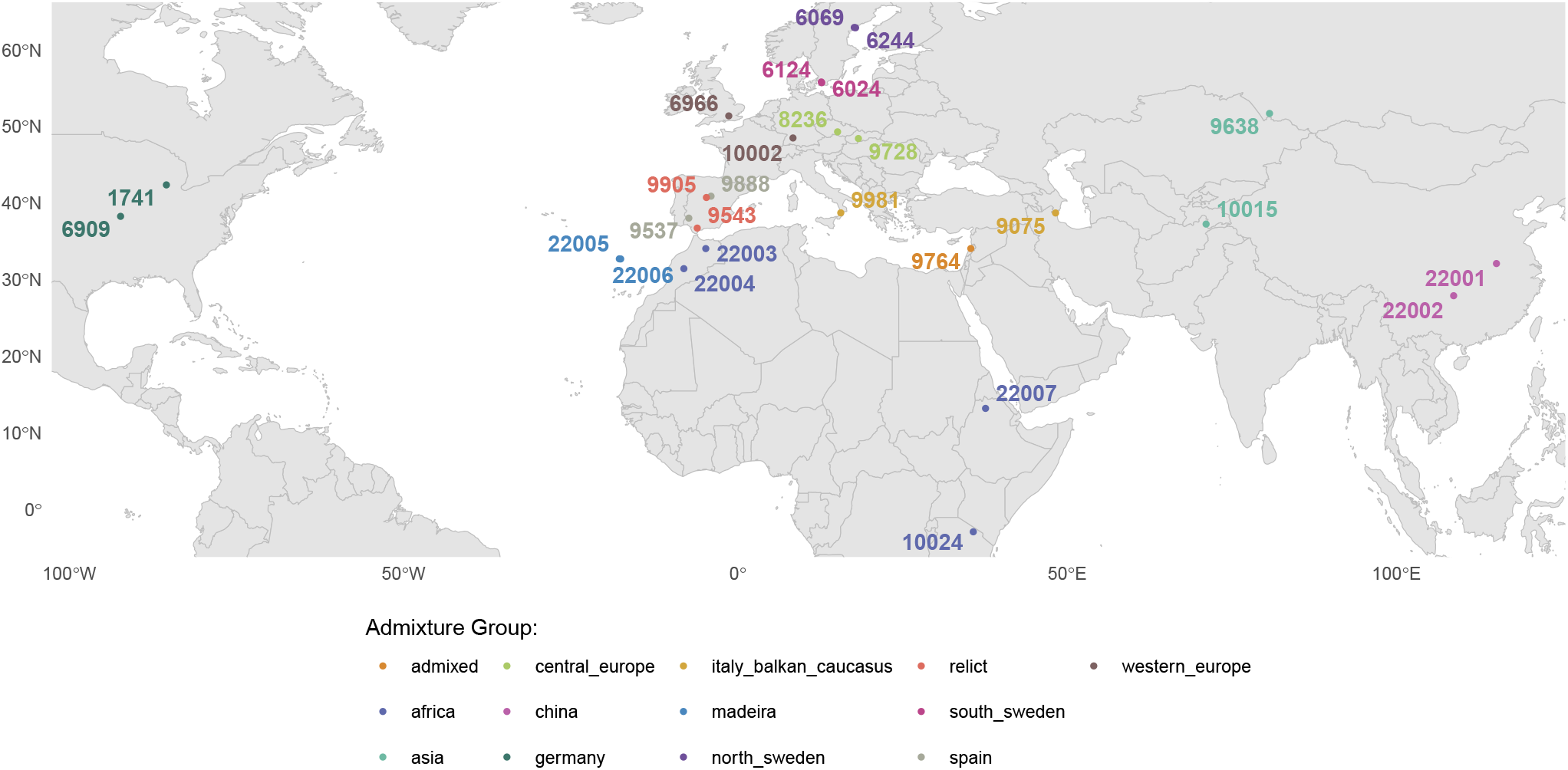
Origin of the sequenced accessions. Colors indicate “ADMIXTURE group” ^34^. Note that 6909, which corresponds to the TAIR10 reference genome (Col-0) lacks reliable collection data (but hails from Central Europe based on genotype as well as historical records).

**Extended Data Figure 2.**
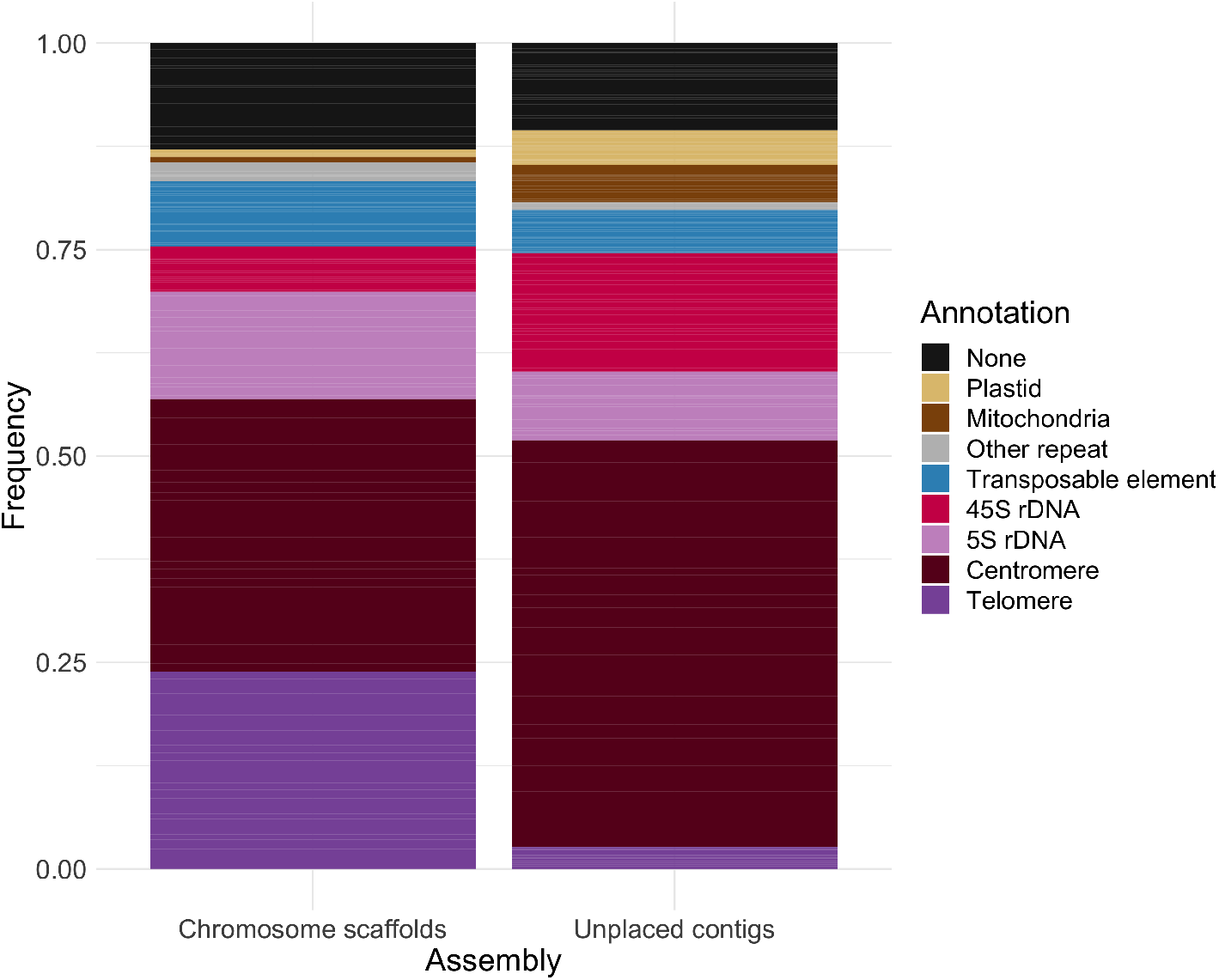
The causes of contig breaks. Stacked bar chart summarizing the type of repetitive element closest to each contig edge across the 27 assemblies, separately for scaffolded and unplaced contigs. 72% of the latter end with centromeric or rDNA repeats.

**Extended Data Figure 3.**
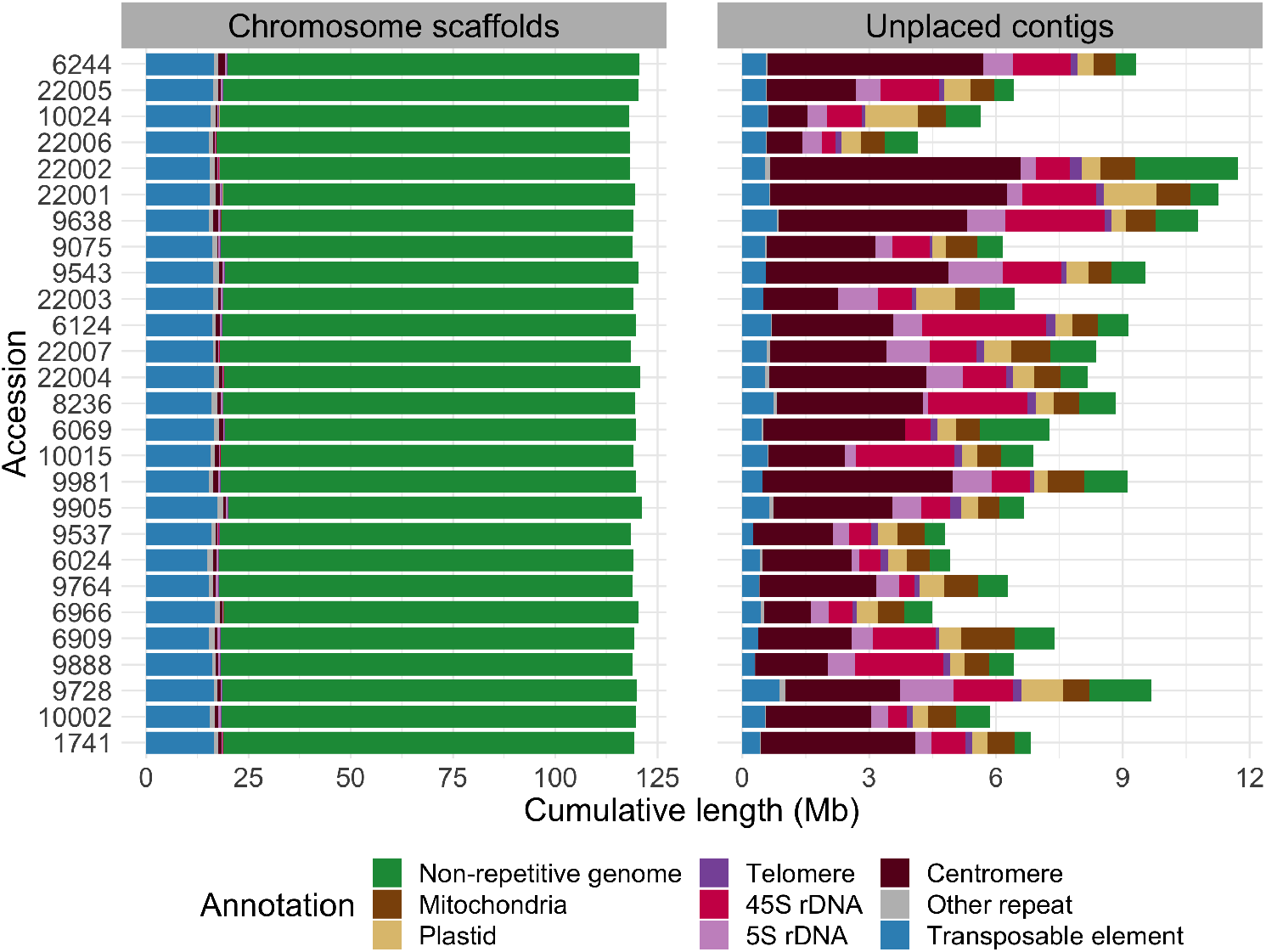
Scaffolded vs. unplaced contigs. The former correspond to the chromosome arms and contain mostly non-repetitive sequence and TEs, while the latter mostly contain centromeric and rDNA repeats, as well as organellar DNA sequence (*cf*. Fig. 1).

**Extended Data Figure 4.**
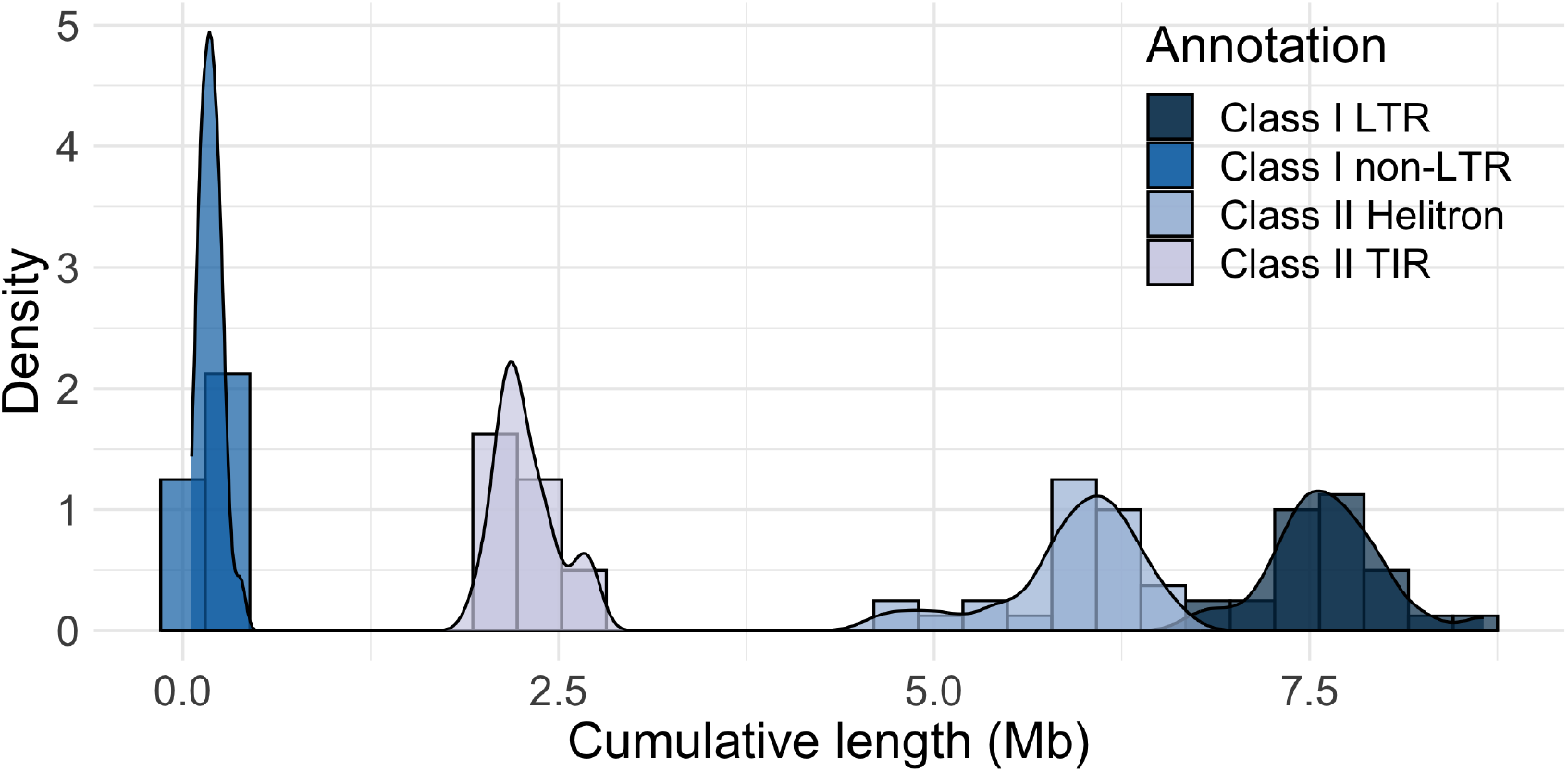
TE class size distribution. The cumulative length of all copies of a given TE class differs greatly across accessions, but the total TE content does not.

**Extended Data Figure 5.**
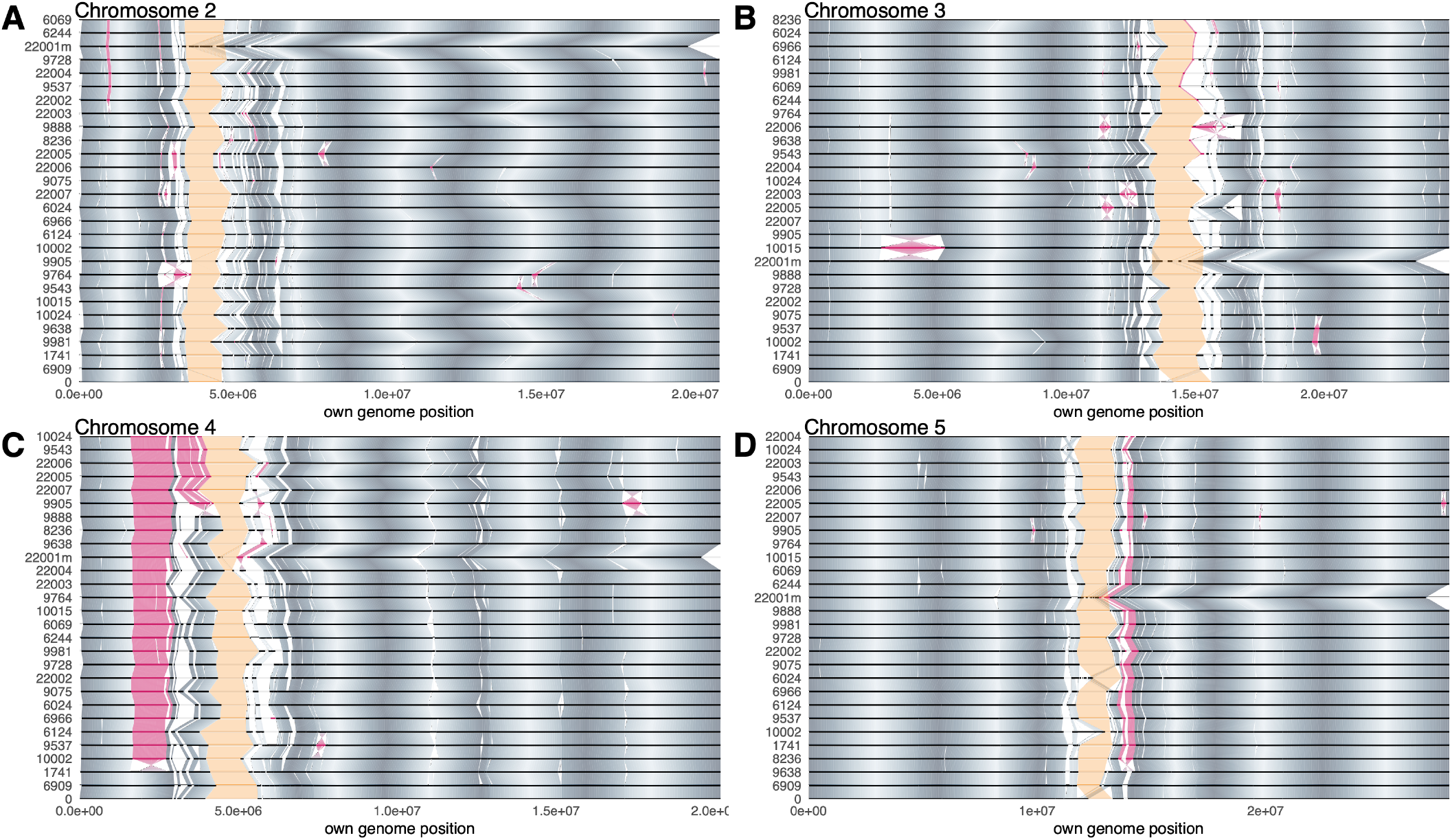
Whole-genome alignments. **A**. Chromosome 2. **B**. Chromosome 3. **C**. Chromosome 4. **D**. Chromosome 5. For chromosome 1 and further explanation, please see Fig 2A.

**Extended Data Figure 6.**
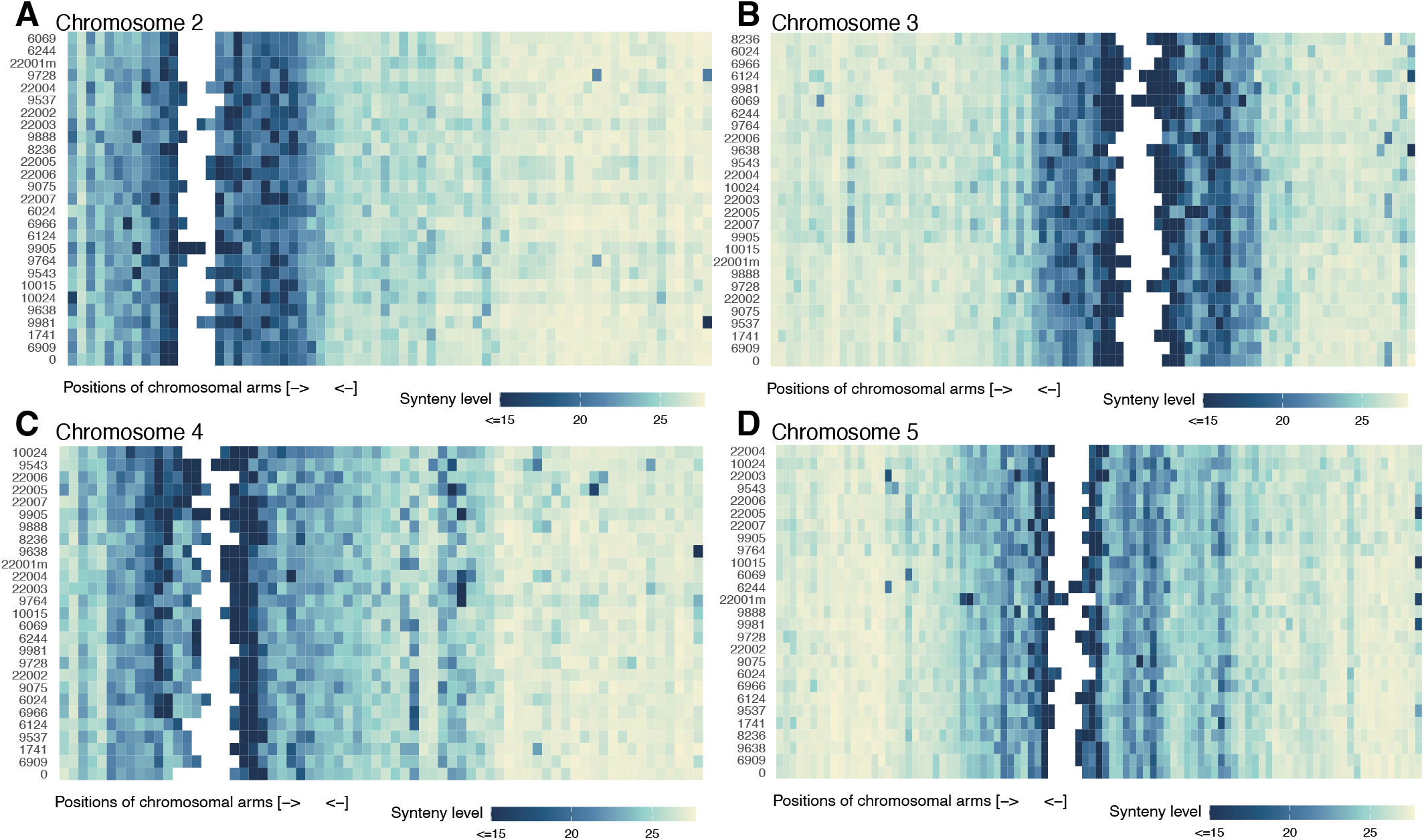
Density of PGGB graph bubbles. **A**. Chromosome 2. **B**. Chromosome 3. **C**. Chromosome 4. **D**. Chromosome 5. For chromosome 1 and further explanation, please see Fig 2B.

**Extended Data Figure 7.**
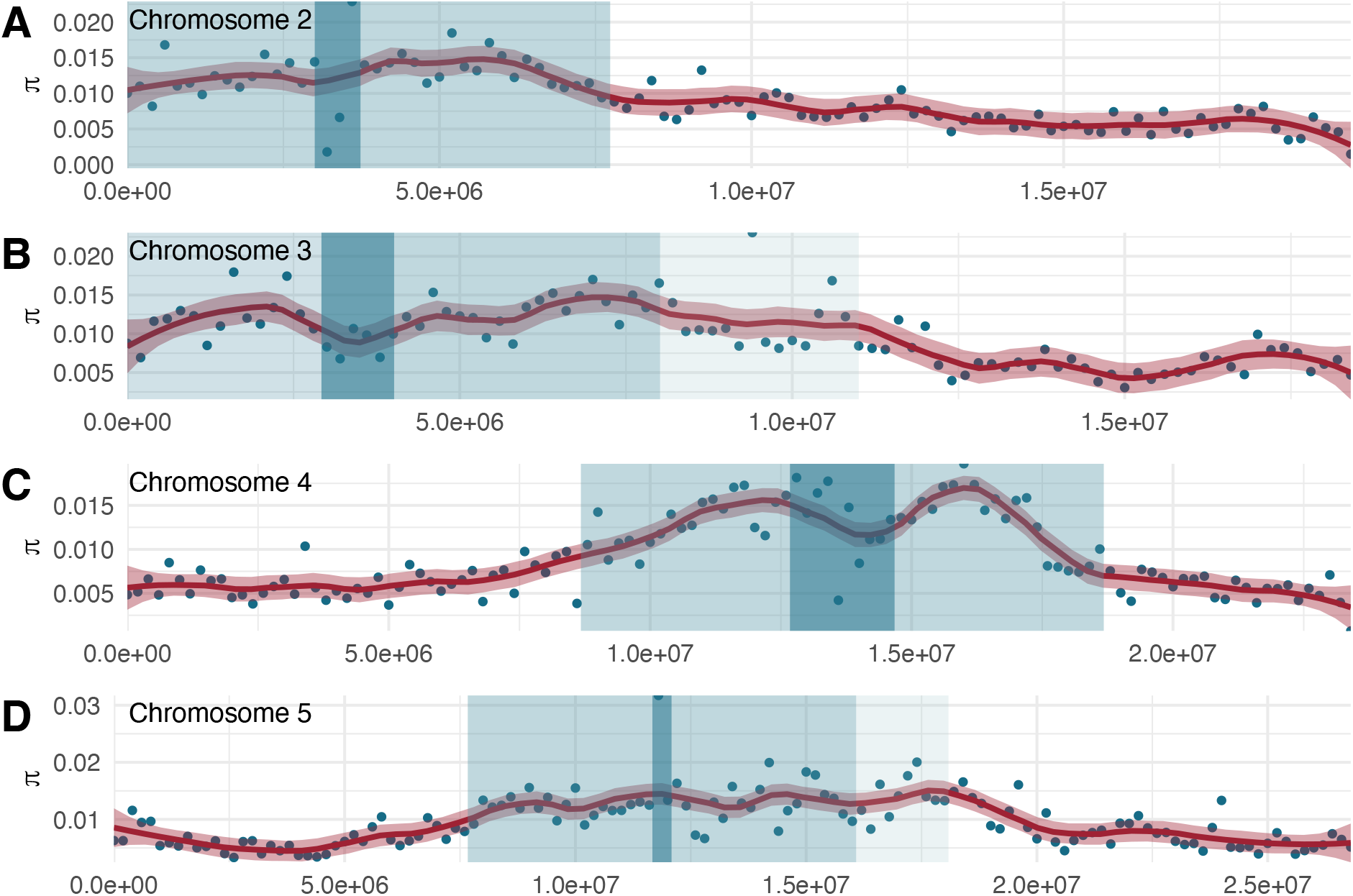
Nucleotide diversity across the genome. **A**. Chromosome 2. **B**. Chromosome 3. **C**. Chromosome 4. **D**. Chromosome 5. For chromosome 1 and further explanation, please see Fig 2C.

**Extended Data Figure 8.**
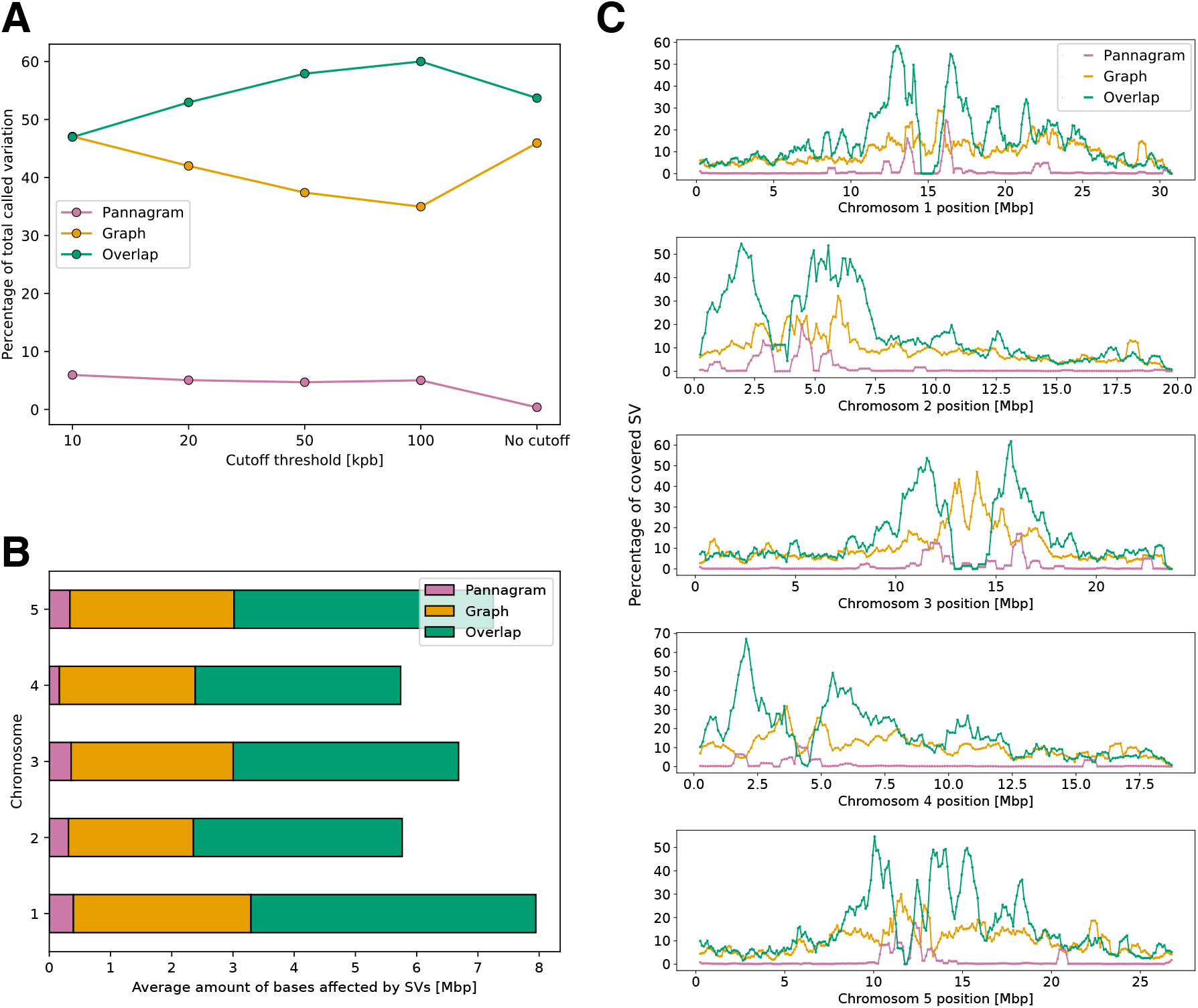
Comparing SVs from Pannagram and PGGB graphs. **A**. Scatter plot of overlap and method-specific SVs as a function of eliminating SVs above a certain length-cutoff. Overall, the overlap between the two methods is 50%, but the overlap can be increased by removing large SVs (demonstrating that disagreement is disproportionally due to large SVs. **B**. Comparison of Pannagram and graph-based SVs across chromosomes (average per accession), demonstrating that there are no major differences between chromosomes. SVs shorter than 15 bp were not included in this figure. **C**. Position of overlapping and method-specific SVs for each chromosome of accession 6909 (Col-0). Large discrepancies are more pronounced close to the centromeres (there is no overlap inside centromeres, as these are masked by Pannagram). Each dot represents a 100 kb window, using a moving average of five windows.

**Extended Data Figure 9.**
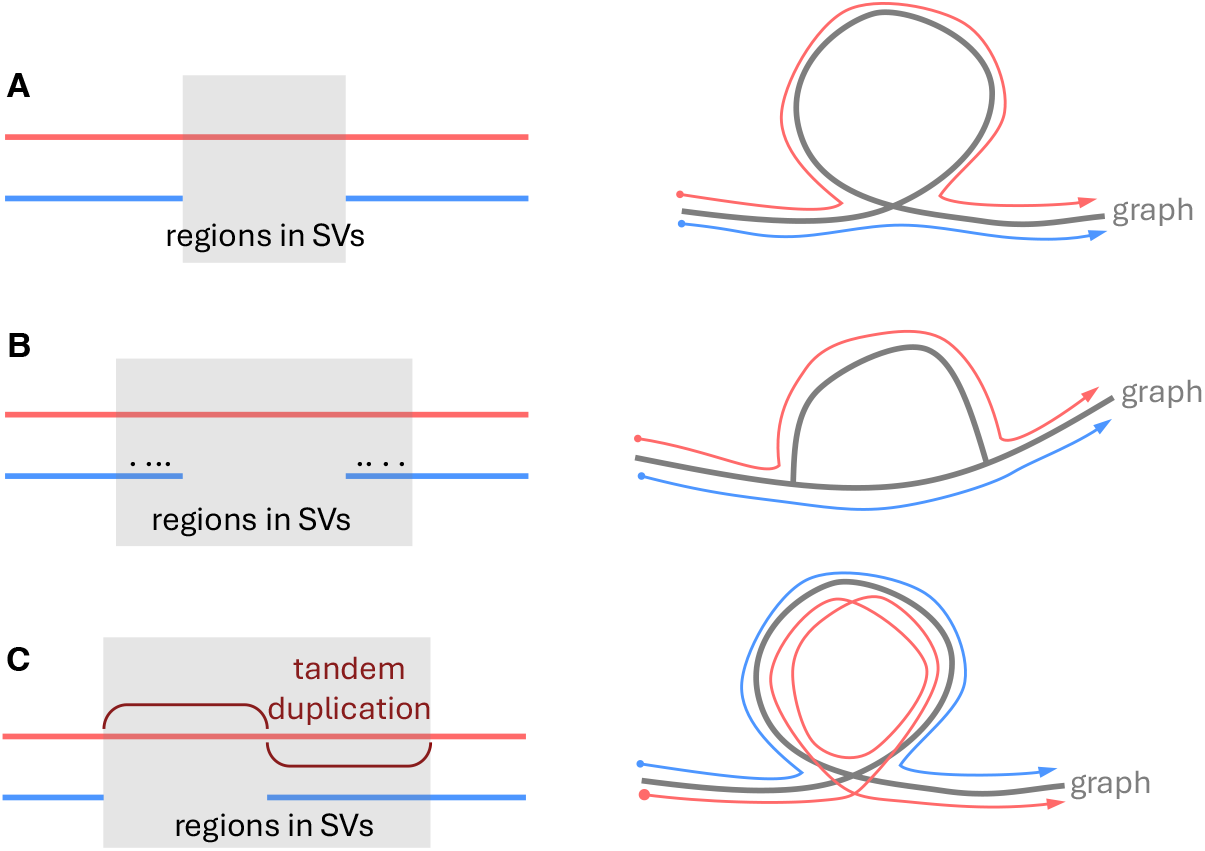
Cartoons illustrating cases where graph SVs are longer than Pannagram SVs. **A**. Two genomes (red and blue) differ by a single simple SV (Extended Data Fig. 10), which can be represented as a gap in the alignment, or a loop in the graph. Pannagram and the graph give the same result and the length of SV (indicated in grey) is identical. **B**. However, if SNPs, represented by dots, are linked to the SV, causing imperfect alignment in the flanking regions around the SV, PGGB may recognize longer haplotypes, resulting in an arrangement that resembles a hat. In this case, the graph SV is not merely a presence-absence variant, but a complex SV with two alleles. The entire region affected (in grey) is longer than the SV recognized by Pannagram (still the same as in A). **C**. When the SV is formed by a tandem duplication, the graph representation of the SV is topologically similar to scenario A, but the SV covers both the original sequence and its duplicated copy (grey region), while the SV identified by Pannagram is still the same as in A.

**Extended Data Figure 10.**
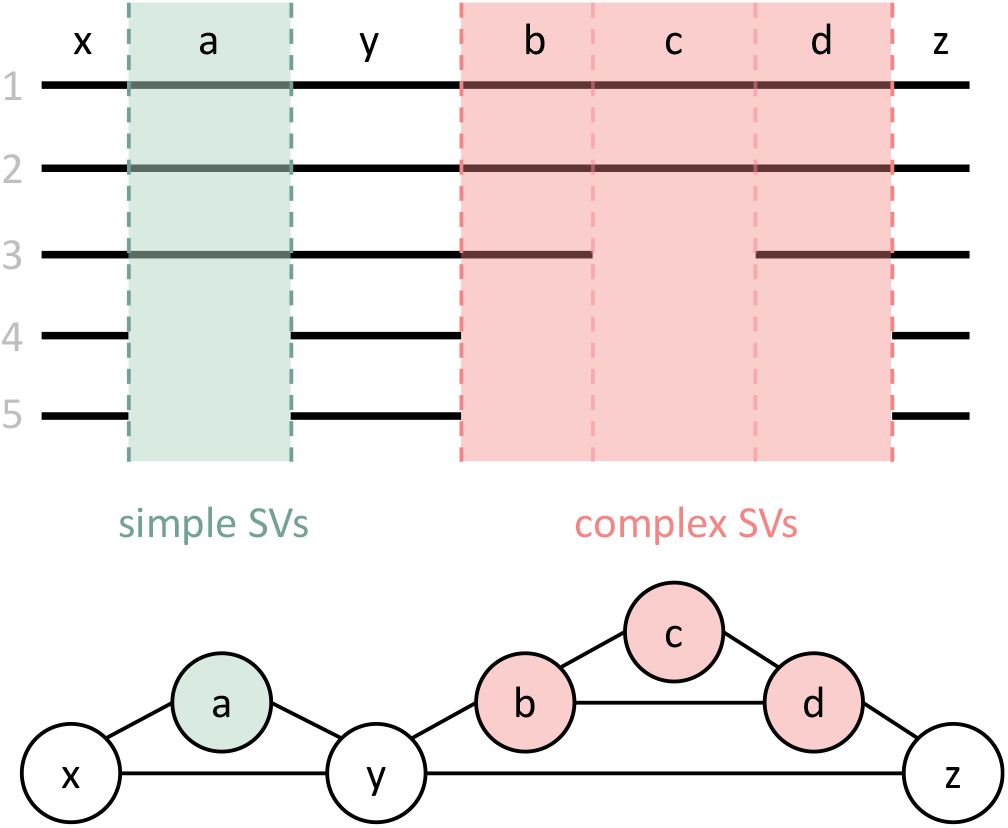
Simple and complex length variants. Cartoons illustrating our classification of length variants into simple and complex structural variants (SVs) in the whole-genome alignment and graph representations.

**Extended Data Figure 11.**
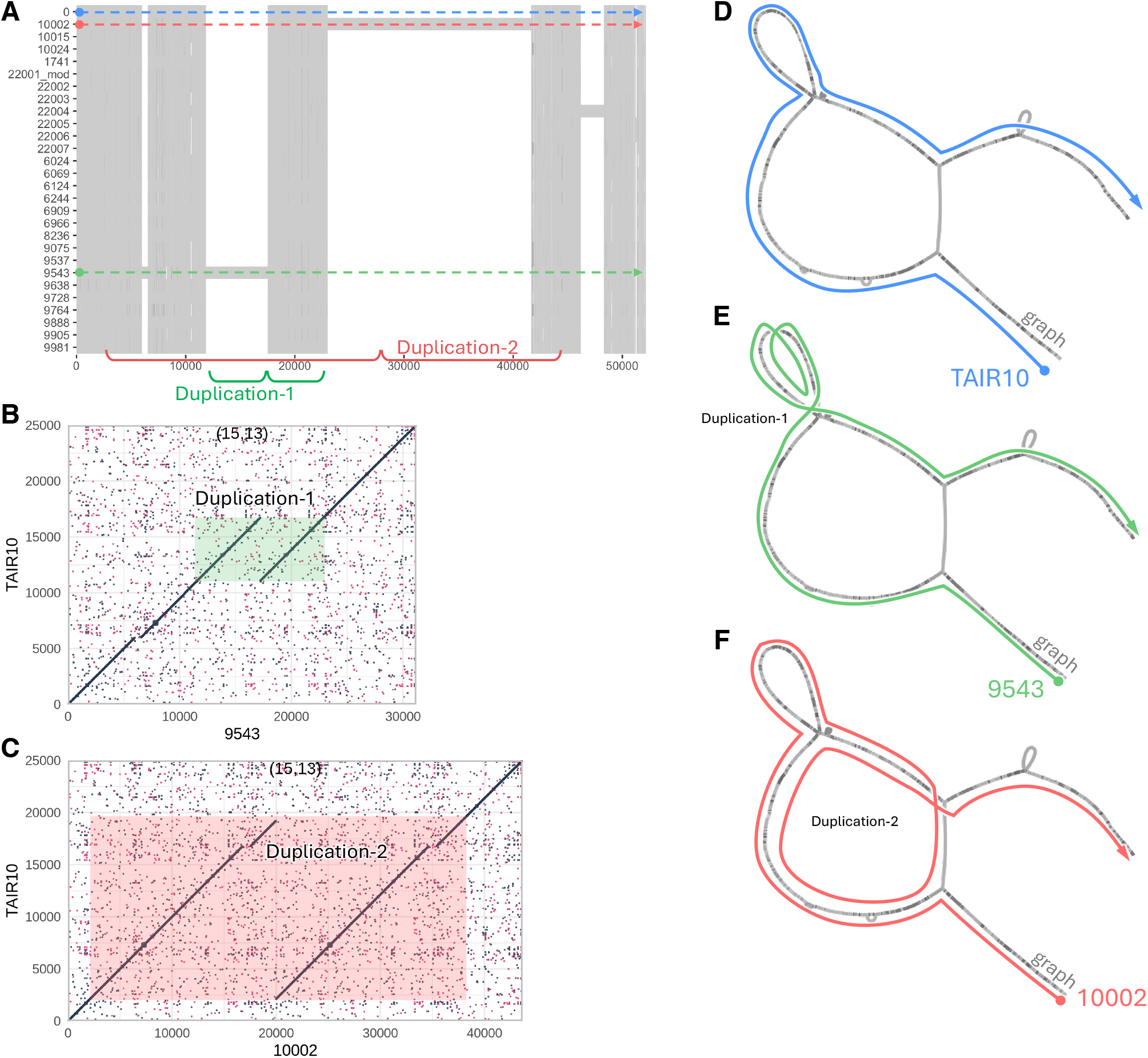
An example of how Pannagram and the PGGB graph each handle closely linked duplications. The region depicted corresponds to coordinates 285,000-310,000 bp on chromosome 1 in accession 1741. The Pannagram alignment (panel **A**) identifies four simple SVs, with the two longest ones being due to duplicated sequences in accession 9543 (panel **B**) and accession 10002 (panel **C**). The PGGB graph representation of this region is shown on the right, along with paths corresponding to three different haplotypes. Panel **D** shows the path taken by accession 0 (TAIR10), which carries the majority haplotype. Panel **E** shows the path of accession 9543, which carries a duplication, and hence goes around the small loop twice. Panel **F** shows the path of accession 10002, which has the longest duplication, and hence goes around the big central loop twice. Thus, while Pannagram identifies four simple SVs (the longest one being 18.6 kb long), the PGGB graph SVs involve all accessions and cover almost the entire region shown. Note that, as in the cartoon example (Extended Data Fig. 9), similar PGGB graph topologies may result from very different types of sequence differences between accessions.

**Extended Data Figure 12.**
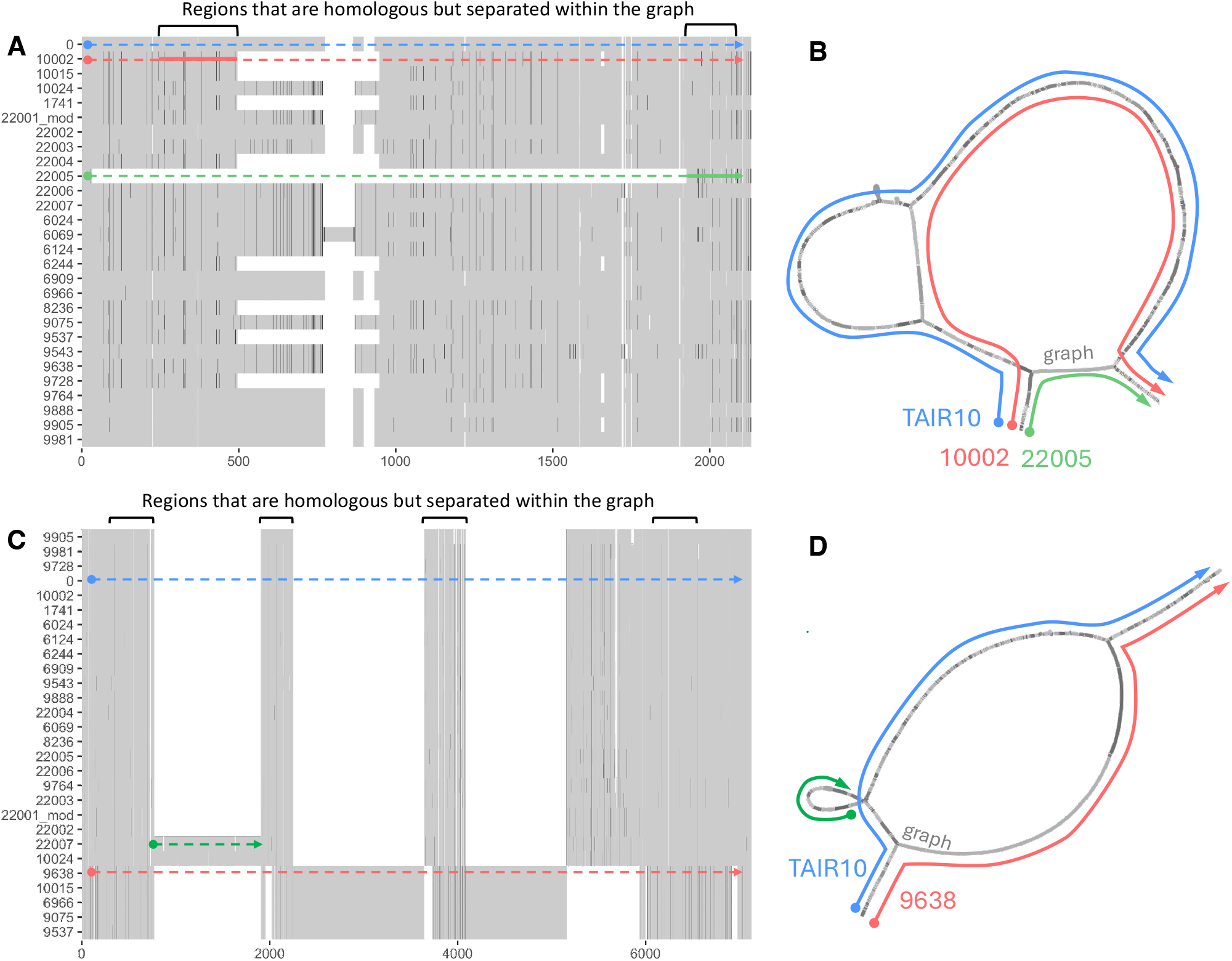
Another example of how Pannagram and the PGGB graph each handle regions that are difficult to align. **A**. Pannagram alignment of a highly polymorphic region corresponding to coordinates 21,956,400-21,958,060 bp on chromosome 1 in accession 1741. Pannagram identifies a complex SV covering most of the region. **B**. The PGGB graph also recognizes these SVs, but merges them with flanking SNP variation, resulting in two nested hat-like structures (*cf*. Extended Data Fig. 9B). As a result, the sequence covered by SVs is longer. **C**. Pannagram alignment of the region corresponding to coordinates 1,183,130-1,186,590 bp on chromosome 1 in accession 0 (TAIR10). Pannagram identifies several, mostly simple SVs separated by short alignable regions. **D**. The PGGB graph does not align these regions, and merges most variants into two longer haplotypes. In this case as well, the graph SVs cover more sequence than the Pannagram SVs.

**Extended Data Figure 13.**
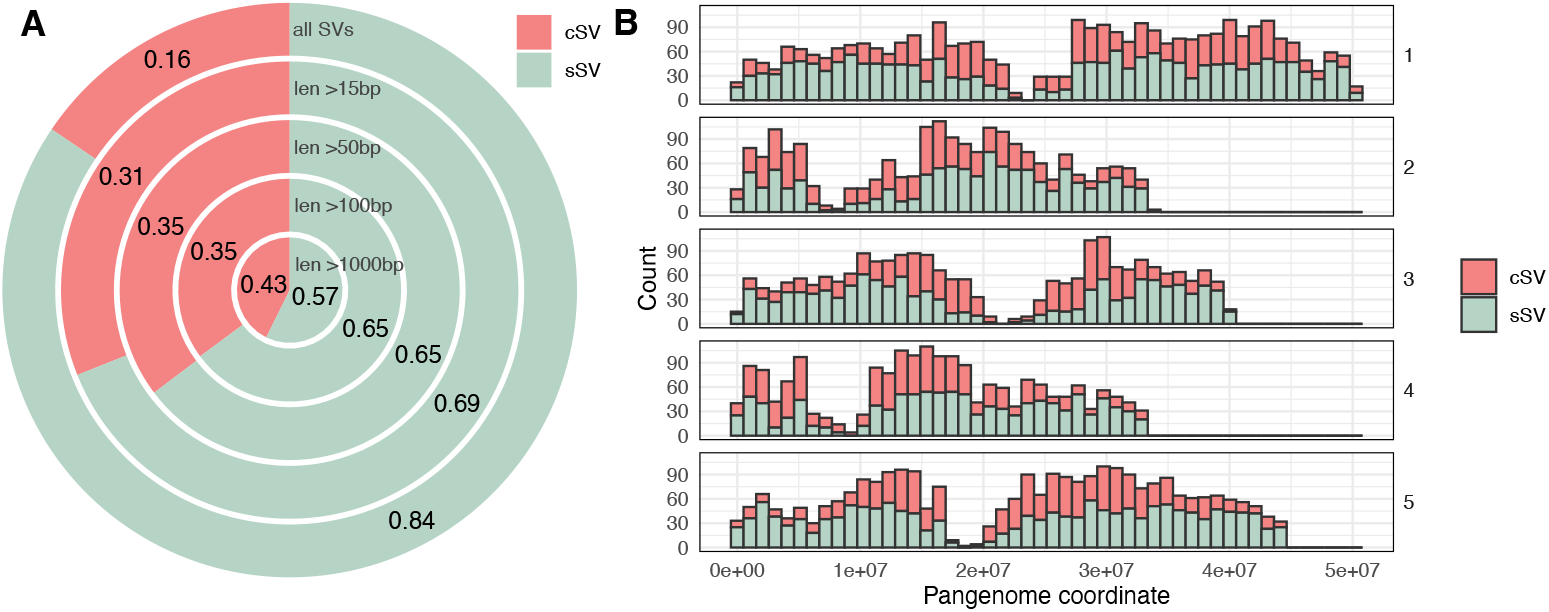
Complex versus simple SVs. **A**. Fraction of simple SVs (sSVs) and complex SVs (cSVs) by length. The prevalence of short sSVs is at least partly explained by the low probability of the multiple events required for cSVs occurring in small DNA segments. **B**. The distribution of sSVs and cSVs across chromosomes—the latter are enriched in pericentromeric regions.

**Extended Data Figure 14.**
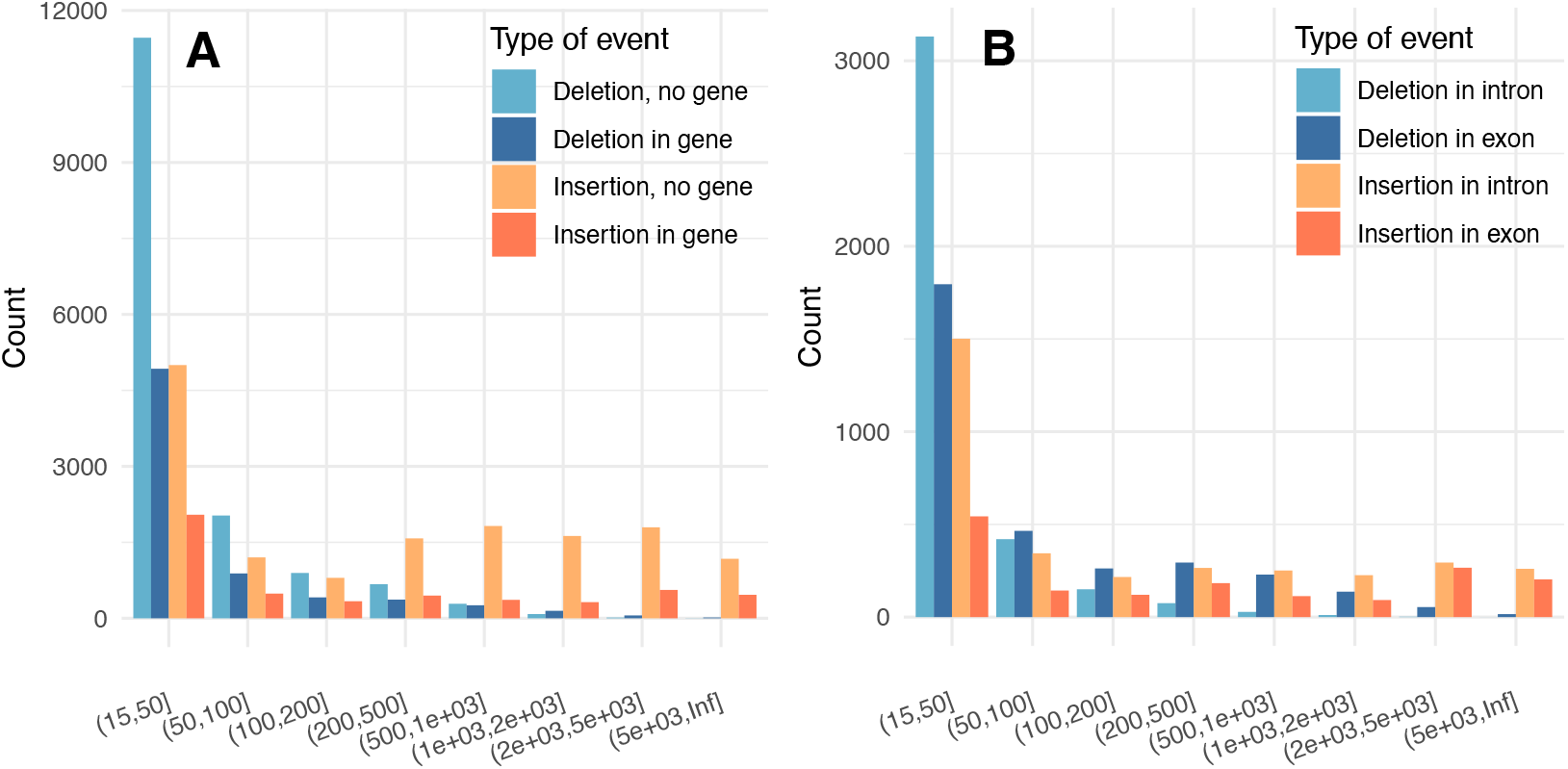
Location of sSVs with respect to genes. **A**. The number of putative insertions (presence allele in 1-3 accessions) and deletions (absence allele in 1-3 accessions) of varying lengths in gene and intergenic regions. **B**. Same for exons and introns.

**Extended Data Figure 15.**
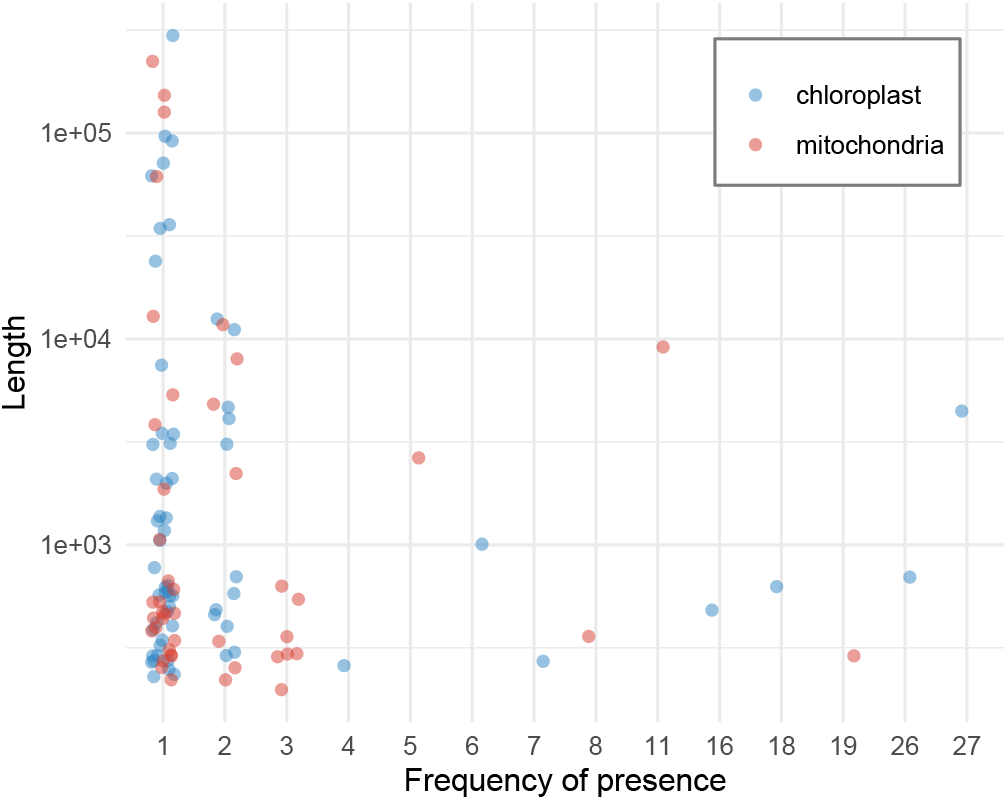
Frequency and size distribution of organellar insertions. Seven insertions are excluded because they were associated with contig breaks, and we could therefore not determine their exact length (Extended Data Fig. 2).

**Extended Data Figure 16.**
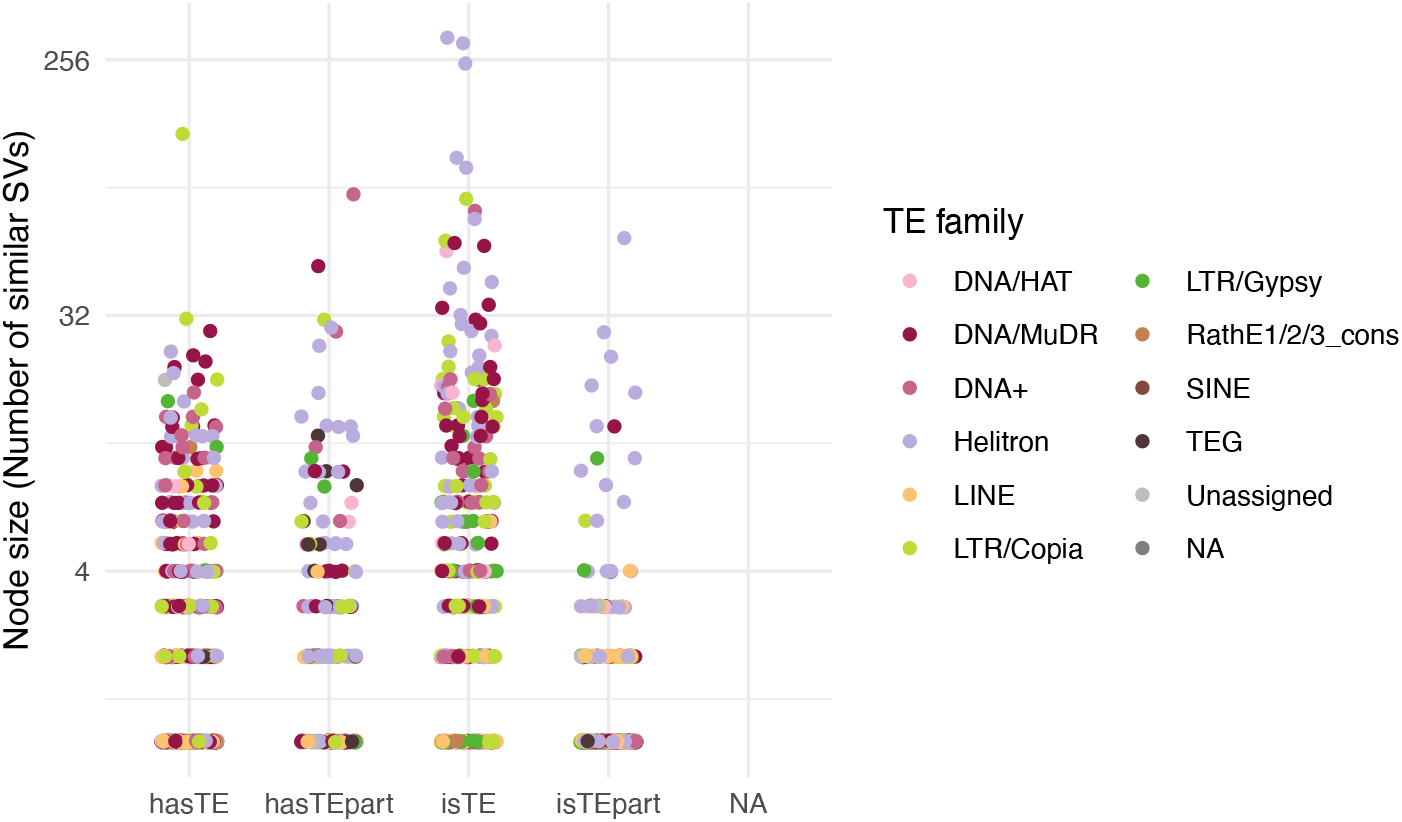
Size-distribution of graph nodes by TE-content. Large nodes are found not only for presence alleles that correspond to complete annotated TE, but also for other categories, demonstrating that these are also part of the mobile-ome.

**Extended Data Figure 17.**
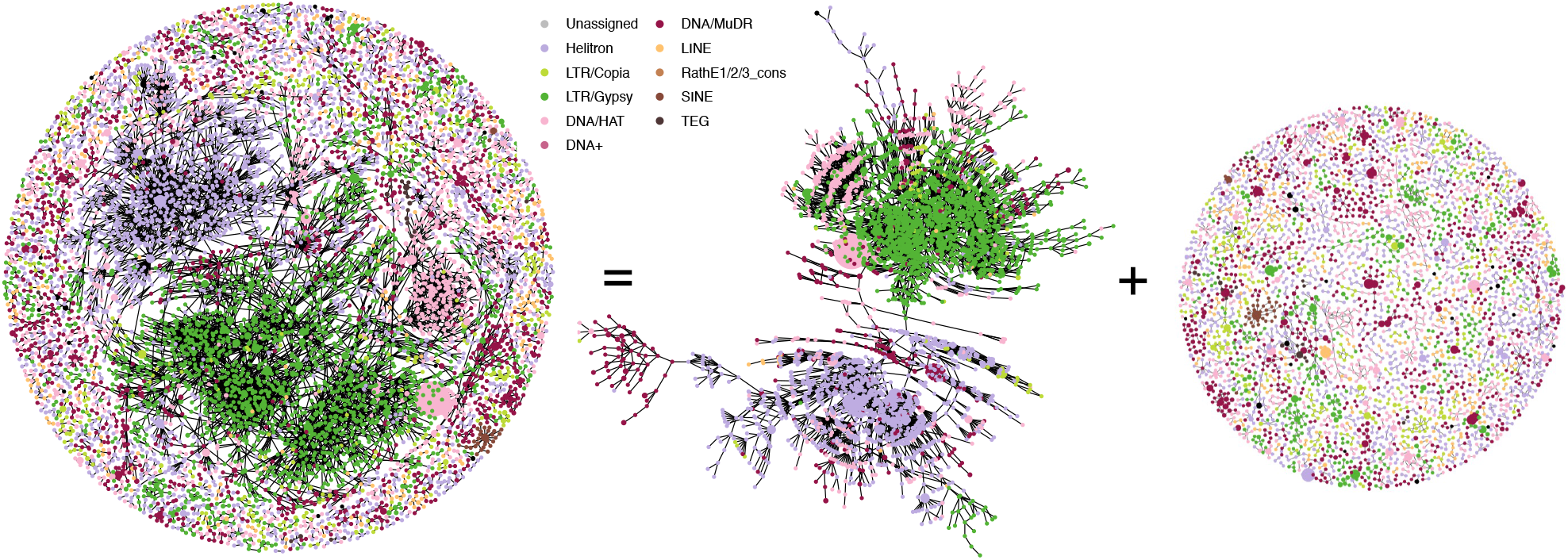
The graph of the nestedness of existing *A. thaliana* TE annotations. Each node is a cluster of similar sequences (with length and identity thresholds of 0.85), where the size of the node indicates its relative abundance. The graph can be decomposed into one dominant connected component and several smaller ones, as shown on the right.

**Extended Data Figure 18.**
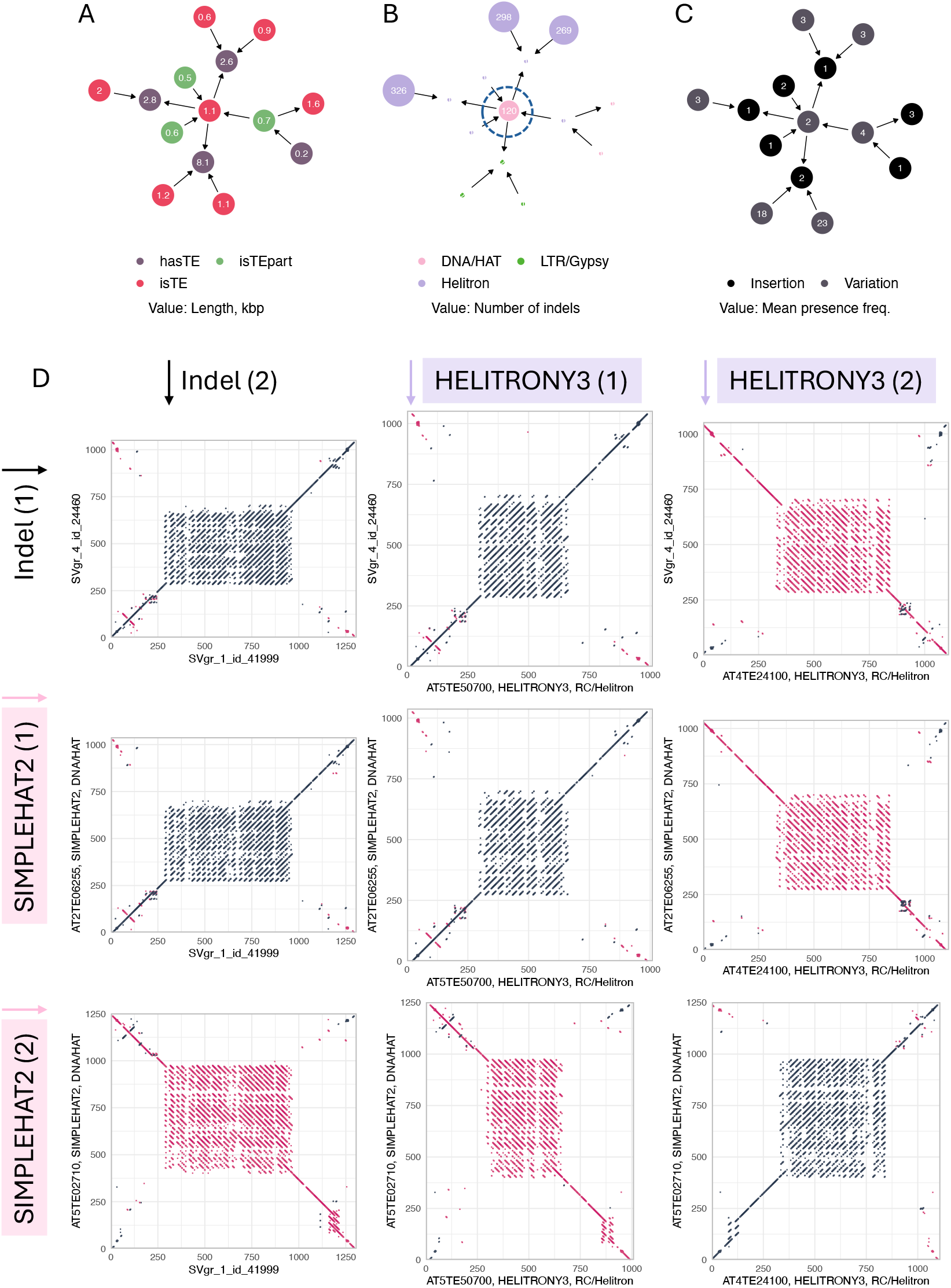
Different TE annotations of Very similar mobile elements. **A-C**. A particular part of our graph of nestedness (Fig. 5) colored separately by TE content, TE superfamily, and presence frequency. **D**. Dot plots comparing sSVs from the central node of the graph. Dark color reflects the similarity on the forward strand, pink color on the reverse complement. The dot plots were constructed with a window parameter of 15 and the number of matches set to 12. Sequences in the central node are very similar to members of both HELITRONY3 and SIMPLEHAT2 families, which are also similar to each other, demonstrating how confusing TAIR10 TE annotation leads to nodes connecting different TE superfamilies.

**Extended Data Figure 19.**
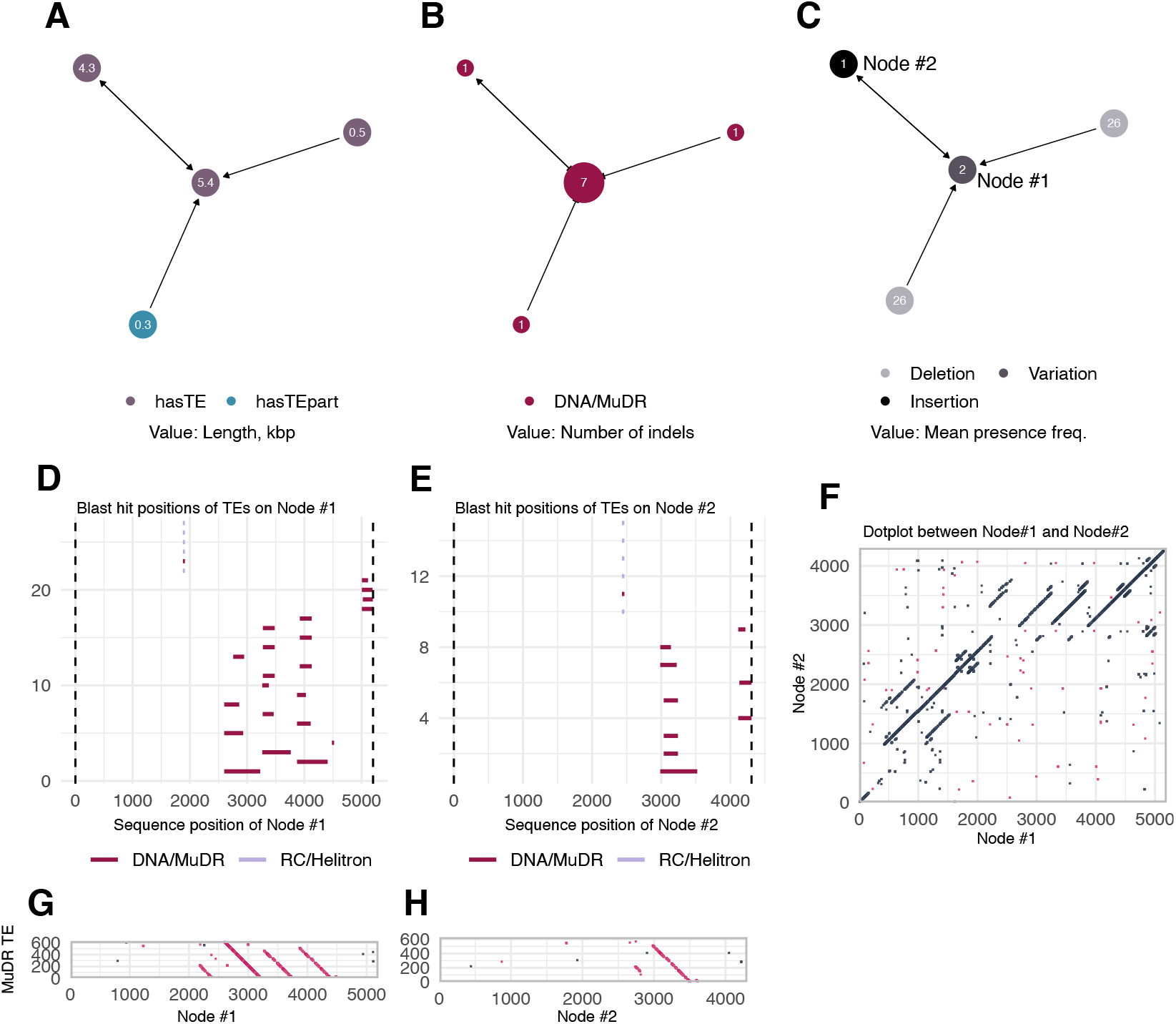
An example of a mobile element containing annotated TE sequences. **A-C**. A connected component from the graph of nestedness (Fig. 5) colored separately by TE content, TE superfamily, and frequency (indicating likely insertion/deletion status). The component is characterized by putative insertions of a large element (5.4 kb) containing annotated DNA/MuDR element. As illustrated by BLAST results (**D-E**) and dot plots (**F-H**), the Node #1 presence alleles contains several copies of an annotated MuDR element, while the Node #2 allele contains one copy. The nature and mechanism of transposition of this mobile element is unclear.

**Extended Data Figure 20.**
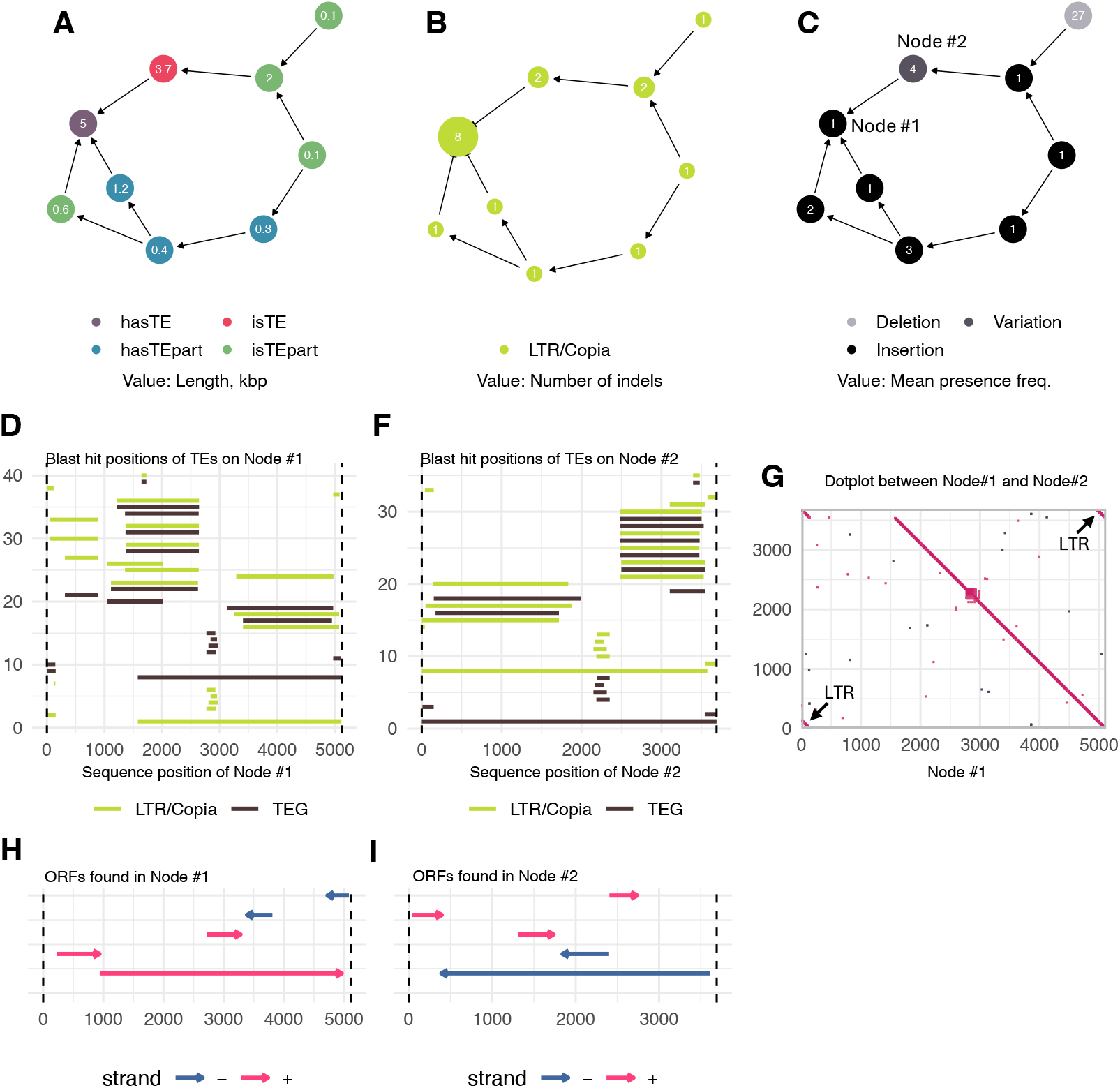
Another example of a mobile element containing annotated TE sequences. **A-C**. A connected component from the graph of nestedness (Fig. 5) separately colored by TE content, TE superfamily, and frequency (indicating likely insertion/deletion status). The component is characterized two active elements, a 3.7 kb one (Node #2) corresponding to an annotated LTR/Copia element, and an apparently more active one (based on copy number) that is larger (5 kb, Node #1) and contains a very similar LTR/Copia element plus additional sequence, including LTR/Copia fragments. Both elements have (matching) LTRs (**G**), as well as LTR/Copia ORFs. Unlike the example in Extended Data Fig. 19, the nature and mechanism of this element is clearer, and we have previously described an LTR/Copia element that is longer than the existing TAIR10 annotation suggests ^57^.

**Extended Data Figure 21.**
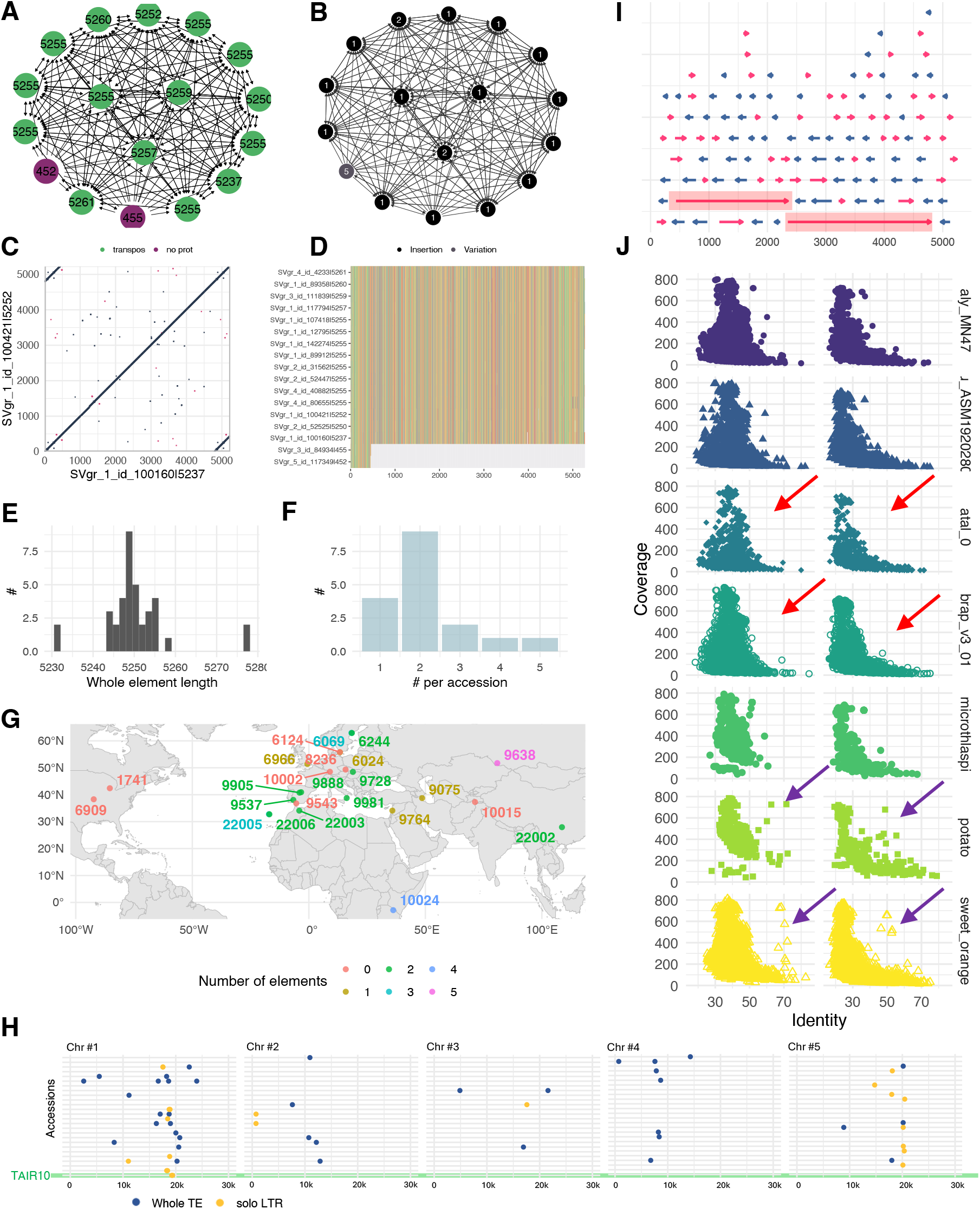
An un-annotated TE with evidence of putative horizontal transfer. A component from the graph of nestedness (Fig. 5) built on sSVs that have no overlap with annotated TEs in *A. thaliana*. **A-B**. The component consists of many rare insertions of a ∼ 5.3 kb element containing an ORF matching “transpos*” in protein BLAST, plus a few more common insertions of a ∼450 bp element without coding potential. A dot plot (**C**) and multiple alignment of all alleles from the graph (**D**) reveal the presence of LTRs, and identify the ∼ 450 bp element as a solo-LTR. The length of the element (**E**) is highly conserved, and it is mostly present in low copy number (**F-G**). The element is most likely un-annotated in TAIR10 because it is not present in the TAIR10 reference genome—only a solo-LTR is found in TAIR10 (**H**, which contains all detected instances, not only those identified as sSVs). The element contains two long putative ORFs (**I**) without matches in genomes of closely related species, but possible matches in potato and sweet orange (**J**).

**Extended Data Figure 22.**
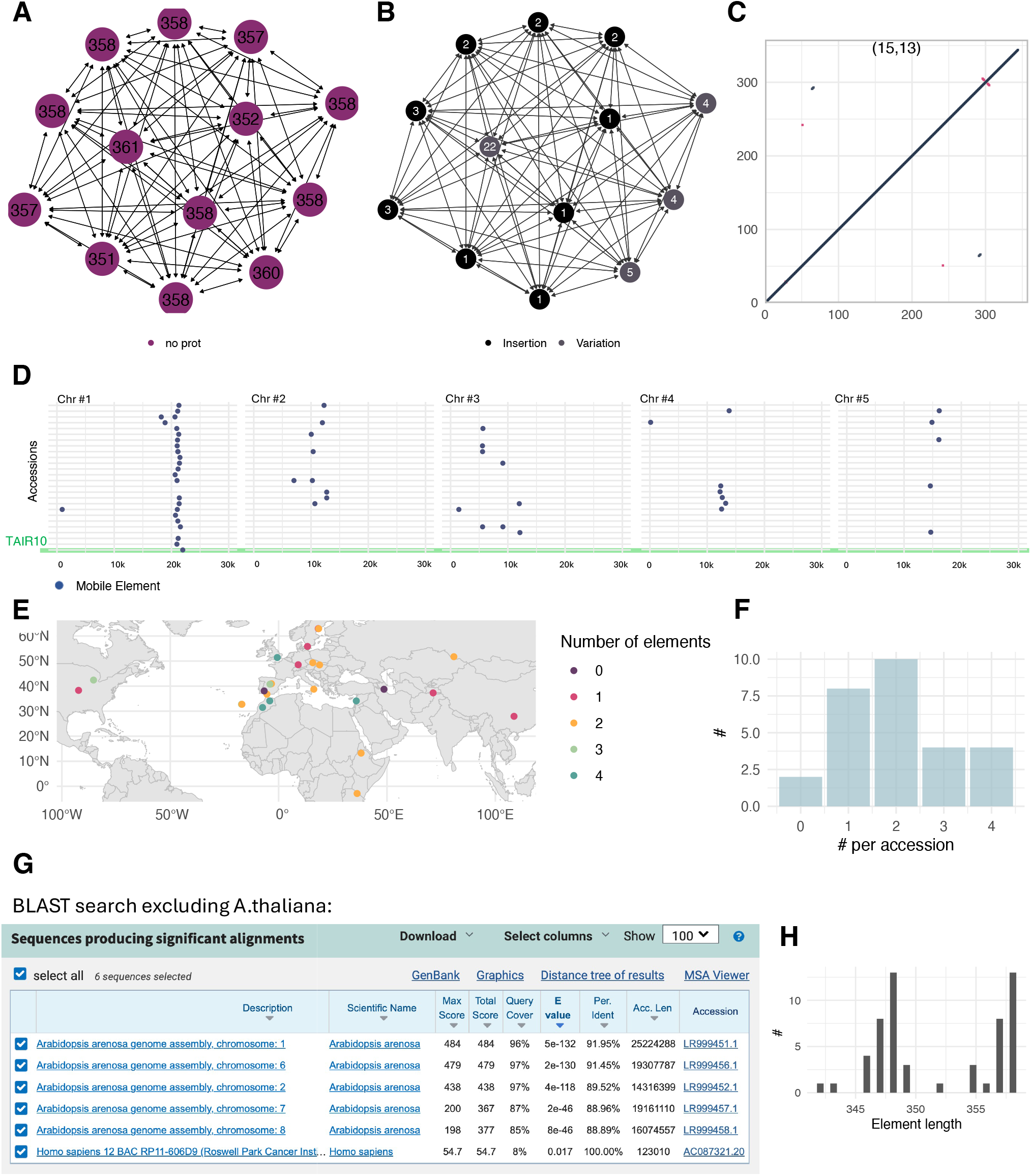
A putative novel mobile element family. A connected component from the graph of nestedness (Fig. 5) built on sSVs that have no overlap with annotated TEs. **A-B**. The component consists of many variable-frequency insertions of a short sequence of 351-361 bp without coding potential. The element is not repetitive (**C**) but is surrounded by putative target-site duplications (not shown). The element is low copy-number, but insertions are sometimes common (**D-F**; D contains all detected instances, not only those in sSVs). An NCBI BLAST search excluding *A. thaliana* (**G**) identified no matches except in the closely related *A. arenosa*. **H**. The length distribution of the element, again including all detected instances, not only those in sSVs.

**Extended Data Figure 23.**
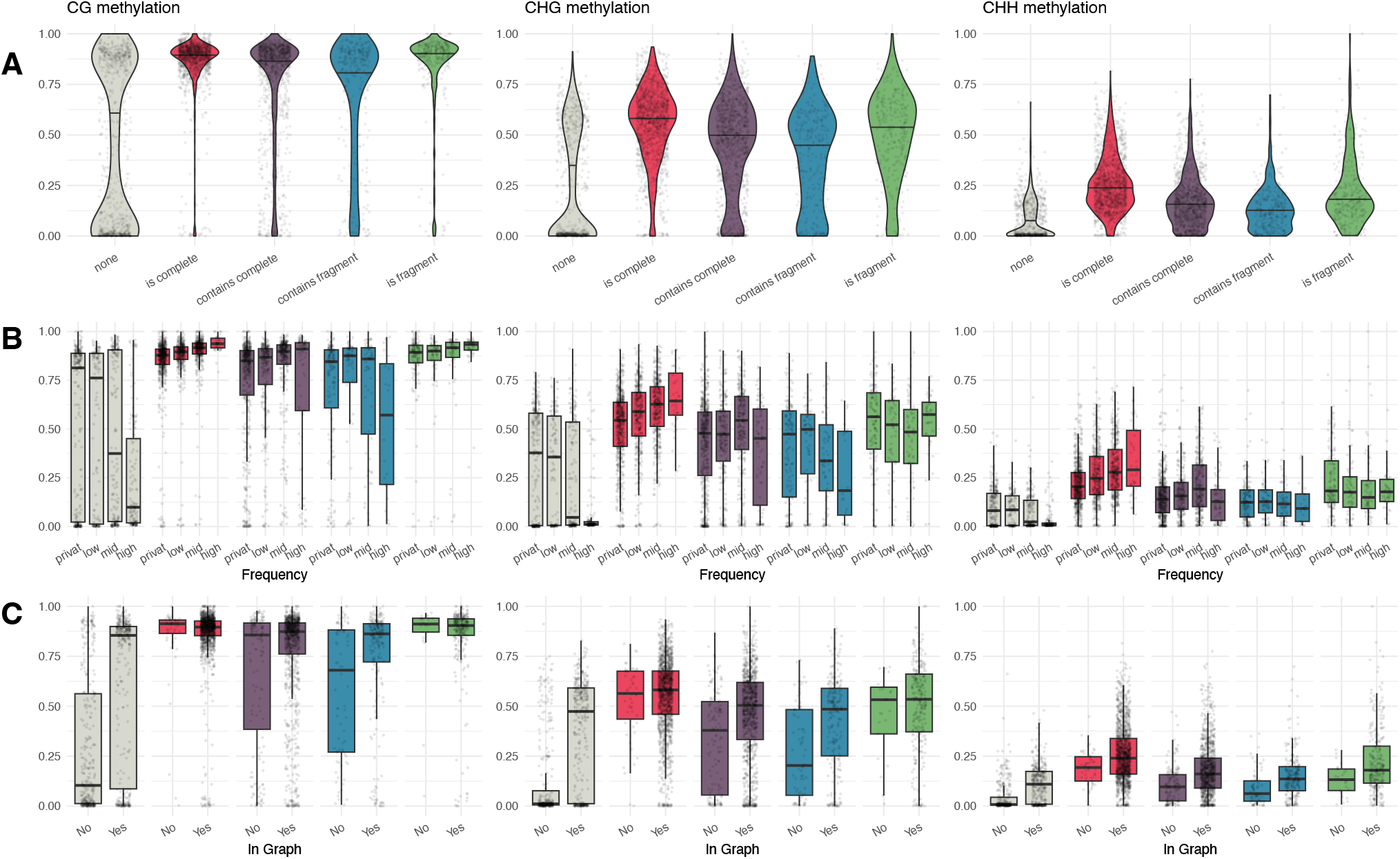
Methylation of sSVs. **A**. Methylation levels of sSVs categorized by their TE content. Those corresponding to complete TEs or TE fragments are most highly methylated. **B**. Methylation levels of sSVs categorized by their TE content and frequency of presence (“private” = 1, “low” = 2–3, “mid” = 4–24, “high” = 25–26). For those corresponding to complete TEs, methylation increases with frequency (which is correlated with age). **C**. Methylation levels of sSVs categorized by membership in the graph of nestedness (Fig. 5). Methylation levels are always higher for sSVs in the graph, and the difference for those without annotated TE content is striking. Methylation on SVs was estimated using the BS-seq data from ^62^ 12 accessions mapped to their corresponding genomes (Methods). Only simple SVs are plotted. Each data point in the plots corresponds to the maximal methylation level of each sSV among the accessions where this sSV is present.

**Extended Data Figure 24.**
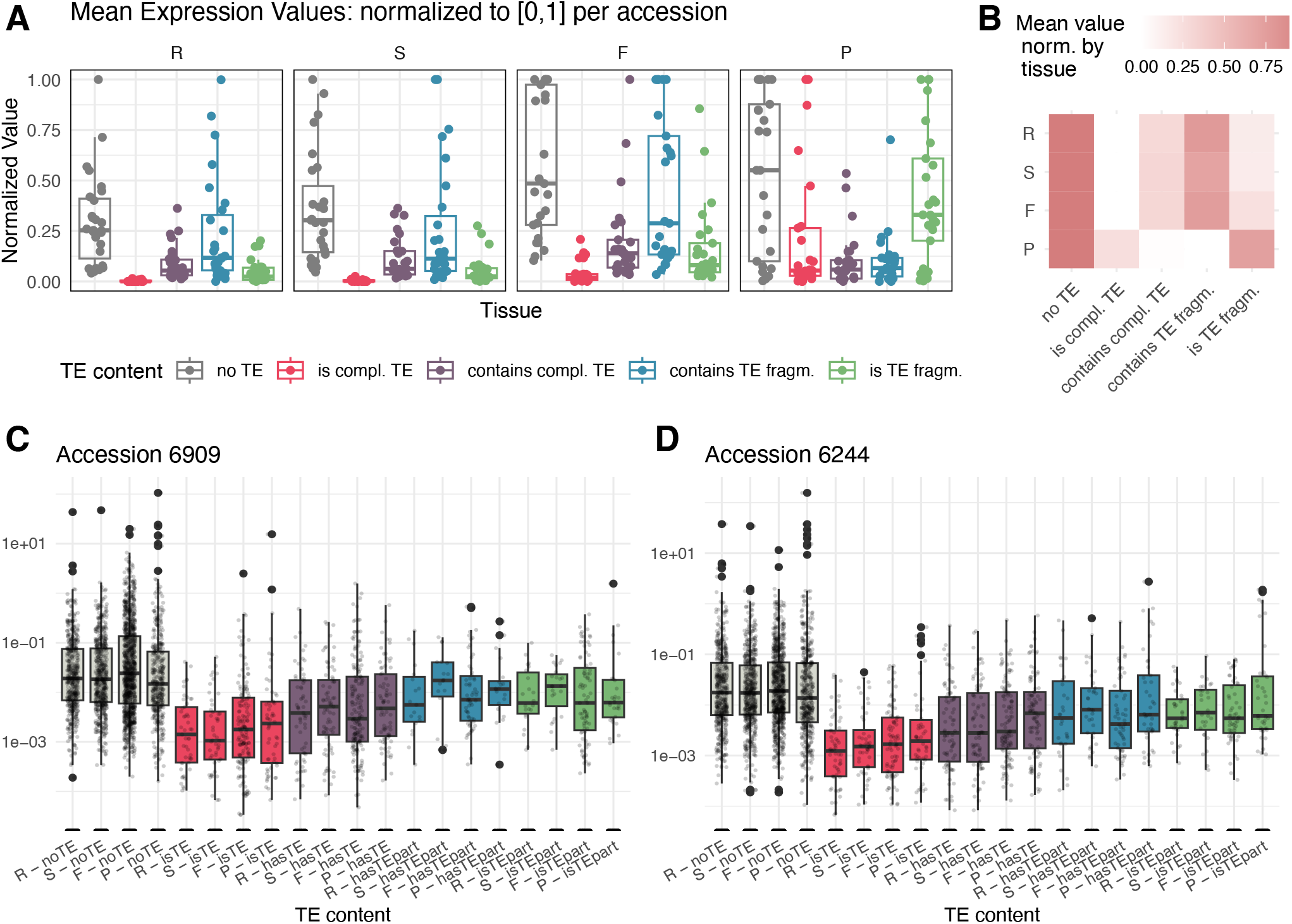
Expression of sSV sequences. The distribution of expression levels as a function of different TE-coverage categories and tissues (R = rosette; S = seedling; F = flower; P = pollen), illustrating that sequences in sSVs that are more likely to be part of the mobile-ome have low expression. For each sSV, read counts were normalized by length. **A**. Each dot is the mean expression of sequences in sSVs in that category for an accession, using values normalized within accessions. **B**. Mean expression of sequences in sSVs for each category normalized within tissue. **C–D**. For two different accessions, the distribution of expression for each category and tissue. Values have been normalized within tissue.

**Extended Data Figure 25.**
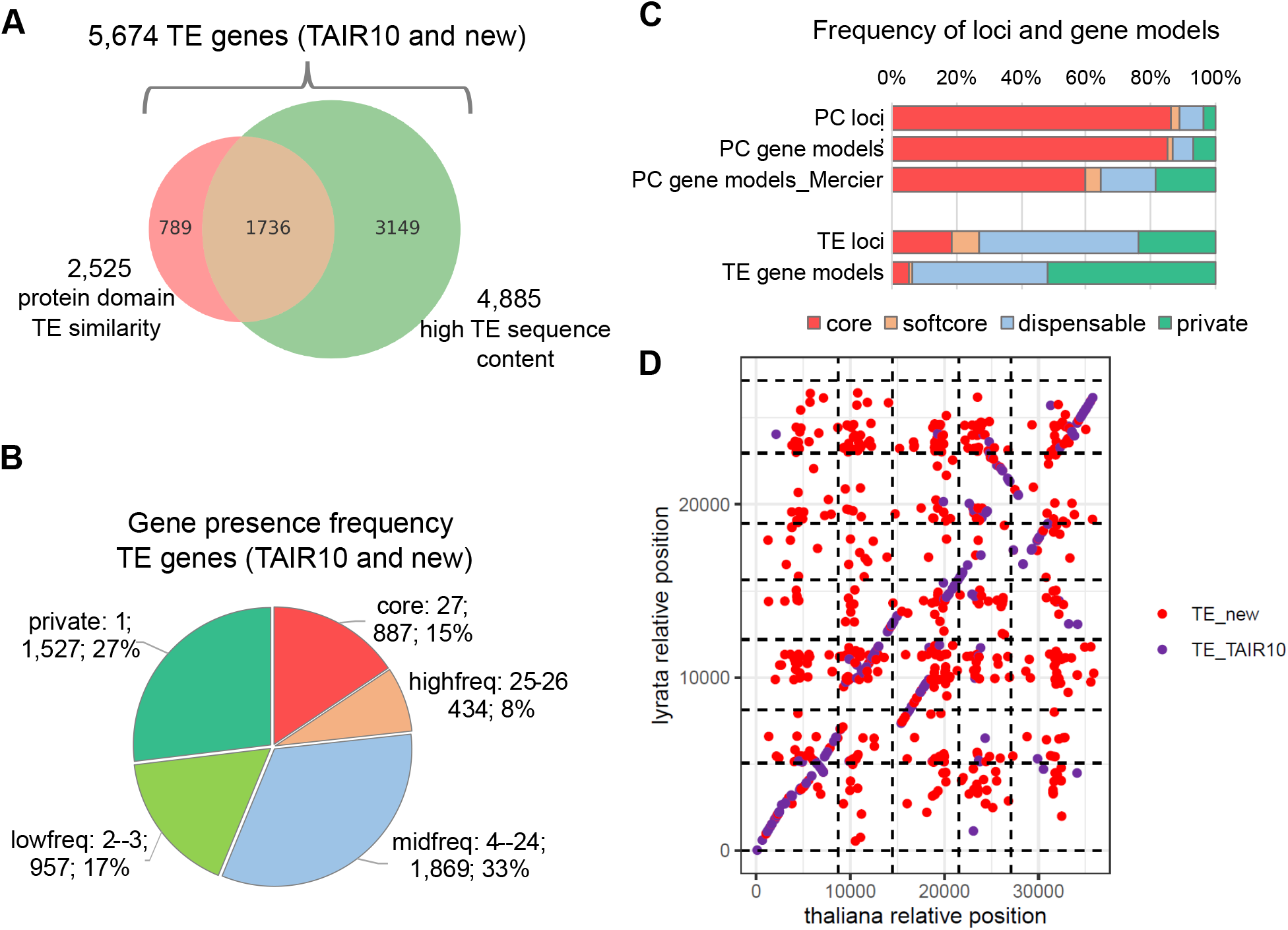
Details of gene-ome analysis. **A**. Overlap between two indicators used to identify TE: a) high nucleotide similarity (locus contains > 50% annotated TE sequence in at least one accession) and; b) protein domain similarity (ORF having similarity to proteins known to be involved in TE function). We defined TE genes as the union of these sets. **B**. Gene presence frequency distribution for TE genes. **C**. Comparison of gene presence variation in this study to a recent analysis of gene model in 69 *A. thaliana* accessions ^23^. For both protein-coding genes and TE genes, presence-absence variability is higher for gene models compared to gene loci. This can be explained by the fact that our gene annotation pipeline might miss lowly expressed genes or genes not expressed in any of the four tissues we used. The gene model variability in the protein-coding gene models of Lian *et al*. ^23^ is noticeably higher than in our data. This could reflect that the larger population size in their study, but the different approaches used for matching genes between accessions and estimating gene model variation: while we used pan-genome coordinates, they used an orthogroup search approach. For example, if the gene locus is present in every accession and has a gene model in every accession, but in three accessions a shorter gene model was annotated, the orthogroup approach would produce two gene families for this locus and call both gene models dispensable, while our approach would treat this locus as one and would identify such a gene (and gene models corresponding to this gene) as part of the core, since it is represented by a gene model in every accession. **D**. Synteny and sequence similarity analysis in *A. lyrata* for TAIR10 and new TE genes with the same stringency as in Fig. 6: 80% sequence similarity (see Methods). Genes are plotted by their gene IDs along the chromosomes. The small number of data points compared to protein-coding genes is due to the fact that most TE genes were not found in the *A. lyrata* genome.

**Extended Data Figure 26.**
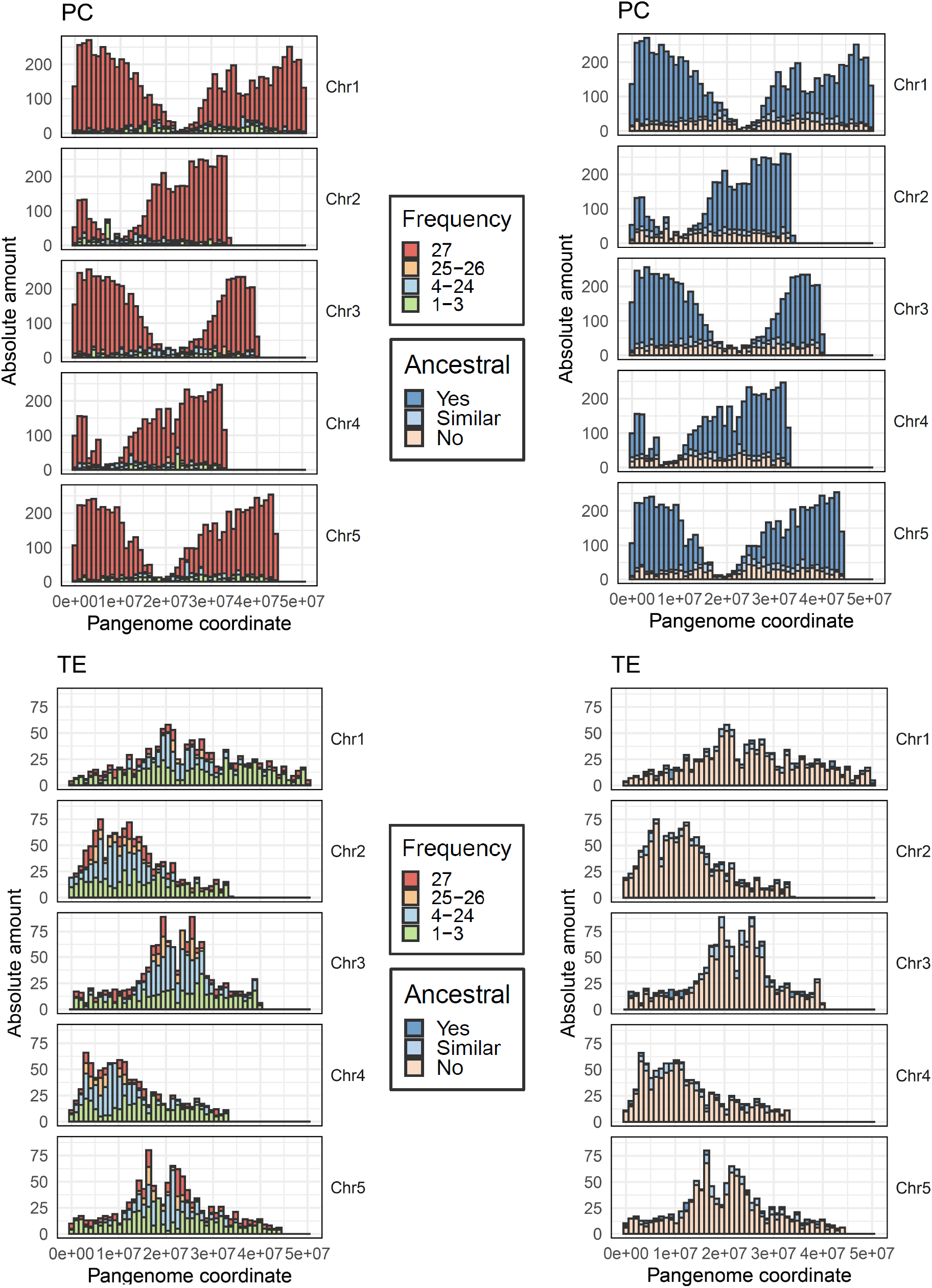
Chromosomal distribution of genes and TEs. The distribution of protein-coding genes and TEs along all five chromosomes is shown broken down by frequence of presence and ancestral status. *Cf*. Fig. 6.

**Extended Data Figure 27.**
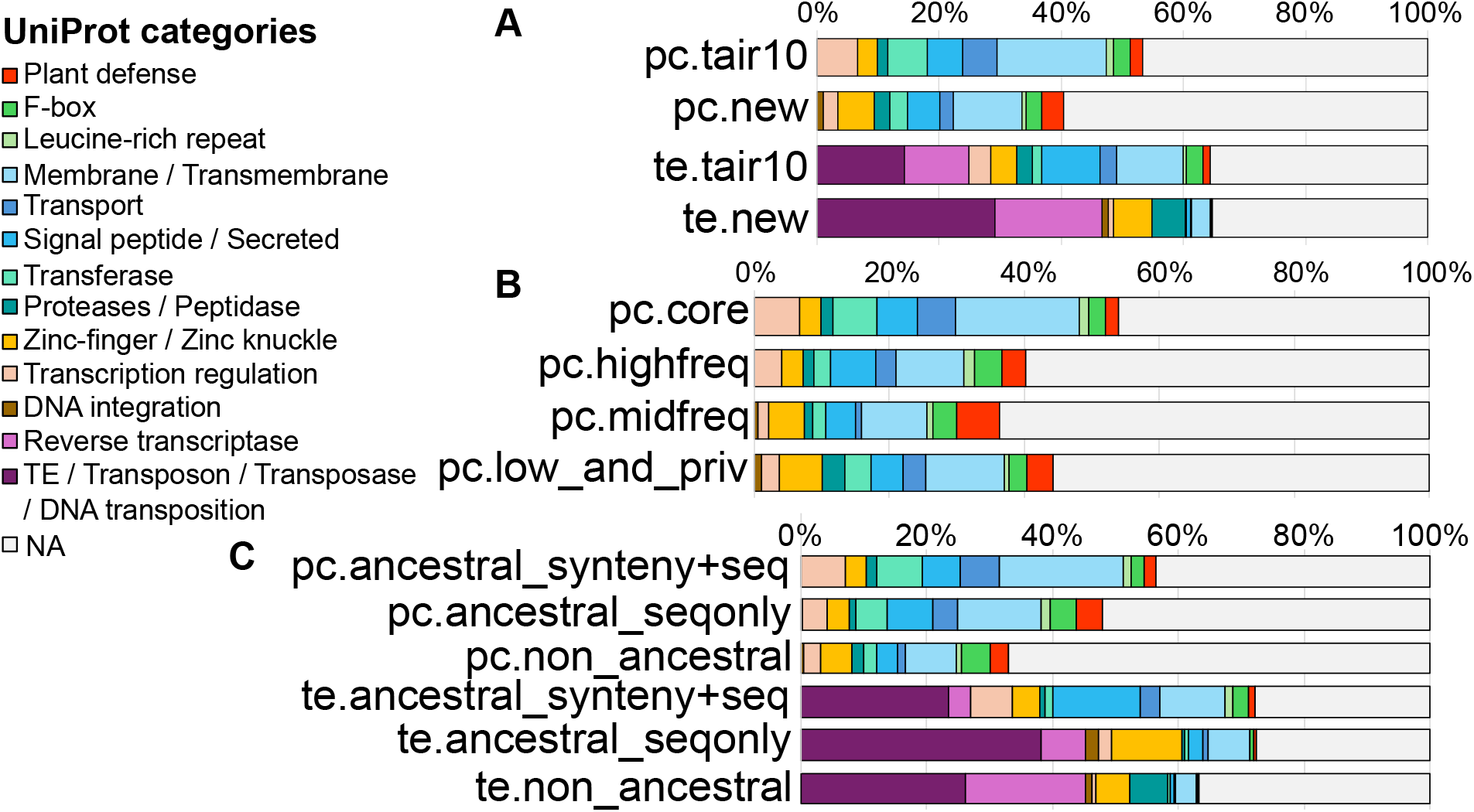
Additional functional domain gene distributions. Functional domains identified by UniProt (see Methods). “NA” indicates genes where no functional domains could be found. **A**. Genes grouped by new vs. old and protein-coding vs. TE. **B**. Protein-coding genes grouped by frequency of presence. **C**. Protein-coding and TE genes by ancestral status.

**Extended Data Figure 28.**
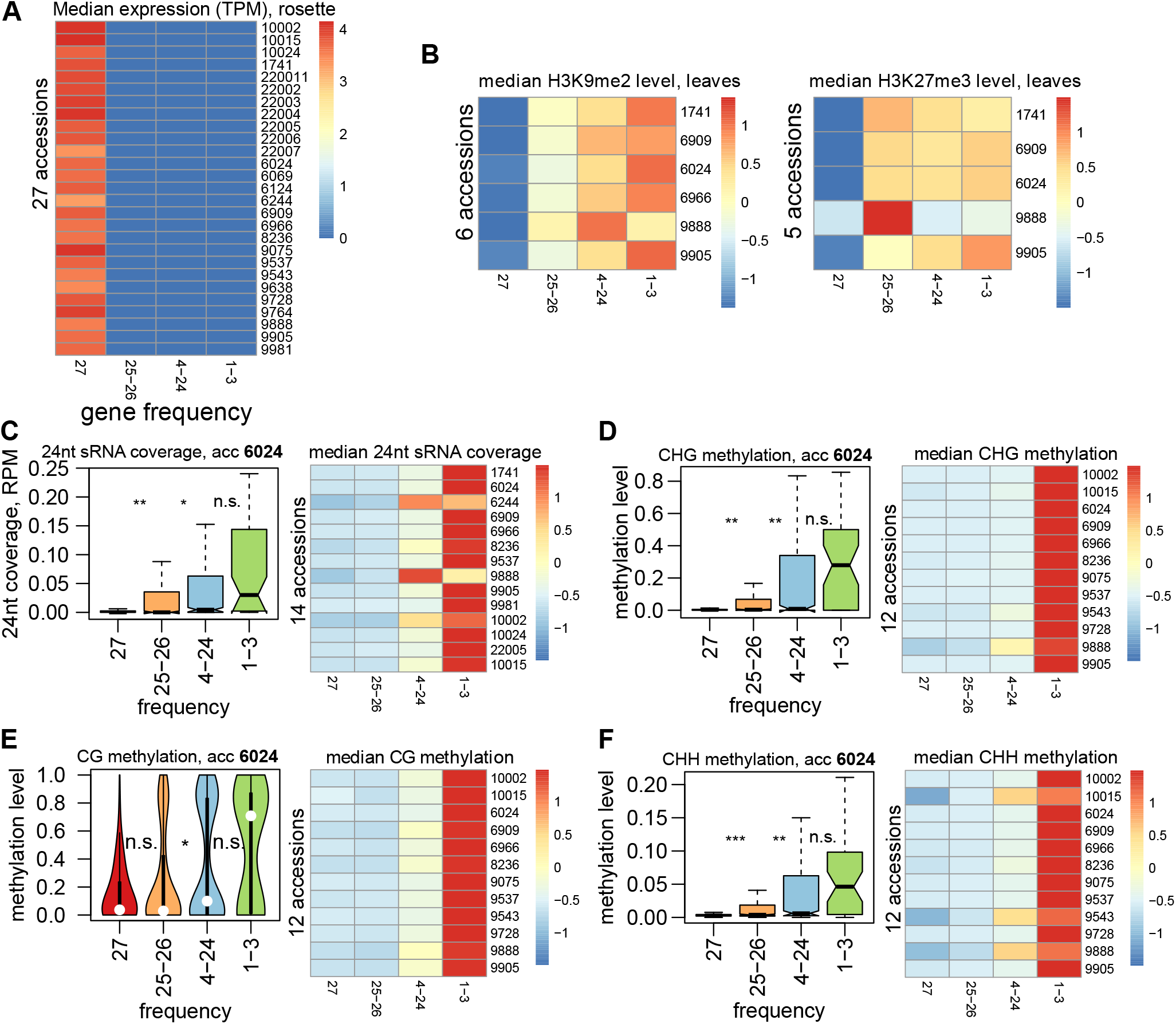
Protein-coding gene silencing by frequency category. **A**. Expression of genes vs. their presence frequency. The median TPM for each gene category is plotted for each accession. **B**. H3K9me2 and H3K27me3^63^ of genes vs. their presence frequency. The median of quantile- and input-normalized ChIP-seq signals for each gene category is plotted for each accession. **C**. 24-nt sRNAs targeting of genes of different presence frequency. Left: 24-nt locus coverage in flowers of accession 6024. Right: the median of 24-nt coverage is plotted for each gene category and accession. **D–F**. Locus-wide CHG, CG and CHH methylation levels in leaves ^62^ for genes of different presence frequency. Left: accession 6024. Right: the median of CHH methylation level is plotted for each gene category and accession. All heatmaps were created using the heatmap package in R and all except for expression are scaled by row to facilitate within- and between-accession comparison. Significance estimates are from Mann-Whitney tests (^***^: *P* < 10^−10, **^: *P* < 10^−5, *^: *P* < 10^−2^, n.s.: *P* ≥ 10^−2^).

**Extended Data Figure 29.**
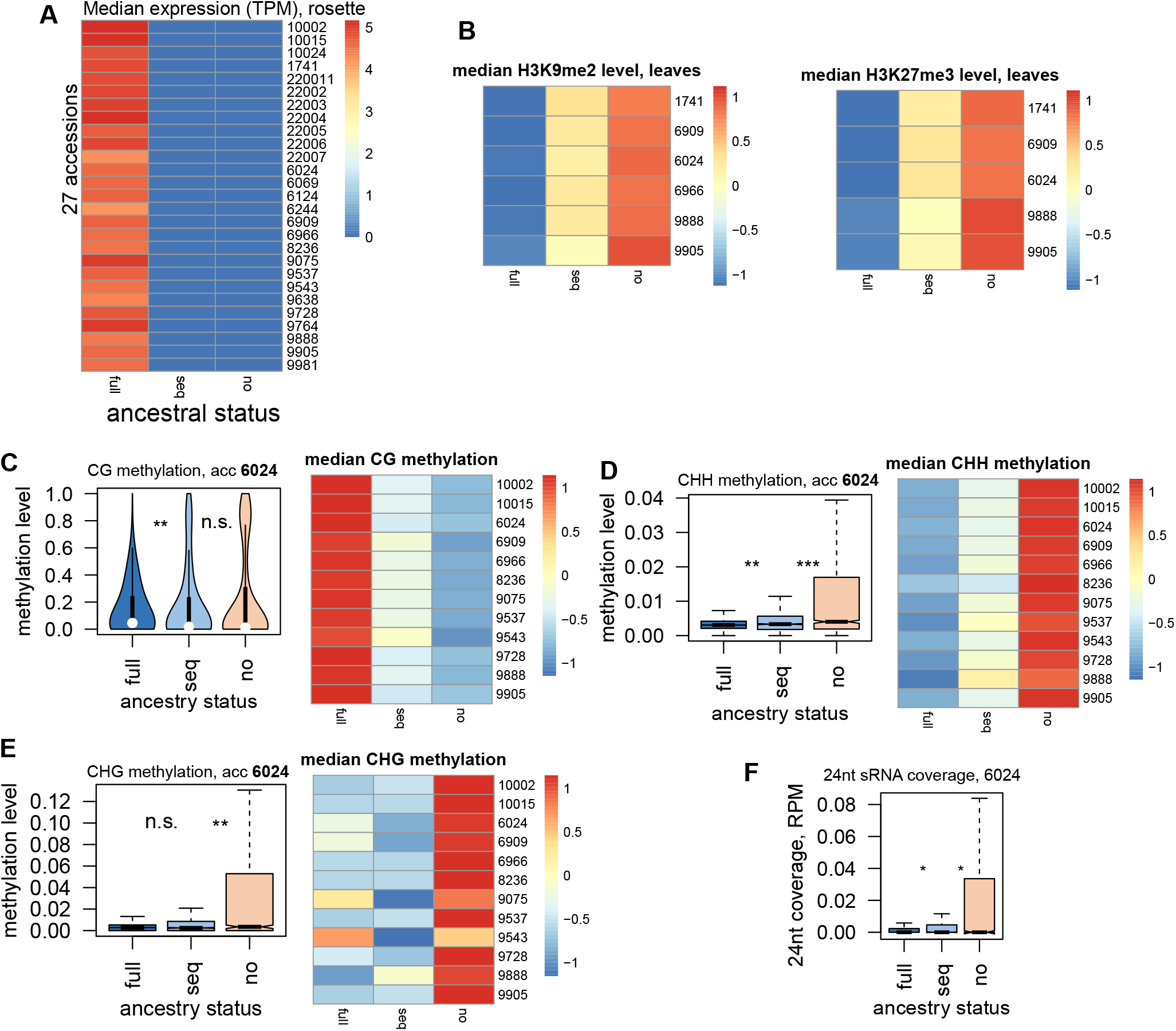
Protein-coding gene silencing by ancestral status. **A**. Expression of genes vs. their ancestral status. The median TPM over the whole gene locus for each gene category is plotted for each accession. **B**. H3K9me2 and H3K27me3 silencing ^63^ of genes vs. their ancestral status. The median of quantile- and input-normalized ChIP-seq signals for each gene category is plotted for each accession. **C-E**. Locus-wide CG, CHH and CHG methylation levels in leaves ^62^ for genes of different ancestral status. Left: accession 6024. Right: the median methylation level is plotted for each gene category and accession. **F**. 24-nt sRNAs locus coverage in flowers of accession 6024 for genes of different ancestral status. The medians heatmap is not plotted due to zero median values in many accessions. All heatmaps were created using heatmap package in R and all except for expression are scaled by row to facilitate within- and between-accession comparison. Significance estimates are from Mann-Whitney tests (^***^: *P* < 10^−10, **^: *P* < 10^−5, *^: *P* < 10^−2^, n.s.: *P* ≥ 10^−2^).

**Extended Data Figure 30.**
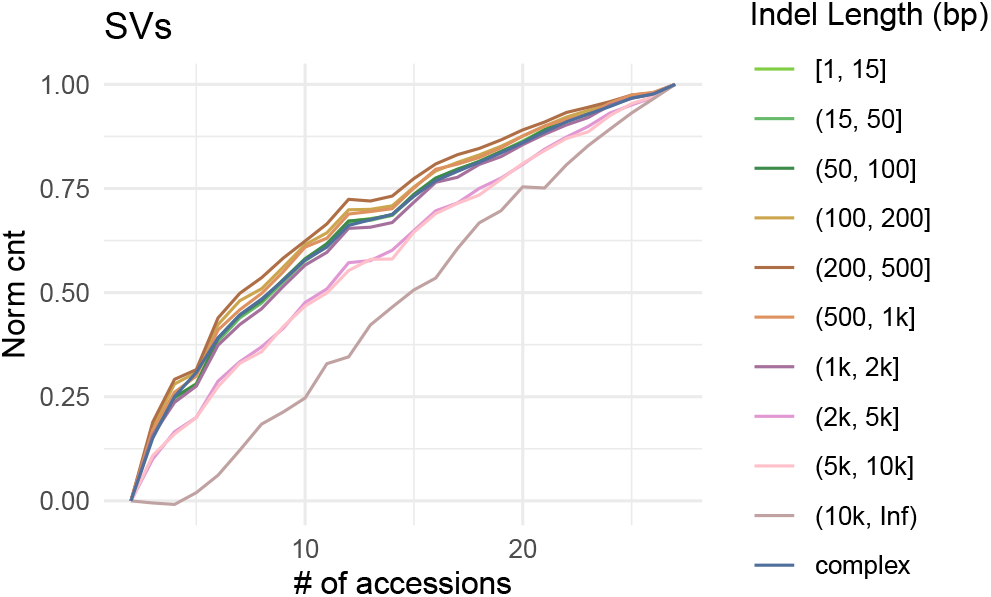
Saturation curves for sSVs for different lengths. Each curve is drawn through points that are averaged values from 20 repetitions. The curves were normalized to start and end at the same points. Larger sSVs saturate more quickly.

**Extended Data Figure 31.**
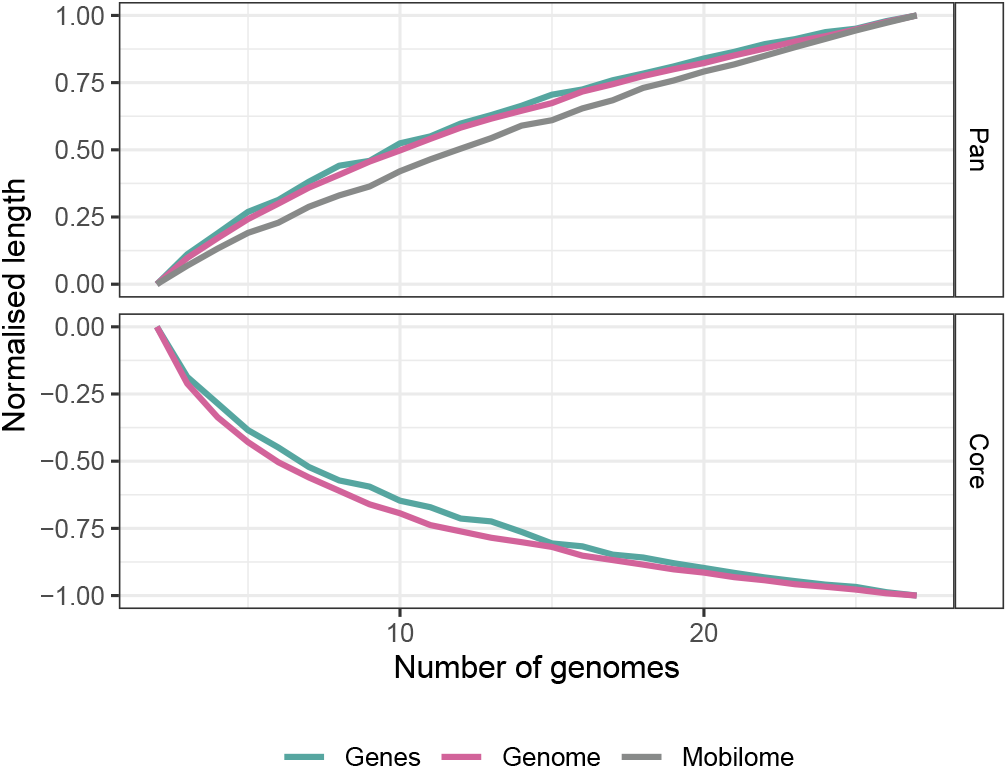
Normalized saturation curves for different genome components. The dependence on sample size for the union (“pan”) and intersection (“core”) sequence length, separately for the full genome, the mobile-ome, and the gene-ome. Within the “pan” curves, the mobile-ome saturates the fastest, the gene-ome the slowest, and the entire genome at an intermediate rate. Gene-ome and whole genome “core” curves show similar trends.

**Extended Data Figure 32.**
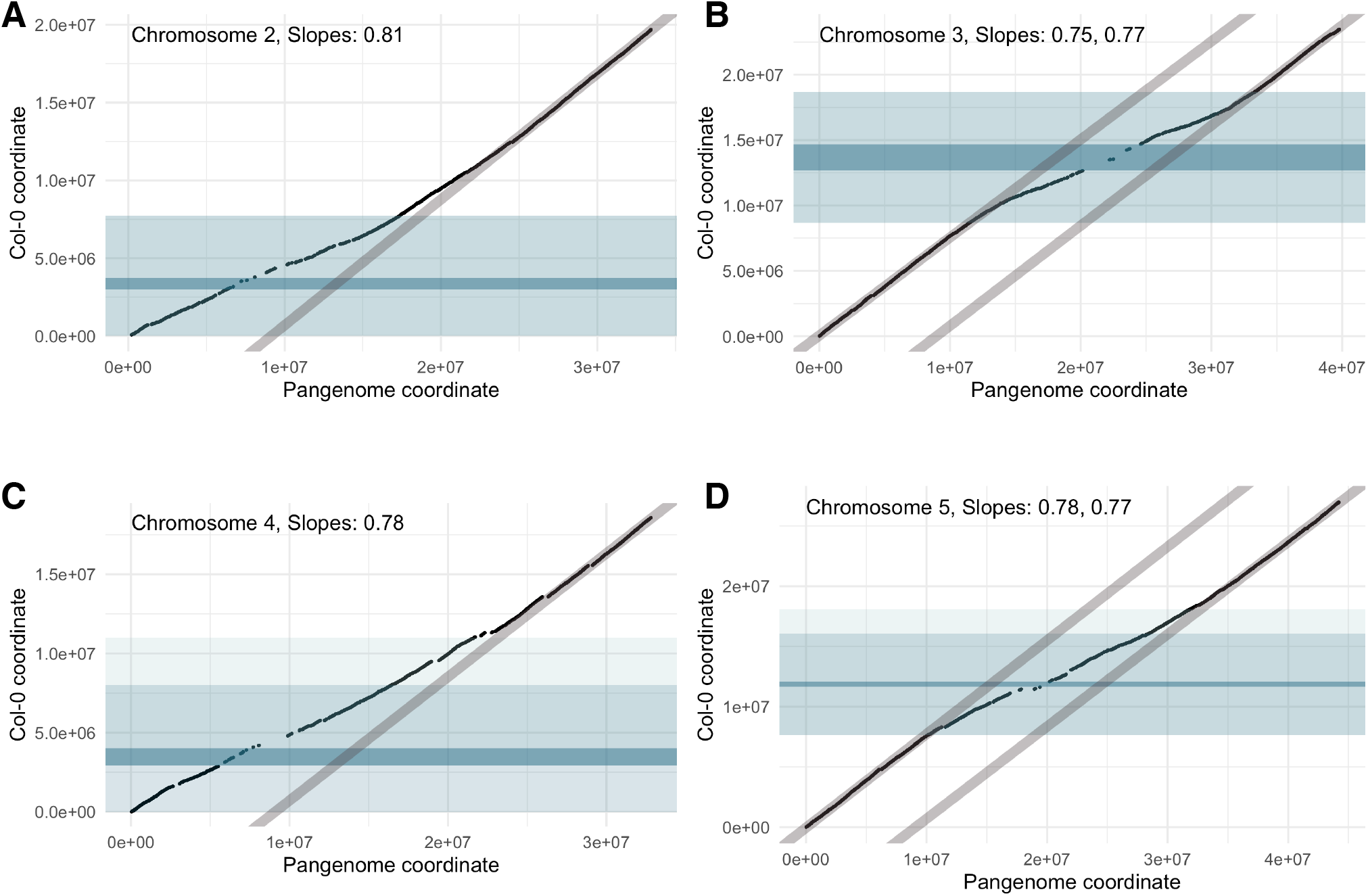
Pan-genome vs. reference genome coordinates for each chromosome. The pericentromeric region (light blue) shows a higher “dilution” of the spatial coordinates due to the increased number of SVs in this region. The centromeric region is dark blue.

**Extended Data Figure 33.**
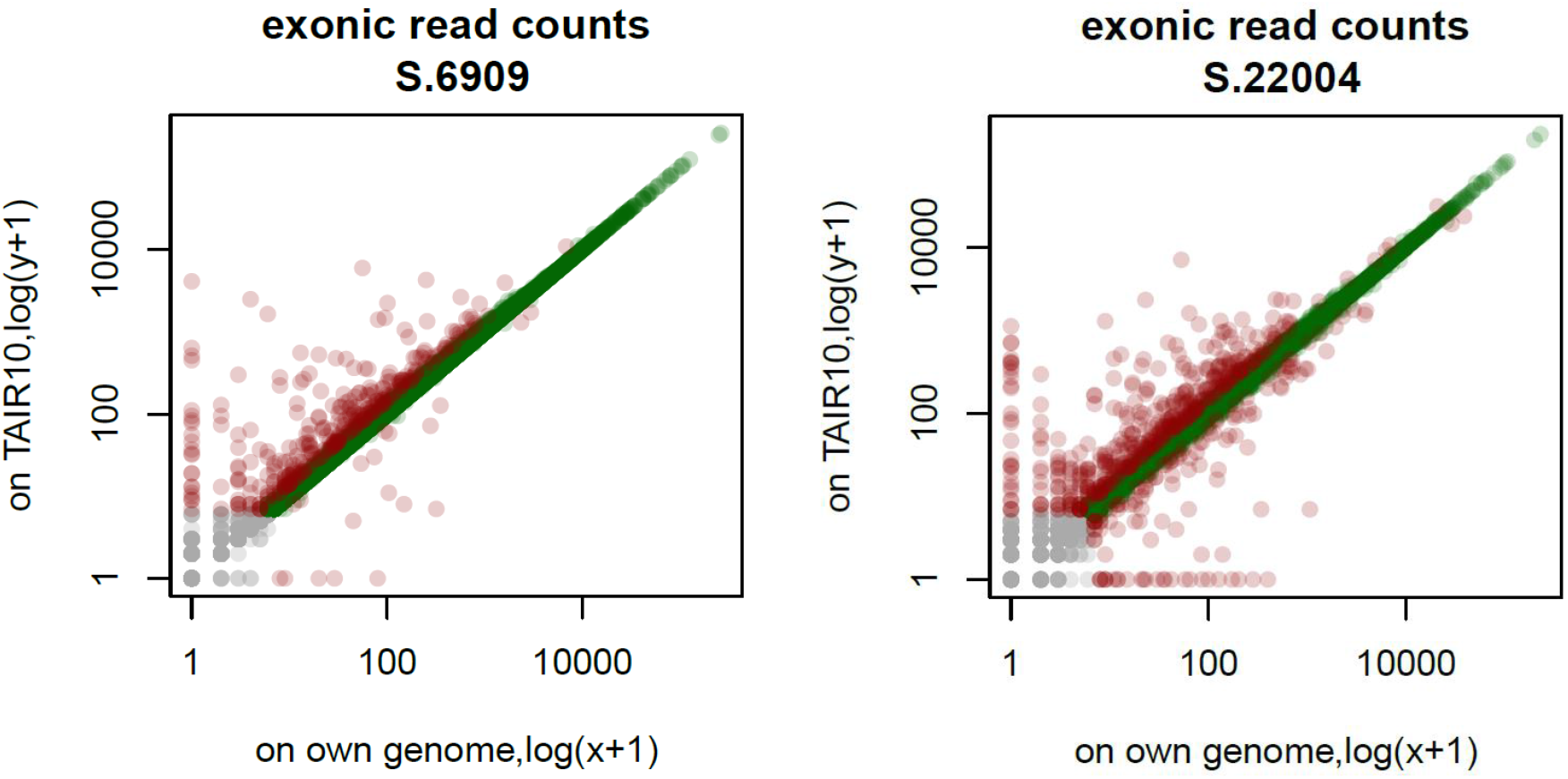
Reference bias when estimating gene expression. Raw counts in seedling RNA-seq samples for genes mapped to TAIR10 (using TAIR10 gene annotation) and to an accession’s own genome (using our annotation). Only one-to-one matched genes are shown.

## Supplementary information

- Supplementary Note – Detailed information about specific analyses, with figures.
- Supplementary Table 1 – Origin and sequencing data for the 27 accessions.
- Supplementary Table 2 – Large inversions.
- Supplementary Table 3 – Organellar inserts.
- Supplementary Table 4 – RNA-Seq read-map data.

## Supplementary Note

## 1 Genome assembly

### 1.1 Reciprocal translocation in 22001

**Supplementary Figure 1.**
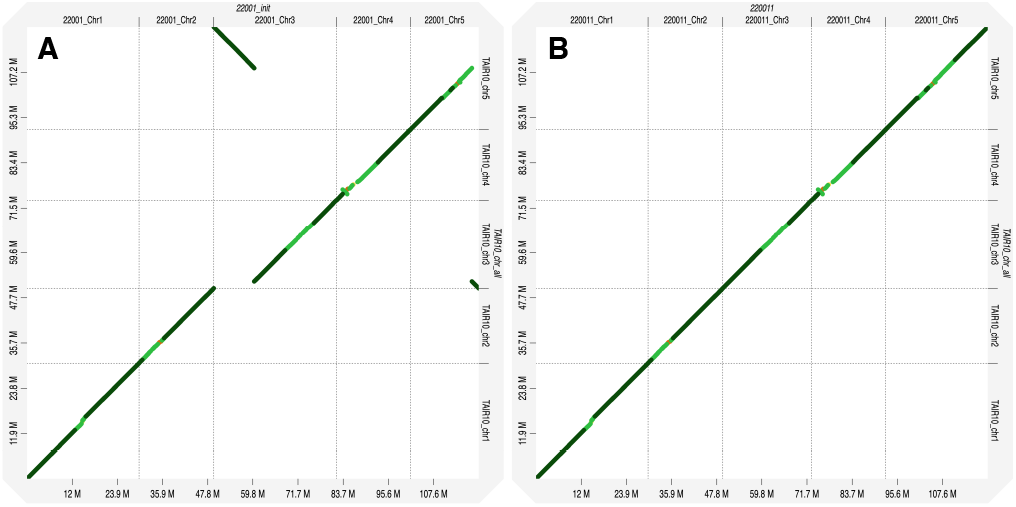
Reciprocal translocation in accession 22001. Dot plot of the original assembly 22001 (**A**) and of the modified assembly 22001m against (**B**) accession 22002. The translocation is readily seen at the beginning of chromosome 3 and the end of 5. Dot plots were created with D-GENIES.

We discovered a very large reciprocal translocation in accession 22001 (alternative name 85-3) from the Yangtze River region, which swapped the distal portions of chromosomes 3 and 5 (Supplementary Fig. 1). We validated the translocation by PCR with two sets of primer pairs designed to either amplify the standard arrangement of chromosomes 3 and 5 of Col-0, or the two translocation junction regions in accession 22001 (Supplementary Fig. 2). This rearrangement, which would presumably lead to decreased fertility in heterozygotes, appears to be quite rare as we did not identify other examples in a sample of 117 accessions sequenced with Illumina short reads from the same region ^35^. For the purposes of this study, we manually rearranged this genome to match the ancestral organization. To identify the exact breakpoints, we aligned chromosome 3 of 22001 to all other sequences of chromosome 3 with minimap2 (`-x asm5`). After filtering the alignments to retain those longer than 50 kb (`fpa drop -l 50000`), we removed the sequence from the start to the first position of alignment and added the reverse complement to the end of chromosome 5. A collection of the scripts can be found at the project GitHub repository.

Similar steps were followed for the segment in chromosome 5, using the sequence starting at the last position of the alignment to the end of the originally assembled chromosome. The sequence was removed from chromosome 5 and added to the beginning of chromosome 3.

We searched the repeat annotation for clues as to what type of sequence might have been responsible for the translocation, but found no obvious cause.

### 1.2 Genome size estimation

To estimate genome sizes from PCR-free reads, we employed a k-mer based approach ^98^ after pre-processing the datasets. First, we trimmed adapters from the raw reads and removed low quality sequences with cutadapt v2.4^116^ (`-q 20,15 -trim-n -minimum-length 75`). We aligned the trimmed reads to the organellar genomes of TAIR10 and the bacteriophage phiX174 genome with bwa-mem v0.7.17^119^, and executed a series of samtools v1.9^120^ commands to keep only reads for which both pairs did not align to any of these genomes. Briefly, we used `samtools view -b -f 12 -F 256` to obtain unmapped read pairs; `samtools view -b -f 4 -F 264` for paired-reads alignments in which read1 was unmapped and read2 was mapped; and `samtools view -b -f 8 -F 260` paired-read alignments in which read1 was mapped and read2 was unmapped. Then, we combined the three outputs of the described steps with `samtools merge`, discarded supplementary alignments with `samtools view -b -F 2048` and converted the BAM file to FASTQ format with `bedtools bamtofastq`. To avoid biases due to different read lengths for all subsequent analyses, we trimmed reads in all data sets to the common minimum of 124 bp and removed reads shorter than that with cutadapt v2.4^116^ (`--length 124 --minimum-length 124`). Finally, we counted 21 bp long k-mers with the commands `count -C -m 21 -s 5G` and `histo` from Jellyfish v2.3.0^121^, and the outputs were processed by the findGSE tool ^122^ to estimate genome sizes.

**Supplementary Figure 2.**
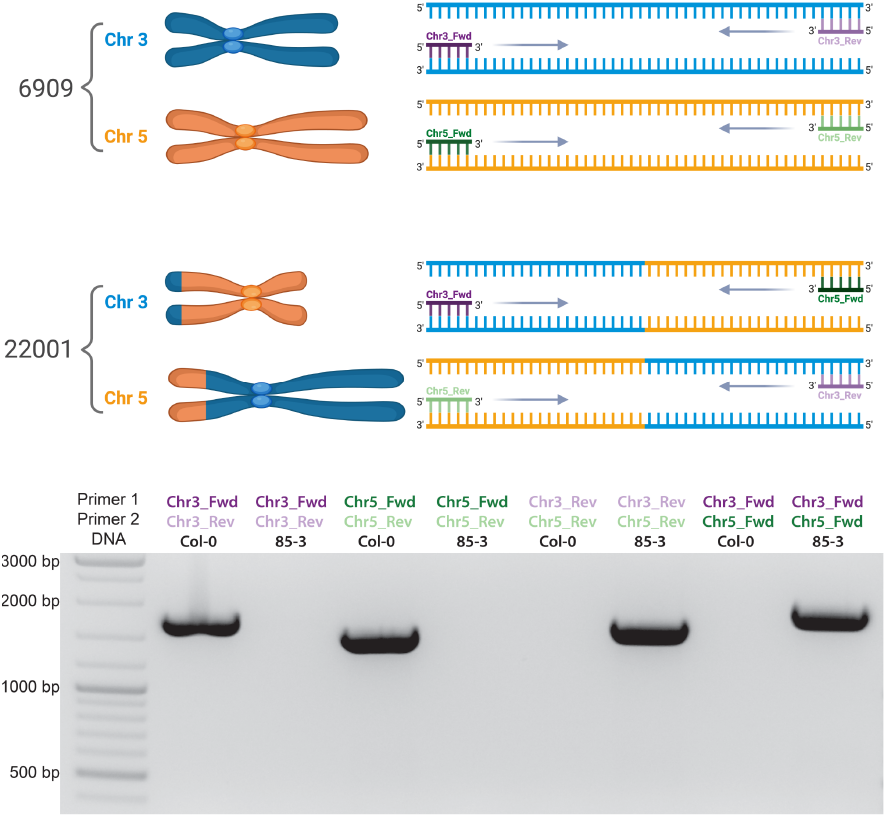
Validation by PCR of reciprocal translocation in accession 22001.

### 1.3 Estimation of satellite repeats

To estimate the contribution that the three main classes of satellite repeats make to the genomes — centromere, 45S and 5S rDNA repeats — from Illumina PCR-free reads, we used BLAST v2.2.29^123^ (`blastn -evalue 1e-10 -soft masking false -dust no -max hsps 1 -outfmt`), with reads trimmed to 124 bp in length as queries (see above) against a database with representative sequences of some of the most abundant or genetically diverse CEN159 and CEN178 satellite repeats collected from a recent pancentromere study ^16^, as well as three consensus 5S rDNA units ^89^, and a reference 45S rDNA copy ^90^. For each repeat class, the full-length of reads with a blast hit was added, and their sum was divided by the coverage estimated by `findGSE`. The sum of centromere and rDNA repeats estimated by this method and the difference between genome size (estimated by the k-mer approach) and the scaffolded assembly were highly correlated (*r*^2^ = 0.9639; *P* < 2.2 × 10^−16^; see Supplementary Fig. 3).

**Supplementary Figure 3.**
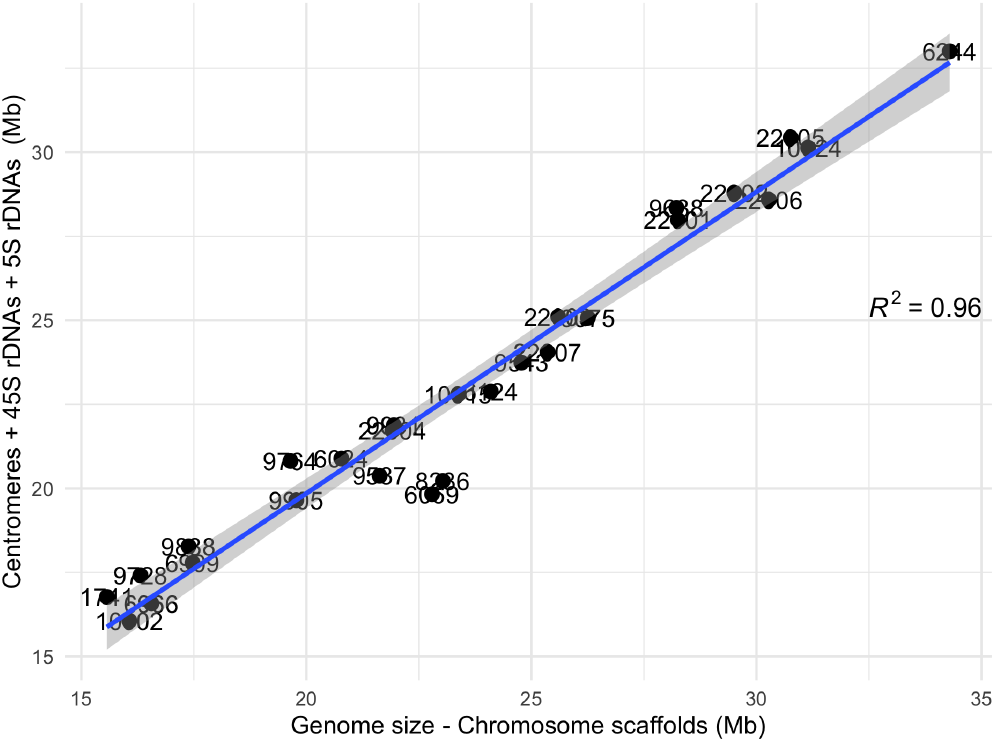
Correlation between the combined sum of centromere and rDNA repeats estimated by a BLAST-based approach with PCR-free reads, and the difference between the total genome size estimated by a k-mer approach with PCR-free reads and the summed lengths of the chromosome scaffolds.

## 2 Graph statistics

### 2.1 Deconstructing the graph

For variation detection we used the vg toolkit (v1.46.0 “Altamura”) ^103^. vg deconstruct was run on every path in the graph. Bubbles represent variation in graphs and are returned by the vg deconstruct subcommand in a modified VCF file. Bubbles have a start and end position (node based), and provide all different traversals from start to end position. This is reflected as a sequence of nodes and their direction and additionally with the covered base sequence. The resulting VCF files were merged based on start and end nodes of bubbles. The merged VCF file was further processed to only retain relevant information and converted to BED format for comparison with Pannagram results.

sSVs and cSVs in the graphs were defined as follows:

- SVs are representing indels, having one traversal that is very small (deletion) and a large one containing the SV sequence (insertion). Bubbles were identified as a sSV if the bubble was shared by all accessions in the graph (here 28), and as cSV if not. Traversals covering the insertion are at least 15 bp long and must be of high similarity (95% sequence). The deletion part of the bubble should be small, at most 5% of the length of the inserted sequence.
- Most cSVs correspond to bubbles that have a complex structure and/or are sub-bubbles of larger bubbles.

**Supplementary Figure 4.**
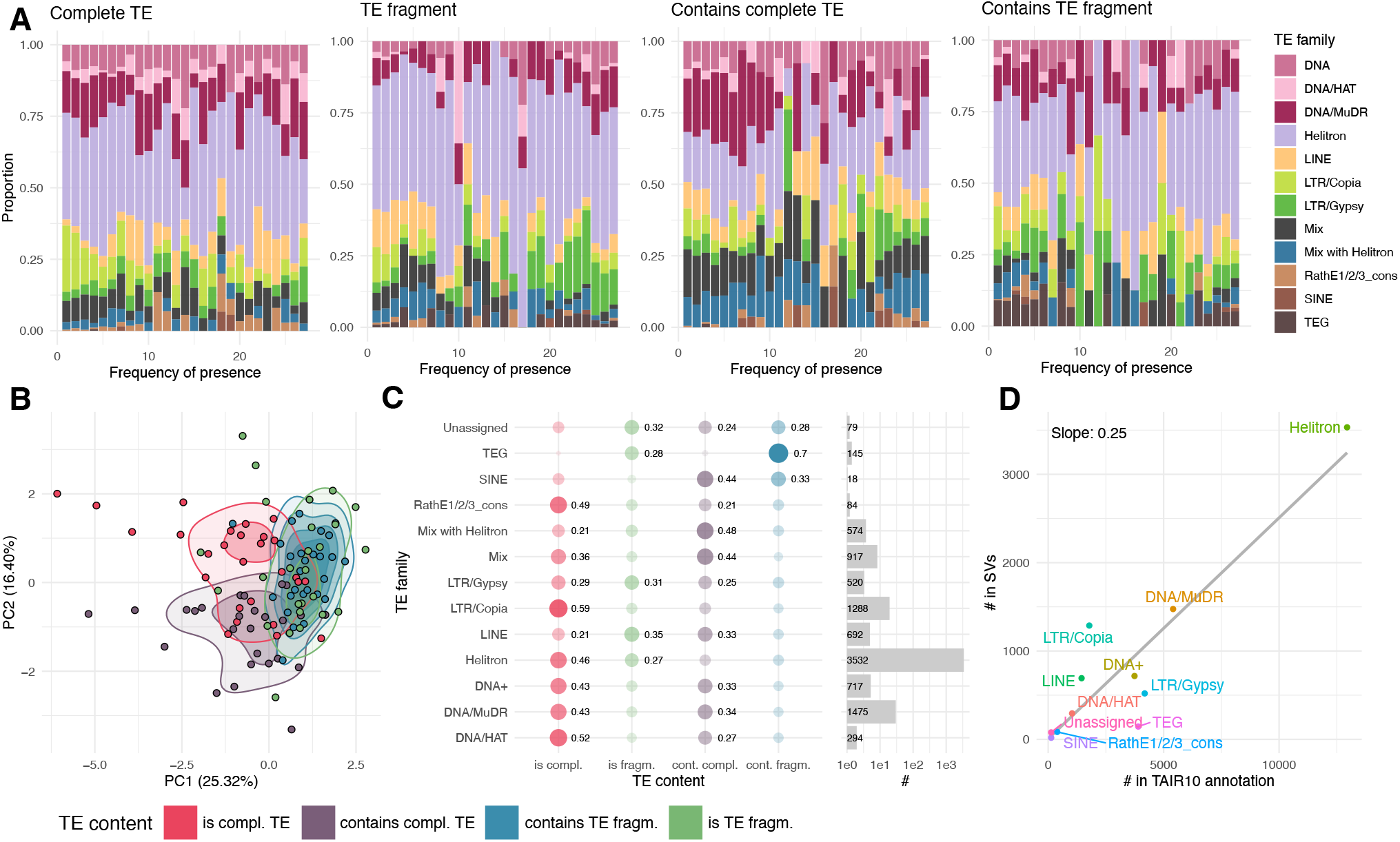
TE-superfamily content in SVs. **A**. Distribution of absolute numbers of sSVs across different TE content categories, based on the frequency of the presence allele. **B**. To confirm that that distributions in A differed, all columns in these subplots were taken as observations, and a PCA was performed. The second PC described the difference between categories of TE content based on their TE-superfamily content, in support of differences in TE-superfamily content in sSVs from different TE-overlap categories. **C**. Normalized distributions of TE-superfamily content according to the frequency of presence for each category of TE overlap. **D**. Correspondence between the number of TE superfamily members in the TAIR10 annotation and the representation of TE superfamilies in sSVs.

### 2.2 General pan-genome

To perform a reference-free pan-genome analysis, we utilized genome graphs built for each chromosome separately. The complete graph contains 18.3 million nodes and 20.9 million edges and has a total size of 225 Mb, with a mean compression rate over all chromosomes of 6.75%. Similar to other genome-wide analyses in this study, the large-scale reciprocal translocation in accession 22001 was masked to maximize linearity and increase resolution in the variation graph.

## 3 sSVs and TEs

We showed that most sSVs correspond to different categories of overlap with annotated TE sequences (Fig. 4). This analysis can broken down further to the level of TE superfamilies. Different TE superfamilies show very different patterns with respect to these overlap categories, presumably reflecting both the biology of the superfamilies and the quality of the annotation (Supplementary Fig. 4A). That there is a difference between the categories can also be seen in a Principal Components analysis, where the two first PCs of TE super-family composition distinguish four TE-content categories, consistent with different mechanisms underlying formation of different groups of TEs (Supplementary Fig. 4B).

We found several differences between TE superfamilies. For example, LTR/Copia elements appear to be both active and fairly well annotated, with roughly 50% of matches corresponding to presence-absence polymorphisms that include apparently complete elements, whereas matches to LINE elements rarely correspond to complete elements (Supplementary Fig. 4C). Also notable is the relationship between presence in sSVs and representation in the annotation. In general, they are strongly correlated (Supplementary Fig. 4D), but LTR/Copia elements, for example, seem over-represented in sSVs, consistent with their continuing to be active ^42^.

## 4 The gene-ome

### 4.1 Details about reconciling annotations

Our *de novo* annotation pipeline provided independent gene annotations for each accession. To find correspondences in annotated genes between accessions, we utilized the pangenome coordinate system. Mapping annotations to the common coordinates revealed that many genes had discordant annotations. The disagreements were of several types: (1) differences in exon-intron organization; (2) inconsistency in gene length; and (3) variability in the number of genes in a region. We resolved (3) by deciding to either merge or split genes based on a majority vote (Supplementary Fig. 5). We did not alter exon-intron organization and grouped genes into non-overlapping blocks for each strand to cover the maximum length (Supplementary Fig. 6). After identifying annotation groups on the pangenome coordinate, we compared the obtained gene and mRNA sequences between accessions. If, within a single annotation group, the sequences for accessions differed by more than 85% of their length or similarity, such genes were filtered out from the analysis. Such situations are associated with regions enriched with structural variations (SVs) or regions containing two or more haplotypes. Lastly, overleaping gene models on the pangenome coordinate system after the above-mentioned procedure were excluded from the analyses, ensuring that only one annotation group was considered. The resultant annotation groups were considered as genes.

After generating a consistent annotation, we projected it back onto all accessions. If a locus was present in an accession but not annotated by the *de novo* annotation, we still identified it as a gene, though without the exon-intron model. This procedure helped us to avoid underestimating the number of segregated genes (Supplementary Fig. 7). Supplementary Fig. 8 illustrates the final consensus annotation.

### 4.2 Genes and TEs

*Arabidopsis thaliana* annotations group were compared with TAIR10 annotations for pseudogenes and TEs based on their position in the pan-genome coordinate systems. Annotation groups without overlap to TAIR10 were defined as ‘new gene’. We then separated genes into two main categories: Protein-coding genes (TAIR10 and new) and TE genes (TAIR10 and new, plus all TE-like genes). New genes with more than 50% overlap to TEs in TAIR10 were classified as TE. To further characterize these ‘new’ genes, their coding sequence were compared using DIAMOND’s blastp module against UniProt DB (version 2024 06). The best hit in UniProt that shared at least 50% sequence identity over at least 50% of the *A. thaliana* gene length was further used. The functional and taxonomical annotation of UniProt hits were retrieved from uniprot KB using uniprot id-mapping service. Annotation groups were classified as TE if their uniprot annotation met any of the following conditions: uniprot protein names including transposon, retrotransposon, transposase, transposable, reverse transcriptase; GO terms including transposase, DNA transposition; their UniProt domain annotation includeing transposase, transposon, reverse transcriptase; had Pfam domains PF14223, PF03078, PF03732. Additional functional categories were defined based on protein name, keywords, domains, GO terms, and subcellular location annotations associated with the UniProt hit. We further categorized ‘new’ genes as ‘TAIR10 high similarity’ if they shared more than 80% sequence identity over 80% of the coding sequence length and as ‘TAIR10 medium similarity’ if they share between 45% and 80% sequence identity with the TAIR10 hit (see further details in Supplementary Note 4.3).

**Supplementary Figure 5.**
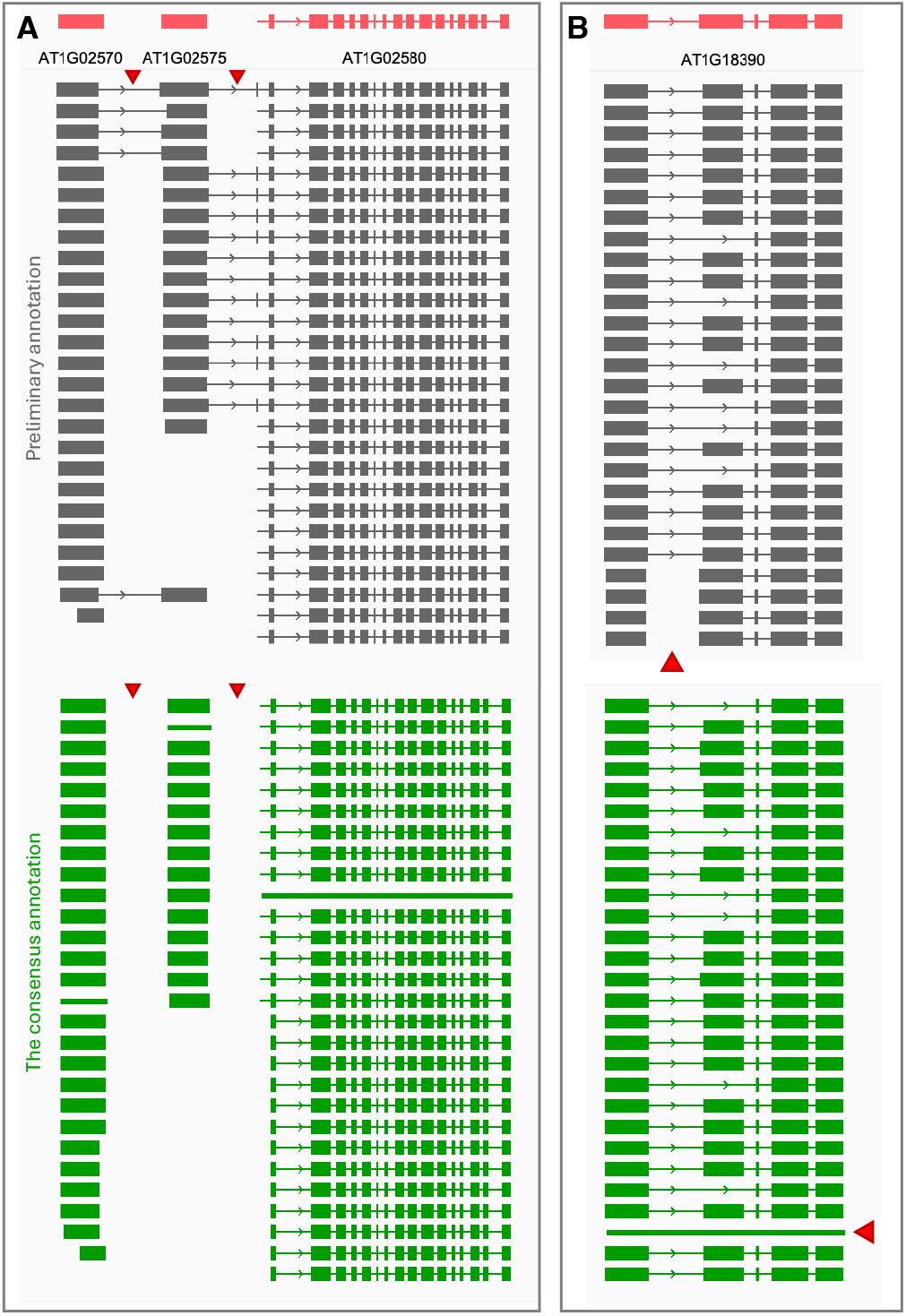
Reconciling ambiguous gene annotation with majority voting. **A**. Situation where the solution was to split. Red triangles denote regions to solve. **B**. Situation where the solution was to merge. Red triangles denote the region to solve and the resultant merged gene.

**Supplementary Figure 6.**
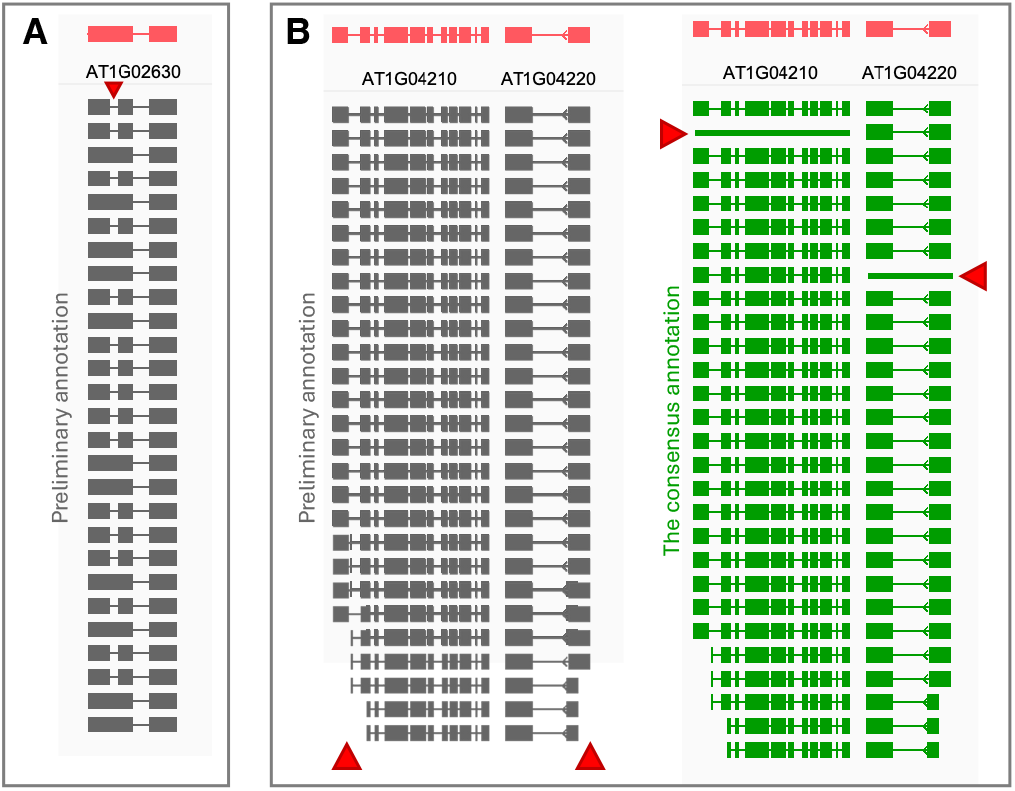
Variation in gene models. **A**. Variation in exon-intron structure. Red triangles denote regions of exonintron variation in the *de novo* annotations between accessions. **B**. Variation in gene length. Problems and how they were solved are indicated with red triangles. Red triangles on the gray plot indicate regions where transcript lengths vary between accessions. Red triangles on the green plot indicate regions of genes in the final annotation that cover the positions of all transcripts.

**Supplementary Figure 7.**
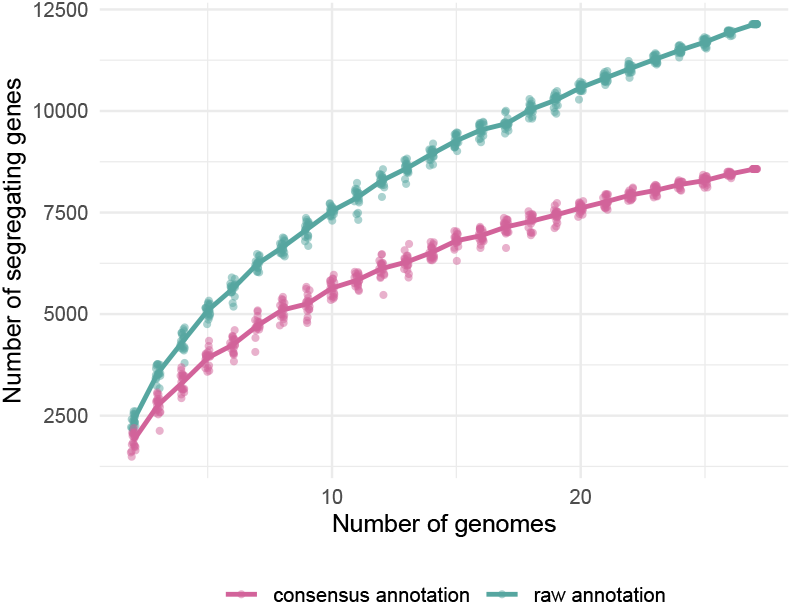
Number of segregating genes. Comparison of the estimates based on the raw *de novo* annotation and on the final consensus gene annotation.

**Supplementary Figure 8.**
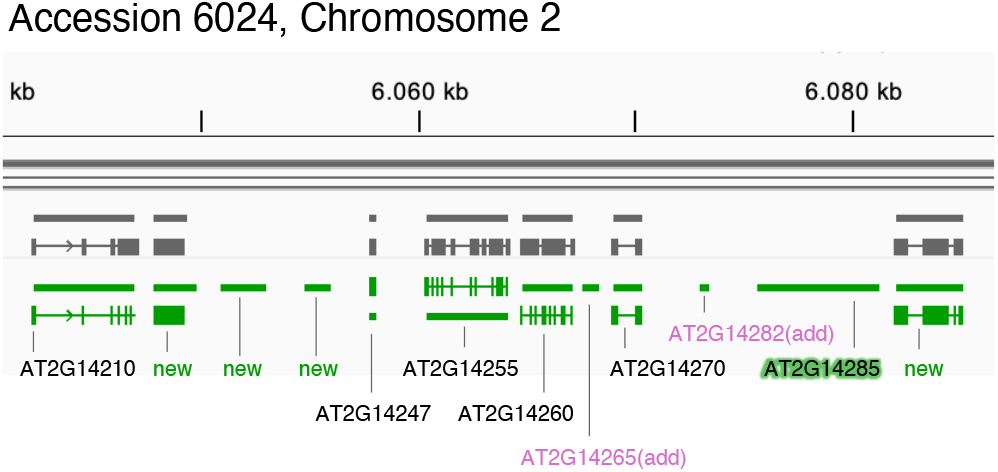
An example of the final consensus annotation. This region of accession 6024 on chromosome 2 shows different sources of annotation: (i) TAIR10 genes, which were found in this accession (black names), (ii) TAIR10 genes, which were found in other accessions (black names with a green glow), (iii) TAIR10 genes, which were not found in any accession (purple names), (iv) new genes, which were found in this accession (denoted as “new” with the gene model) and also in other accessions (denoted as “new” without the gene model, a line only).

### 4.3 New genes

We performed a more detailed analysis of “new genes”— those that did not have a match in the TAIR10 gene annotation—to investigate their origin and characteristics. First, we assessed the chromosomal distribution of new genes, which are enriched around the centromeres (Supplementary Fig. 9A), while TAIR10 annotated genes are depleted around centromeres. Grouping new genes based on their TE-sequence content shows that the pericentromeric enrichment is largely due to genes with high TE content (Supplementary Fig. 9B). Thus we find new protein-coding (non-TE) genes all across the chromosomes with slight depletion around centromeres. We preliminary classified genes into TE and non-TE based on their TE-sequence content: genes with >50% TE-sequence in at least one accession were classified as TE, the rest as non-TE (protein-coding) (Supplementary Fig. 9C). Unsurprisingly, newly annotated genes were strongly enriched for TE genes, compared to TAIR10-annotated genes.

To make sure that we did not miss any additional TE genes, we further looked for amino acid similarity of the genes in our annotation to genes encoding proteins known to participate in TE function, such as transposase (Methods) and added them to genes classified as TE (Extended Data Fig. 25A).

Next, we compared the presence-frequency of old (TAIR10) and new genes (Supplementary Fig. 10A) and found that, as expected, the great majority (94.2%) of TAIR10-annotated genes are fixed in the population, while the majority of new genes are not (82.9%). While new TE genes showed similar frequency distribution to new protein-coding genes, genes that were classified as TAIR10 TE genes are, surprisingly, often fixed (52.2%), which likely is due to the skewed representation of TAIR10 TE genes in our annotation: we only identified a small portion of annotated TE genes (565), and since our annotation pipeline relied on RNA expression, those are likely TE genes with higher-than-average expression. Since expression is markedly higher among fixed genes (Fig. 6I), this might explain an over-representation of fixed “TAIR10 TE genes” genes in our annotation. Finally, high-frequency genes are more likely to have already been already annotated in TAIR10.

In order to identify the origin of the “new” genes in our annotation, we searched for these genes in the genomes of TAIR10, other accessions, *A. lyrata*, other Brassicaceae species, and all other species using UniProt KB (Supplementary Fig. 10B). 28% of previously unannotated protein-coding genes had a very high sequence similarity (at least 80% identity) to TAIR10, indicating that they originated by recent gene duplication. 25% had medium similarity (45-80% identity) suggesting more ancient gene duplications, or, alternatively, a recently duplicated gene modified by a large structural variation. Another 31% did not have a sequence match in TAIR10 but their sequences could be found in other accessions, or in other Brassicaceae species, including *A. lyrata*. Consistent with a likely origin via duplication, new genes were strongly enriched for being present in 2 or more copies (6.0-fold enrichment over TAIR10 genes, Supplementary Fig. 10C), with TAIR10-high-similarity genes showing the highest – 10.0-fold – enrichment. We can assume that a few of the duplications responsible for the creation of the new genes are tandem, as new genes showed a 3.0-fold enrichment in being in a tandem copy in at least one accession, compared to TAIR10-annotated genes, with high-similarity genes showing 6.5-fold enrichment (Supplementary Fig. 10D). We next investigated why the newly annotated genes were missing from the TAIR10 annotation. For the majority (75.9%) of the “new” protein-coding genes, their genetic sequence is simply absent from the Col-0 genome (Supplementary Fig. 10E). New genes might also be missing from the TAIR10 annotation because they are not expressed in Col-0, but we were able to annotate them if they are expressed in other accessions, which was the case for 7.5% of new genes; About 45% of the new genes that are present also in Col-0 (the TAIR10 accession) are also expressed there—albeit at a low level. It is also worth noting that our annotation pipeline identified many new genes that are not expressed in any accession, *i*.*e*., their identification had been solely due to sequence-based prediction (see Methods). From the categories of new genes, the TAIR10-high-similarity genes have the highest rate of the locus being absent from TAIR10 (Supplementary Fig. 10E, bottom), confirming our intuition that these are the most recently arising genes that stem from a structural change of inserting a new gene locus through gene duplication.

**Supplementary Figure 9.**
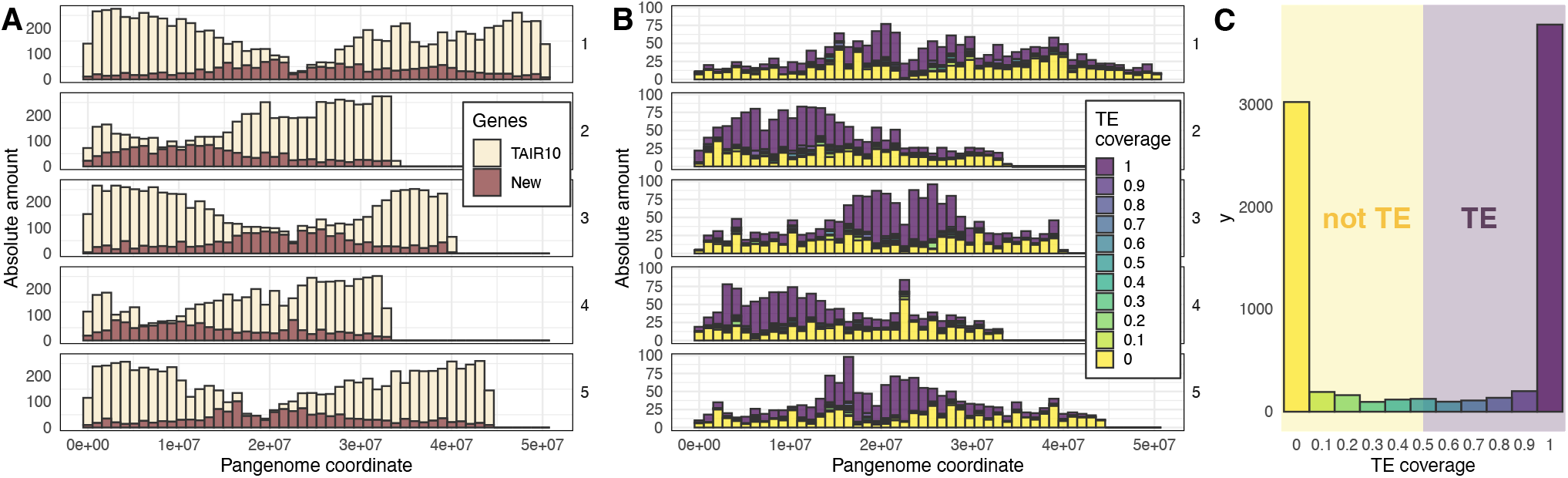
TEs in *de novo* annotation. **A**. “New genes” without correspondence to TAIR10 genes cluster around the centromere. **B**. The distribution of “new genes” along the chromosome is explained by gene models with strong similarity to annotated TEs. **C**. The distribution of TE similarity among these gene models is bimodal, with 59% matching TEs. **D**. A high proportion is (unsurprisingly) not found among annotated TAIR10 genes.

**Supplementary Figure 10.**
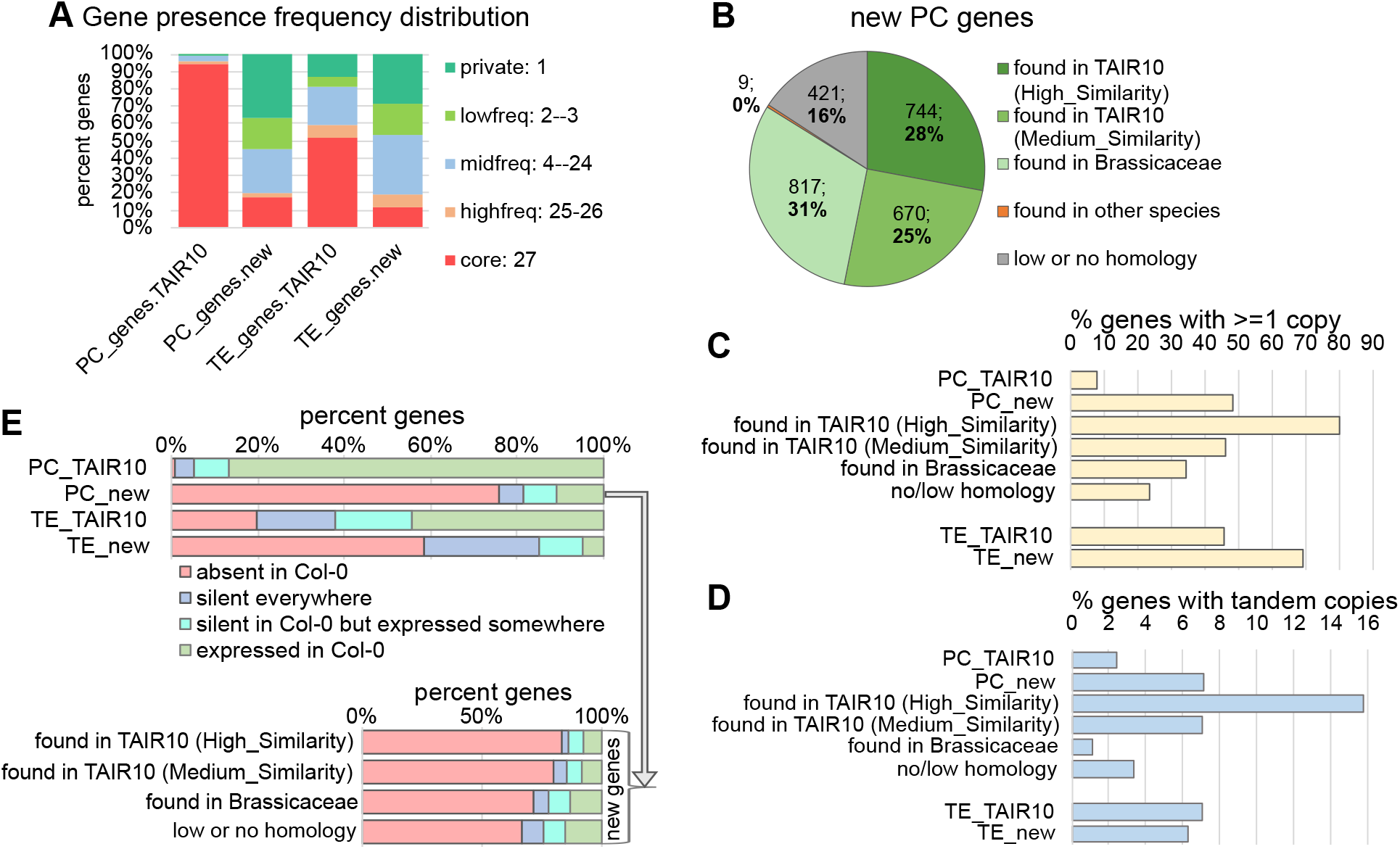
“New” genes. **A**. Presence-frequency for newly-annotated and TAIR10-annotated protein-coding and TE genes. **B**. New protein-coding genes grouped by amino acid sequence similarity to TAIR10 genes and UniProt genes best hit. High and medium similarity to TAIR10 gene corresponds to hits having more than 90% or 45% sequence identity covering more than 80 or 50% of the new PC, respectively. Hits to UniProt genes were considered if they share more than 45% sequence identity covering more than 50% of the new PC. For “New PC” found only in UniProt, the groups were defined based on UniProt taxonomic annotation. **C**. Percent of genes in different categories that are duplicated in at least one accession. **D**. Percent of genes in different categories that are tandemly duplicated in at least one accession. Tandem duplications were defined as copies located within 10kb from each other. Copy search was done using the simsearch tool in the Pannagram package. **E**. Investigating reasons for why a new gene may or may not be in the TAIR10 annotation: the locus is absent from Col-0 (red); the gene is silent in all accessions and tissues (blue); the gene is silent in Col-0, but expressed in at least one other accession (turquoise); the gene is expressed in Col-0 (green). An expression cutoff was used: locus-wide TPM > 0.25. Top: TAIR10- and newly annotated protein-coding and TE genes. Bottom: groups defined in B.

We have shown that many new genes are often physically absent from specific accessions, but even where present, the new genes are often not expressed and show signs of PRC2 and TE-like silencing (Supplementary Fig. 11A). New TE genes show very high levels of silencing, consistent with the idea that many old (fixed) (Supplementary Fig. 10A-D) TE genes manage to increase their frequency by becoming harmless and thus requiring less silencing than new and potentially more deleterious TE genes. This is also supported by our finding that new TE genes are strikingly more enriched in functional TE domains (Fig. 6G). Different categories of new protein-coding genes show no significant differences in expression or silencing, with the exception for slightly lower H3K9me2 and CG methylation level on TAIR10-high-similarity genes (Supplementary Fig. 10E).

## 5 Errors and biases in SNP-calling

### 5.1 Results

To explore the sources of errors in SNP calling, we took advantage of the high-coverage, PCR-free Illumina short-read data that were used to correct PacBio reads during assembly. Briefly, for a pair of accessions, we first identified SNPs using a whole-genome alignment, then designated one of the genomes as a reference and called SNPs using short reads from the other genome. The process was repeated until each genome had been used as reference genome for every other genome in the sample. The results were compared using the whole-genome alignments as “ground truth”.

Using standard parameters, we found that we could call far more SNPs than in our previous work (over 80% of those found in whole-genome alignment), but that the False Discovery Rate,

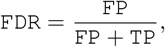

was very high, ∼ 7%. A comprehensive investigation into the extent to which it is possible to decrease the number of false positives without increasing false negatives awaits investigation, but a closer look at the nature of these SNP-calling errors was already informative. As illustrated in Fig. 8A:

- 17.3% of SNPs identified in the whole-genome alignment were entirely missed by short read-based SNP-calling because they lie in regions not covered by reads because of mapping problems. This is the main reason SNP-calling underestimates polymorphism.
- 18.4% of SNPs called using short reads were pseudo-heterozygous (because the material analyzed was highly inbred and true heterozygous sites should not exist), almost entirely because the sample contained duplicated regions not found in the reference genome, leading to erroneous mapping of reads to regions that do not correspond to their true origin.
- 83.1% of false positive SNP calls were due to spurious read-mapping caused by various forms of polymorphism (not only SVs). A minor fraction (13.9%) were technical artifacts due to our two pipelines making different choices about local alignment.
- *Bona fide* false negative SNP-calls were either caused by read-mapping or local alignment problems, in roughly equal proportions.
- Other types of errors (including base-calling and random coverage) made trivial contributions.

It goes without saying that the precise numbers of SNP-calling errors will depend on the parameters used, but the qualitative conclusions will hold.

**Supplementary Figure 11.**
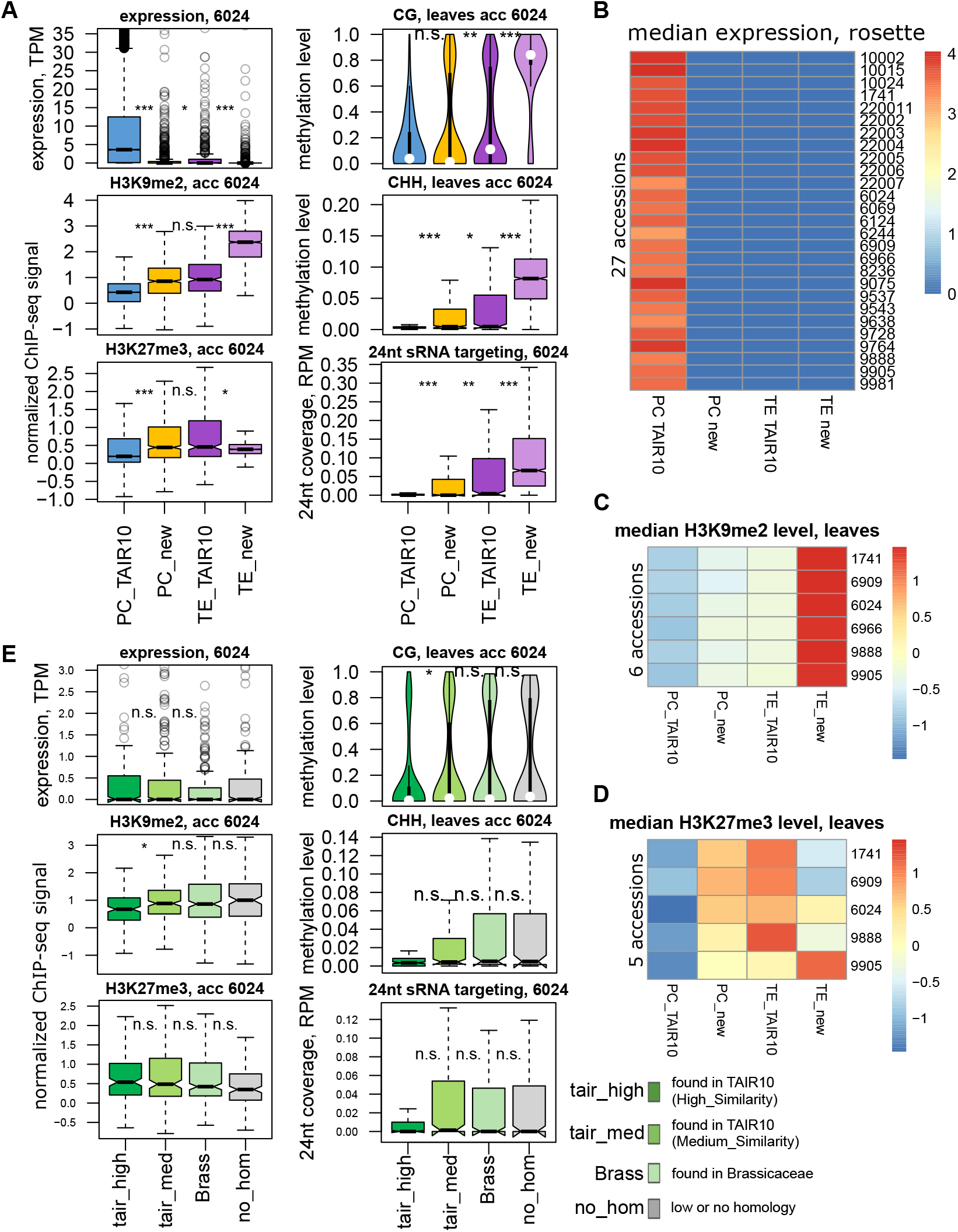
“New” genes silencing. **A**. Locus-wide RNA expression in 9-leaf rosette, the levels of H3K9me2 and H3K27me3 in mature leaves, the levels of CG and CHH methylation in mature leaves and 24nt sRNA level in flowers ^63^ for TAIR10 and new genes. Data for accession 6024 is plotted; other accessions showed similar patterns. **B**. Median expression of TAIR10 and new genes in 9-leaf rosette in 27 accessions. **C-D**. Median H3K9me2 level in 6 accessions and median H3K27me3 level in 5 accessions for TAIR10 and new genes. **E**. Locus-wide RNA expression in 9-leaf rosette, the levels of H3K9me2 and H3K27me3 in mature leaves, the levels of CG and CHH methylation in mature leaves and 24nt sRNA level in flowers ^63^ for the 4 categories of new protein-coding genes defined in (Supplementary Fig. 10B) Data for accession 6024 is plotted; other accessions showed similar patterns.Significance estimates from Mann-Whitney tests (^***^: *P* < 10^−10, **^: *P* < 10^−5, *^: *P* < 10^−2^, n.s.: *P* ≥ 10^−2^).

**Supplementary Figure 12.**
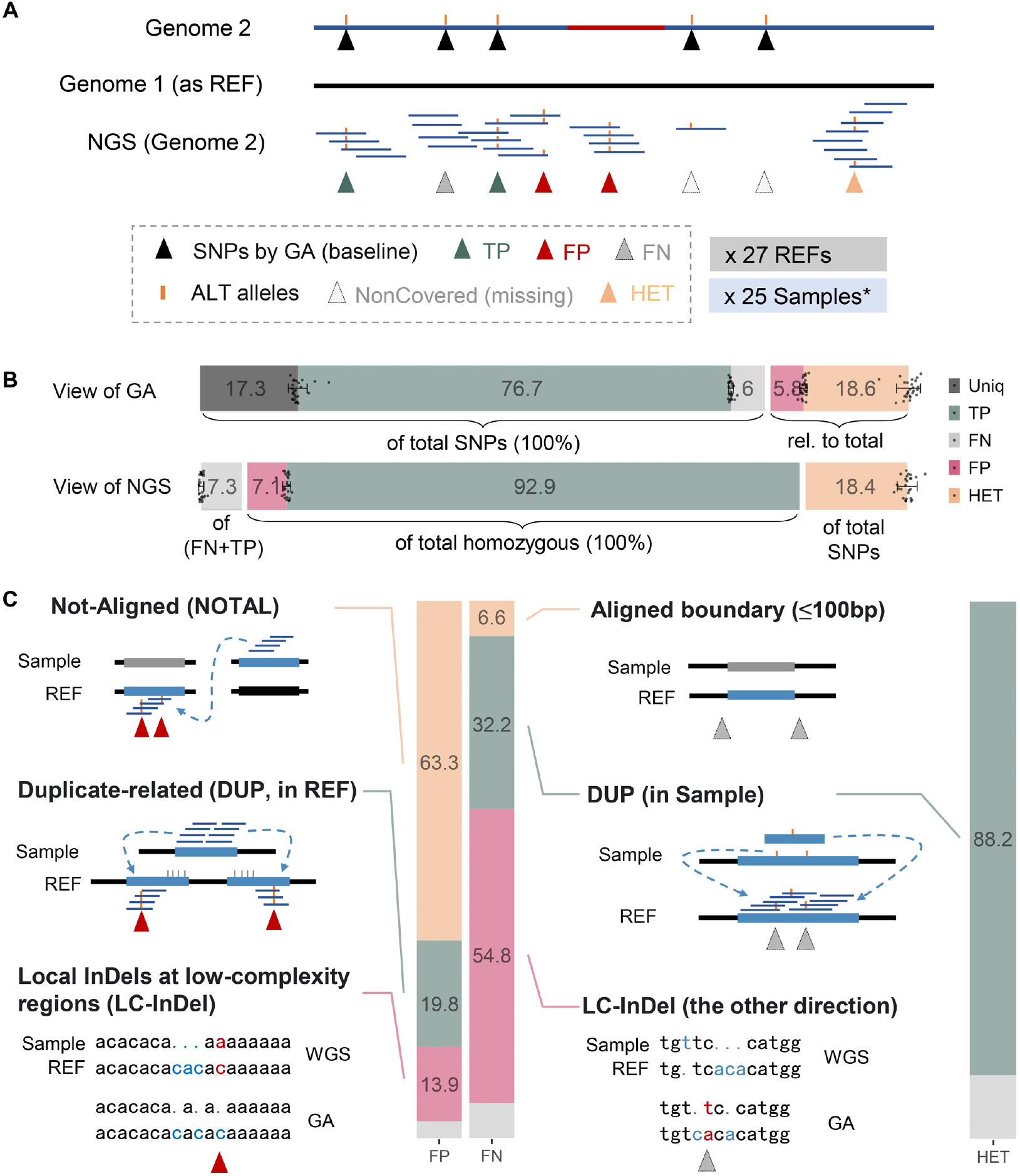
Sources of errors in SNP-calling. **A**. Overview of the approach. Genome 1 is used as reference, and SNPs are identified both by whole-genome alignment to Genome 2 and by SNP-calling using Illumina reads from Genome 2. The SNPs identified in the whole-genome alignment treated when compared with Illumina read-based SNPs. The process was repeated using all 27 genomes as reference, and using 25 NGS read sets with sufficient coverage as samples. **B**. Sources of errors, in percent, for whole-genome alignment SNPs and Illumina read-based SNPs (*cf*. Fig. 8A). Of the whole-genome alignment SNPs, 76.7% (on average) were also called using Illumina reads, while 6% were FNs, and 17.3% were not called because the region was not covered by any Illumina read. Relative to the number of whole-genome alignment SNPs, 18.6% of Illumina read-based SNP calls were heterozygous and 5.8% FPs. From the point-of-view of the Illumina read-based SNPs, 7.1% of homozygous SNP calls were FPs. Heterozygous calls constituted 18.4% of total SNP calls, and the false negative rate, FNR = FN*/*(FN + TP) was 7.3%. **C**. Diagram of the sources of the three types of errors (FP, FN, and HET; see text for details, and Supplementary Figs. 13-16) for concrete examples).

**Supplementary Figure 13.**
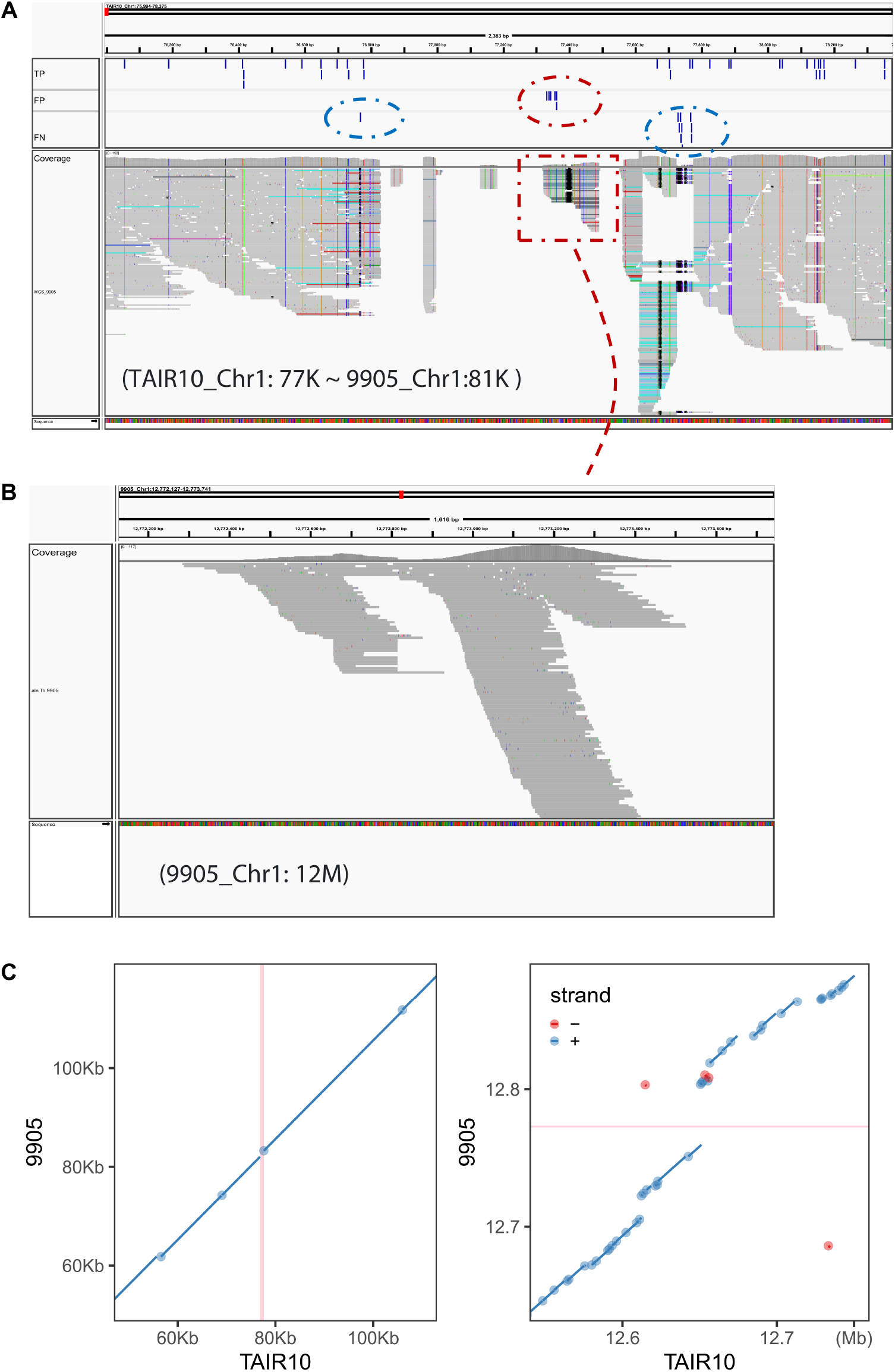
Not Aligned (NOTAL) regions can cause both FP and FN calls. FP calls can be found in NOTAL region, while FN can be found in flanking sequence. **A**. Screen shot from IGV showing FP (red circle) and FN (blue circle) SNPs, and the corresponding read-mapping, using WGS from 9905 to TAIR10 as an example. **B**. Aligning the mis-mapped reads (red square in A) to 9905 instead results in a well-mapped read to another non-collinear region. **C**. Genome alignment showing that both the regions of (A) and were NOTAL, while the vertical red lines represent the mis-mapped (non-collinear, gap) region at TAIR10, from which the reads can be well-mapped to the region in 9905 (horizontal line). The diagonal lines are aligned regions and the gaps are NOTAL regions.

**Supplementary Figure 14.**
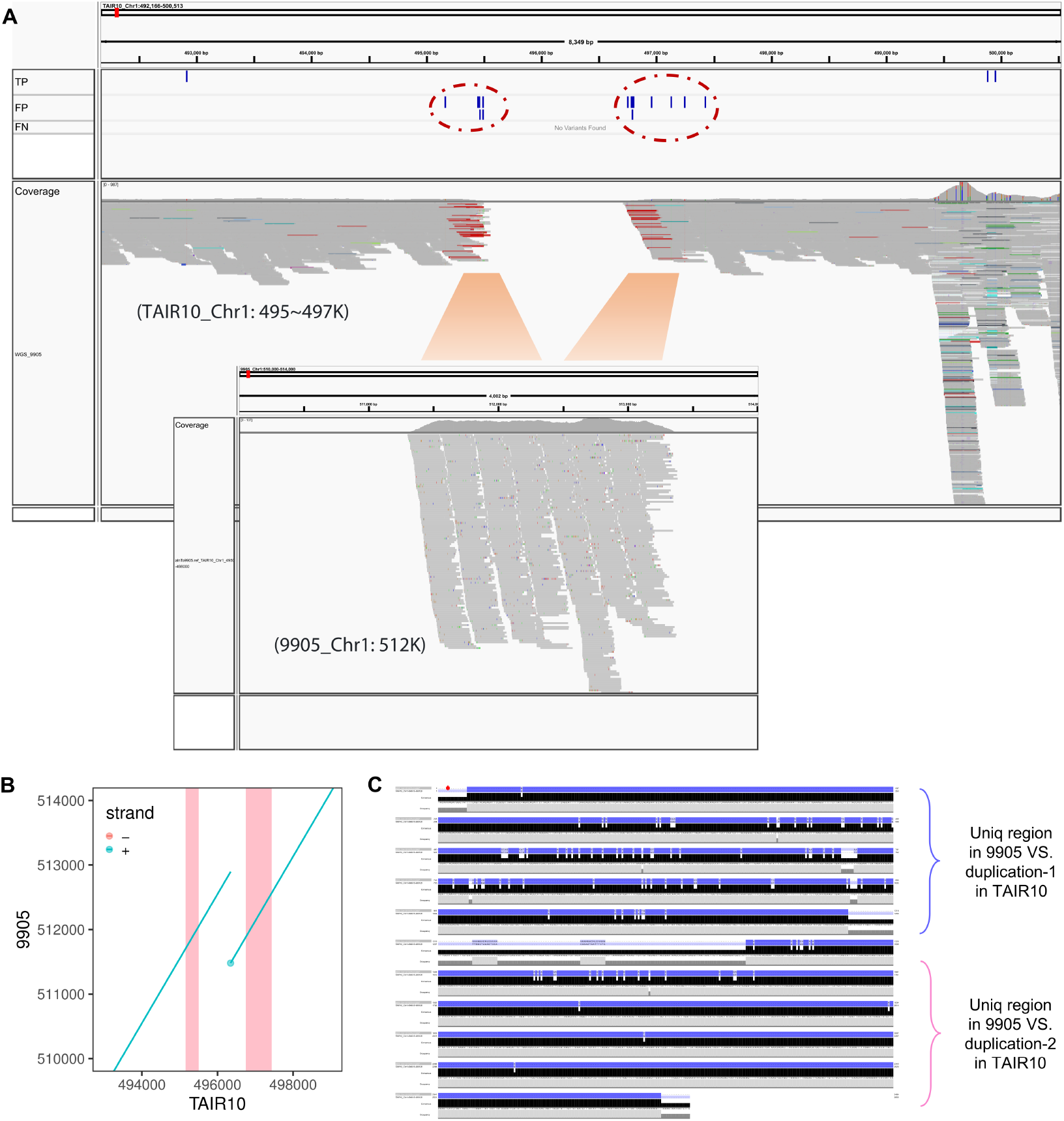
How duplications introduce FP calls. FP calls can occur when there were more duplications in reference than in the new genome. **A**. Screen shot from Integrated Genome Viewer (IGV) shows the FP (framed by red circle) SNPs and corresponding reads mapped situations. The bottom IGV screen shot shows how these reads were mapped to the corresponding genome (9905), suggesting that the reads from one region were mapped separately to two different regions with a large gap between them. **B**. Further investigation showed that the region has only one copy in 9905 and two tandem duplications in TAIR10, resulting in a duplication event called in WGA, but not SNPs. The red regions indicate the two FP clusters based on TAIR10. **C**. This shows why the reads from one region of the 9905 genome were mapped separately to the early part of the first copy and the late part of the second copy of TAIR10. This is because these two regions were more identical to the one copy of 9905, so the reads could still be well aligned elsewhere in the TAIR10 genome, but not multi-mapped, resulting in false SNP calls.

**Supplementary Figure 15.**
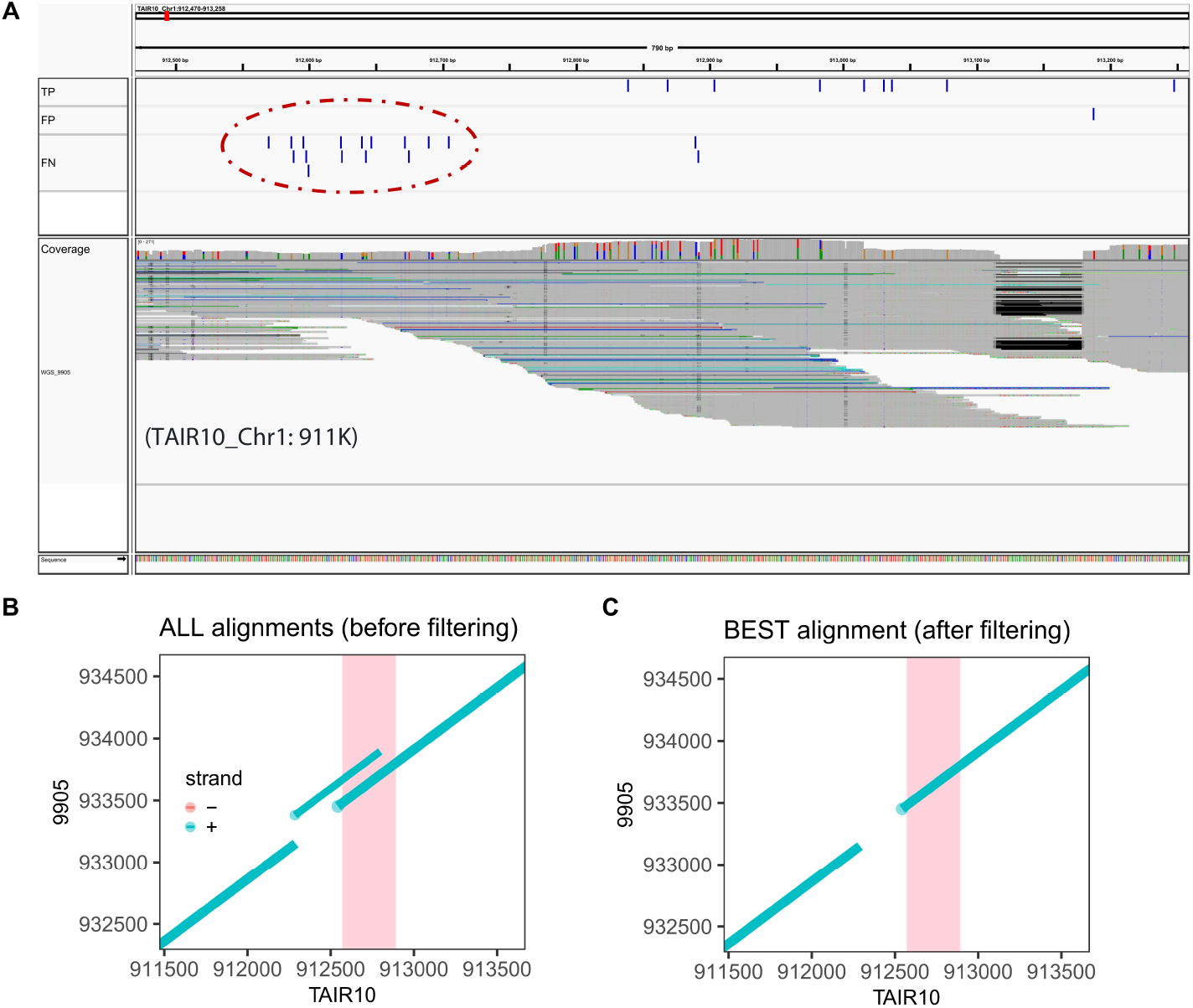
How duplications introduce FN calls. More duplications in the new genome than in the reference can lead to FN SNPs. **A**. Screen shot from Integrated Genome Viewer (IGV) shows the FN SNPs (framed by the red circle) that can only be called in WGA and not in short-read mapping. **B**. Genome alignment showed that this region was duplicated in another short fragment in its own genome. The red region indicates the FN cluster based on mapping to TAIR10. **C**. However, in WGA analysis, it is easy to remove the short non-collinear alignment and use only the collinear alignment to call SNPs, whereas in NGS, this is impossible due to multi-mapping, and real differences are missed.

### 5.2 Methods and parameters

We used pairwise whole-genome alignments as baseline (“ground truth”) for comparison with Illumina read-based SNP calls. Mummer4^124^ was employed for whole-genome alignment between corresponding chromosomes of accession pairs, and the `delta-filter` (`-1 -q -r`) was used to eliminate one-to-many and many-to-many redundant matches, followed by `show-snps` (`-THCr`) to extract variants. Only SNPs were retained for further examination.

For Illumina read-based SNP-calling, we used the PCR-free Illumina reads generated for this study. Not only were these data generated from the same DNA preparations as was used for the PacBio CLR reads, but the coverage was also in general far higher than in previous work ^34^ (one accession, 22002, had less than 10x coverage, and was not used). Reads were mapped to each of the 27 genomes with BWA-MEM v0.7.17^119^, followed by use of Picard tools to remove duplicates, and GATK HaplotypeCaller v4.3^125^ to call variants with `gVCF` mode. Each of the 27 “reference” genomes was used to call SNPs in the remaining 25 samples. Variants were filtered by `QD < 2.0, FS > 60.0, MQ < 40.0, SOR > 4.0`, and genotypes with low quality or coverage were changed to missing with `GQ < 20 or DP < 3`. The VCFs were separated into pairwise comparisons for each combination of investigated accession and “reference”. Since SNPs could be nested with other types variants (for example, with REF and ALT alleles of GTT and TTT,G, respectively), multi-allelic loci were first converted to bi-allelic with `bcftools norm -m -any` and the alternate alleles were realigned against the reference allele (transforming GTT/TTT to G/T) using vcfwave ^126^. Heterozygous SNPs were extracted, and homozygous SNPs (MNPs here were considered as multiple SNPs) were retained for further evaluation.

The gVCF files were used to measure the genome fraction covered by Illumina reads. Regions covered by fewer than 3 reads, or with GQ below 20, together with all heterozygous sites, were all considered not missing and excluded from the analysis (Fig. 8A). Illumina read-based FPs were SNPs called with Illumina reads but not from the whole-genome alignment or SNPs found with both approaches but with different ALT alleles. Illumina read-based FNs were SNPs only called with a whole-genome alignment at sites covered by Illumina reads.

To investigate the sources of SNP errors (Supplementary Fig. 12), the following steps were taken:

**Supplementary Figure 16.**
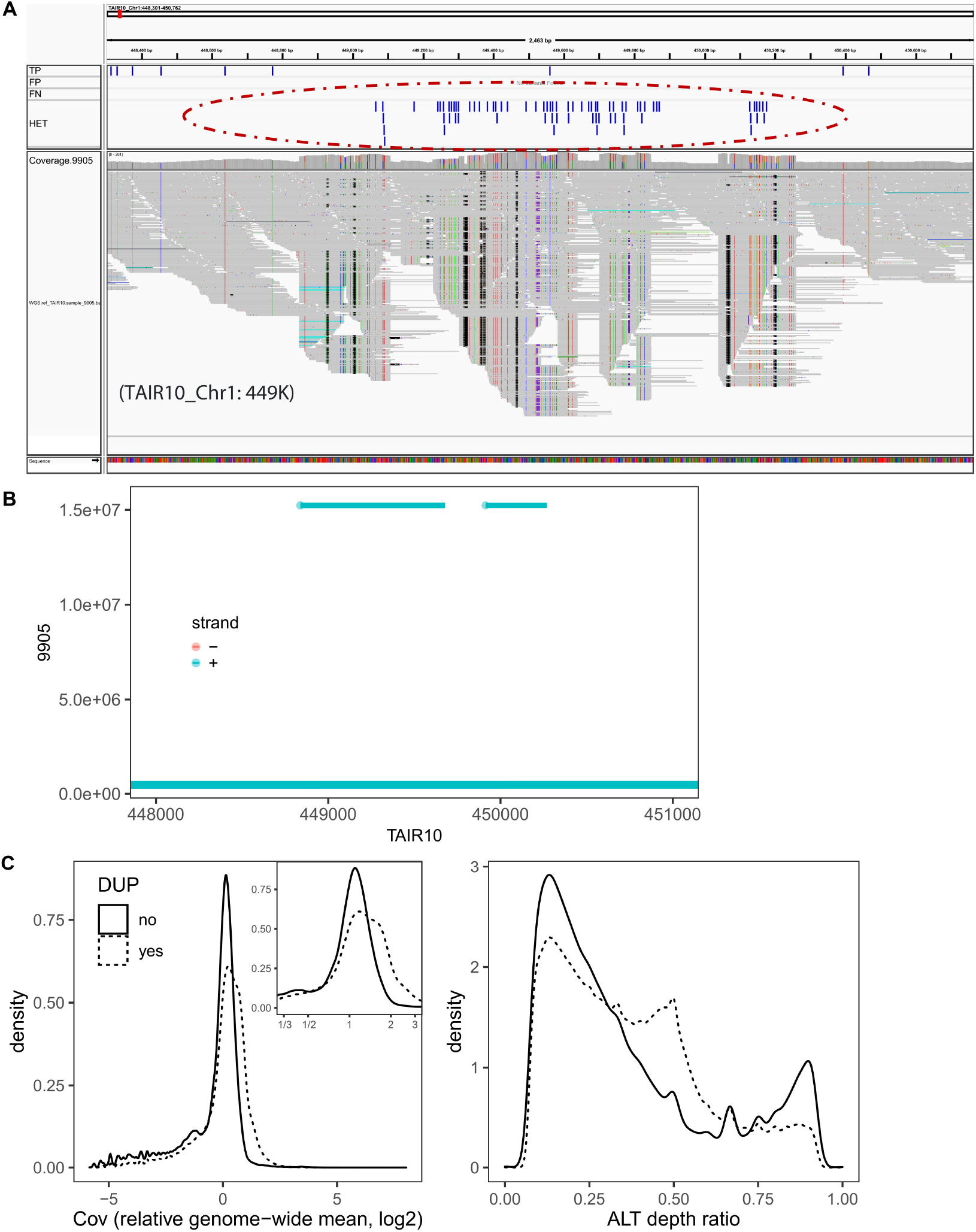
Duplications contribute to the majority of mis-called heterozygous SNPs. Heterozygous (HET) SNPs can be called when there are more duplications in the new genome than in the reference. **A**. Screen shot from Integrated Genome Viewer (IGV) shows that heterozygous SNPs (framed by the red circle) were called in WGS due to more duplications in the sample genome than in the reference. The reads from the duplications were mapped to one location on TAIR10, leading to the identification of HET SNPs. **B**. Genome alignment showed that this large region is a segmental duplication in the new genome. **C**. The HET alleles from duplications that could be identified (DUP=yes, 88.2% of all HET variants) had higher read coverage, with the ALT allele depth ratio (reads with ALT alleles/total reads covered) enriched at 50% than the rest (that the 11.8% of HET variants that could not identify evidence of duplications).

1. The filtered (`-1 -q -r`) “delta” file produced by Nucmer was used to determine the ALN (Aligned, marks a block of aligned sequence between two genomes) and NOTAL (Not Aligned, highlights sections of a genome that did not align with the other) regions, and FP SNPs located within NOTAL regions were considered as “FP NOTAL”, and FN SNPs located within ALN regions but within 100 bp of NOTAL boundaries were considered “FN: Aligned boundary”.
2. The raw delta file including one-to-many and many-to-many alignments was processed using the command `show-snps` of Mummer4, and SNPs covered with multiple alignments (`[R]>0`) were retained to overlap with the FP calls to obtain the “FP: DUP” category. To estimate the fraction of FNs caused by this category, whole-genome alignments (including inter-chromosomes) were produced with Nucmer, followed by `show-coords -THrcl` to obtain aligned corresponding regions.
3. The indels called from the WGA were used to investigate how much of the remaining NGS FP SNPs could be explained, while the indels called from the PCR-free Illumina reads were used to assess that of FN SNPs. Erroneous (both FP and FN) SNPs `within 5 bp` of indels identified by the alternate technology were considered to be caused by local indels. These predominantly occurred in low complexity regions.

## 6 Errors and biases in DNA methylation profiling

Many cytosines can be potentially missed and some of their cytosine contexts (CG/CHG/CHH) are incorrect, because of genetic differences between the TAIR10 reference and a focal genome (Supplementary Fig. 17A).

Even though for those that could be well aligned and with the same cytosine context, a DMR between accessions identified based on TAIR10 can be a FP (no significant differences based on corresponding genomes). To investigate this effect, BS-seq reads of each accession were mapped to TAIR10 (REF) and to the accession-specific genomes (OWN), and the unique and de-duplicated alignments were used to summarize the context-dependent methylation. Differentially methylated regions (DMRs) between two accessions were identified based on mapping to REF and mapping to OWN, respectively. For each comparison, say Acc1 vs Acc2, four analyses were performed including Acc1 REF, Acc2 REF, Acc1 OWN and Acc2 OWN. Mappings from accession-specific analyses (*i*.*e*. Acc1 OWN and Acc2 OWN here) were aligned to the TAIR10 genome, and only aligned regions with a 1-to-1 correspondence were considered further. Only regions covered by at least 3 reads in both REF (either Acc1 REF or Acc2 REF) and OWN (either Acc1 OWN or Acc2 OWN) were retained, and DMRs between two accessions were identified based on REF (Acc1 REF vs Acc2 REF, DMR REF) and OWN genomes (Acc1 OWN and Acc2 OWN, DMR OWN). Cytosines were grouped into 100 bp non-overlapping windows, and a fast Fisher’s exact test (https://github.com/al2na/methylKit/issues/96) was used to identify DMRs (*P* ≤ 0.01) by summing all reads supporting methylation of all cytosines (allC) and total coverage in a given window. The analyses were repeated for individual cytosine contexts (CG, CHG, and CHH) (Supplementary Fig. 17B). Windows with zero read coverage (Cov = 0) in one accession, but over three methylated reads (mC ≥ 3) in the other, were manually assigned as DMRs (*P* = 1*E* − 4). The same analysis was applied to both REF and OWN analysis, and for various contexts: all Cs (allC), only CG, CHG, and CHH.

To determine the enrichment overlapping with annotations of potential False Discovery DMRs (FDR DMR) from REF-based analysis, continuous FDR DMRs were first grouped together, and the obtained intervals in REF were used to measure the overlap with annotations. A permutation overlap analysis (same interval size and number in each chromosome were simulated for each genome comparison, with 100 repeats; implemented with `bedtools shuffle` was considered as background. The median fold change (in log2, 0 overlap was assigned to 0.2 for log calculation) of the observed annotation overlapping the 100 permutations was obtained as an indication of enrichment of each annotation in every genome comparison, and the enrichment of each annotation in all 10 sample comparisons (vs. the other one) were measured as a general trend (Supplementary Fig. 17C).

**Supplementary Figure 17.**
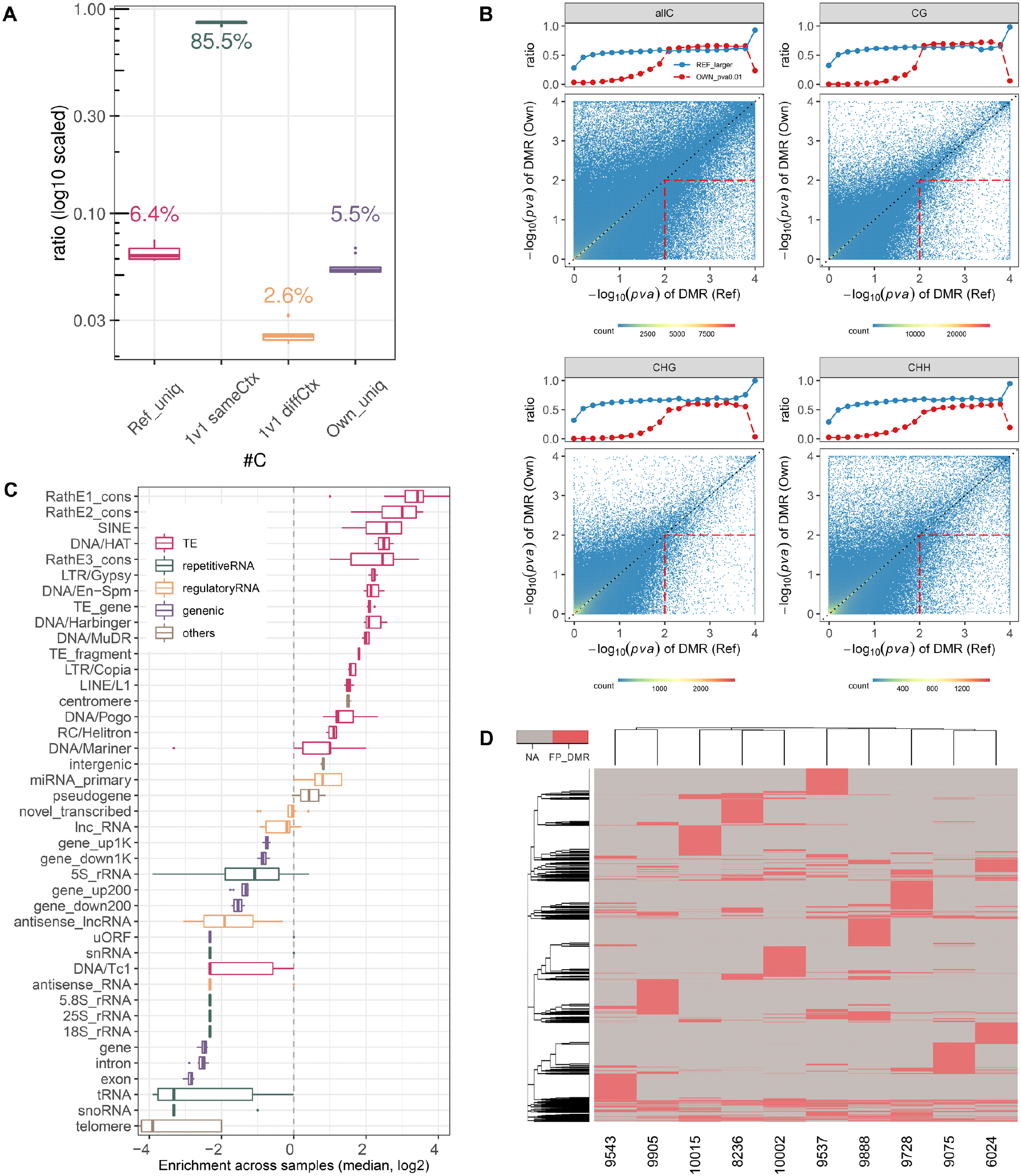
Reference bias: methylation. **A**. Differences in cytosine content between TAIR10 and genomes newly assembled in this study due to sequence variation: “ uniq” indicates unaligned sequences from insertions, deletions, duplications, etc.; “1v1 sameCtx” represents aligned cytosines with conserved CG/CHG/CHH context; and “1v1 diffCtx” have aligned cytosines in a different context. Values are median across 11 samples. **B**. Differentially methylated regions (DMRs) between accessions were identified in 100-bp windows by mapping BS-seq reads to TAIR10 (x-axis) or to an accessions’s own genome (y-axis). The red squares are FPs identified only when using the TARIR10 reference genome. Plots show results for comparison of all accessions to accession 6966. The blue line above shows the fraction of DMRs at a given p-value threshold (along the x-axis) with higher p-values in the TAIR10 reference analysis. The red line shows the fraction of reference-based DMRs that are also significant at *P* < 0.01 in the own-genome analysis. **C**. FP DMRs (*P* < 0.01 only in the reference-based analysis) are strongly enriched for TEs based on the annotations from the TAIR10. **D**. Most of the identified FP DMRs are unique to a specific accession.

